# HUWE1 Amplifies Ubiquitin Modifications to Broadly Stimulate Clearance of Proteins and Aggregates

**DOI:** 10.1101/2023.05.30.542866

**Authors:** Mengying Zhou, Rui Fang, Louis Colson, Katherine A. Donovan, Moritz Hunkeler, Yuyu Song, Can Zhang, Siyi Chen, Dong-hoon Lee, Gary A. Bradshaw, Robyn Eisert, Yihong Ye, Marian Kalocsay, Alfred Goldberg, Eric S. Fischer, Ying Lu

## Abstract

Selective breakdown of proteins and aggregates is crucial for maintaining normal cellular activities and is involved in the pathogenesis of diverse diseases. How the cell recognizes and tags these targets in different structural states for degradation by the proteasome and autophagy pathways has not been well understood. Here, we discovered that a HECT-family ubiquitin ligase HUWE1 is broadly required for the efficient degradation of soluble factors and for the clearance of protein aggregates/condensates. Underlying this capacity of HUWE1 is a novel Ubiquitin-Directed ubiquitin Ligase (UDL) activity which recognizes both soluble substrates and aggregates that carry a high density of ubiquitin chains and rapidly expand the ubiquitin modifications on these targets. Ubiquitin signal amplification by HUWE1 recruits the ubiquitin-dependent segregase p97/VCP to process these targets for subsequent degradation or clearance. HUWE1 controls the cytotoxicity of protein aggregates, mediates Targeted Protein Degradation and regulates cell-cycle transitions with its UDL activity.

## INTRODUCTION

The protein degradation system is critical for maintaining protein homeostasis and a healthy cellular environment by selectively targeting both soluble factors and aggregates in eukaryotic cells. Misregulation of protein homeostasis perturbs a broad range of cellular activities and may lead to accumulation of protein aggregates that are a hallmark of multiple human pathologies, such as neurodegeneration, cardiomyopathy and type-II diabetes (*1*).

The decision to degrade a protein target is largely encoded in its ubiquitylation status which is recognized by both the proteasome and autophagy pathways(*2, 3*). Insoluble targets, such as pathological protein aggregates, are typically enriched with ubiquitin(Ub) or Ub chains that signal the degradation machineries for their clearance(*4, 5*). Increasing evidence suggests that the Ub-dependent p97/VCP segregase/unfoldase complexes, which are coupled with both the proteasome and autophagy pathways, interact with various cellular aggregates/condensates that include stress granules, aggresomes and certain inclusion bodies, and promote their dissolution(*6–13*). The aggregate-processing capacity of p97/VCP has been implicated in the pathogenesis of neurodegeneration and p97-associated diseases(*14*). It is, however, still poorly understood how the ubiquitylation pathways recognize these aggregates and attach appropriate Ub patterns that recruit the p97 segregase to promote aggregate clearance.

Besides autophagy, the proteasome is also involved in dissolving a variety of protein aggregates/condensates(*13, 15–19*). Although the canonical process of proteasomal degradation, in which a substrate is first ubiquitylated by a substrate-specific Ub ligase (E3), and then engages with the 26S proteasome for proteolysis, has been well documented(*20*), recent studies suggest that degradation of some important substrates may involve additional steps and factors. For example, the proteasome-associated Ub ligase Hul5/UBE3C is required for a complete degradation of substrates containing structurally-stable domains(*21, 22*); the p97/VCP complex extracts substrates from cellular structures and mediates the proteasomal degradation of important therapeutic targets(*11, 23, 24*). However, not all degradations require the p97, indicating that additional factors or steps may be involved in controlling the degradation process. The identity of these factors and more broadly how they may change the canonical degradation process to advance its capacity on difficult targets remains obscure.

In a search for factors that stimulate protein degradation, we identified HUWE1 which possesses a novel type of Ub ligase activity that we call Ubiquitin-Directed ubiquitin Ligase, or UDL, and discovered that this activity targets HUWE1 to both soluble factors and protein aggregates and promotes their degradation or clearance. HUWE1 (HECT, UBA, and WWE domain containing protein 1; also known as Mule/ARF-BP1/LASU1) is a 482-kDa HECT-domain Ub ligase that is evolutionarily conserved in eukaryotes, is present in most cell types and is genetically associated with several neurological disorders, such as intellectual disability and autism (*25–27*). HUWE1’s UDL activity recognizes the local density of Ub chains on targets and rapidly expands the Ub modifications to promote both proteasomal degradation and p97-mediated unfolding of targets. We further find that the UDL activity of HUWE1 is critical for the activity of several Targeted Protein Degradation (TPD) drugs and regulates cell division through promoting the timely degradation of cell-cycle regulators. Furthermore, HUWE1 binds and amplifies the Ub signal on protein aggregates to stimulate their clearance via the p97/VCP complex, critical for mitigating the cytotoxicity of these aggregates. We discuss the nature of HUWE1’s mode of action and explain how it is used versatilely in important regulatory processes.

## RESULTS

### HUWE1 conjugates additional Ub chains on ubiquitylated substrates

To discover activators of protein degradation, we synthesized model substrates with structural features that are refractory to processing by the proteasome. In one design, we conjugated a variant Ub (Ub^L73F^), which is partially resistant to deubiquitylation(*28*), to the N-terminal degron domain of cyclinB (cycB) fused with an easy-to-unfold fluorescent protein cpGFP at the C-terminus (cycB-cpGFP- polyUb^L73F^)(*29*), using purified Anaphase Promoting Complex (APC) paired with the E2 UbcH10 (Fig. S1A)(*30, 31*). Unlike wildtype (WT) Ub-conjugated cycB-cpGFP which was degraded efficiently by the purified 26S proteasome, cycB-cpGFP-polyUb^L73F^ was poorly degraded (Fig. S1B). Since substrate deubiquitylation is a prerequisite for efficient proteasomal degradation(*32*), this slower degradation is likely due to defects in Ub^L73F^ removal by proteasome. Interestingly, degradation of both WT Ub and Ub^L73F^-conjugated cycB proceeded efficiently in a crude extract of unsynchronized HeLa S3 cells (Fig. S1B), which required the proteasome activity (Fig. S2A,B). We confirmed that the L73F mutation did not affect the efficiency of APC-mediated polyubiquitylation (Fig. S1A), nor did it affect substrate-proteasome interaction in a single-molecule assay(Fig. S3)(*33, 34*). These results indicate the presence of a non-proteasomal factor in extract that stimulates the degradation of the cycB model substrate.

Western blot analysis of cycB-cpGFP-polyUb^L73F^ showed a rapid wave of further ubiquitylation upon incubation with the cell extract, while the overall Ub distribution in extract remained stable (Fig. 1A). The cycB species that carried extra Ubs disappeared gradually during incubation, but could be stabilized by a proteasome inhibitor. Consistently, an inhibitor of the Ub E1 enzyme (MLN7243) largely stabilized the fluorescent signal of cycB-cpGFP-polyUb^L73F^ in extract (Fig. S2A,B). The stimulation of degradation in extract is not dependent on the cpGFP domain, as the degradation of wtUb-conjugated cycB-EGFP or cycB-mNeonGreen was also enhanced in Hela extract in an E1-dependent fashion (Fig. S2C).

**Figure 1.**
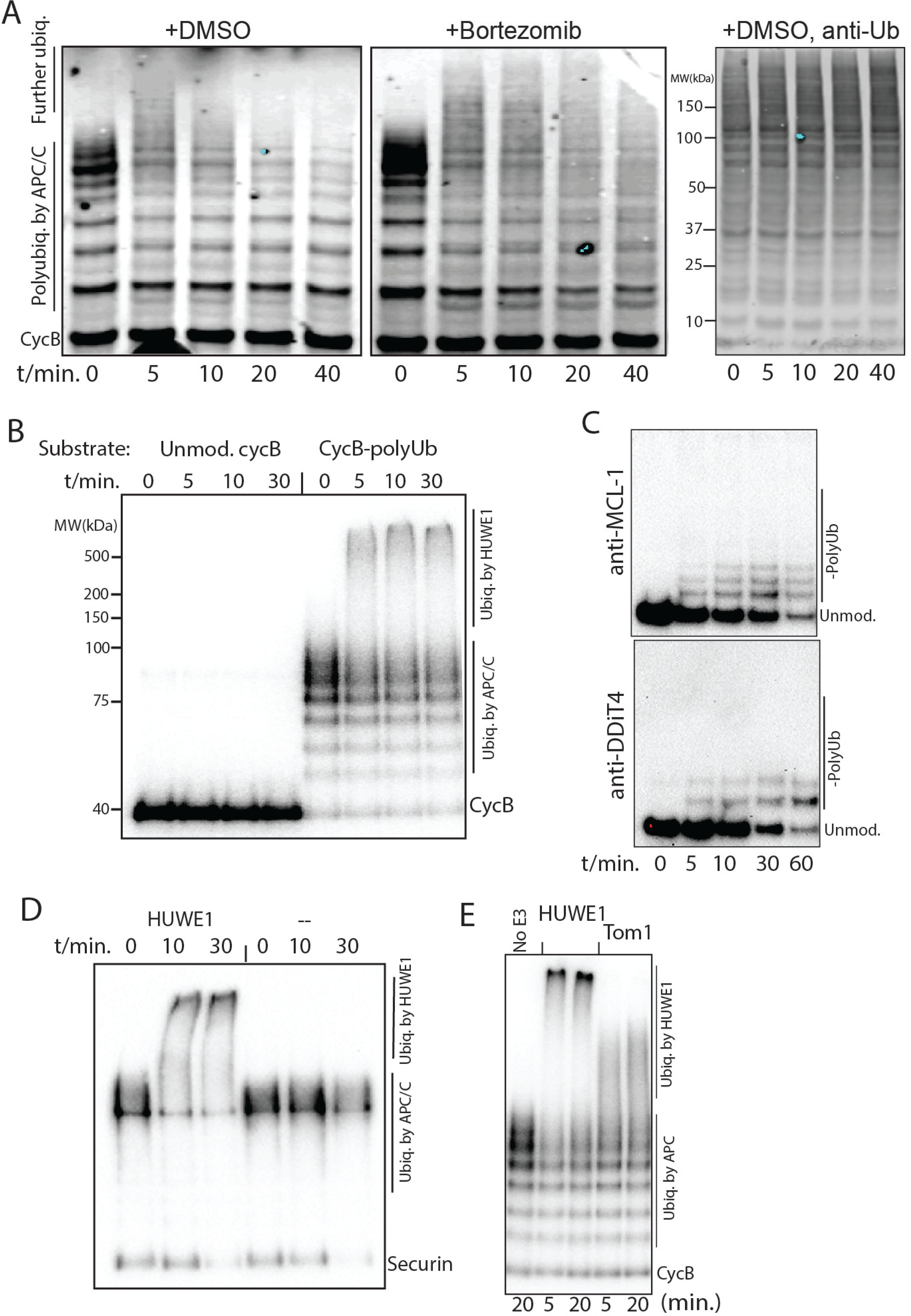
Identification of HUWE1 and its ubiquitin-directed ubiquitin ligase activity. **A.** HA-cycB- cpGFP-polyUb^L73F^ was incubated with HeLa extract and sampled at indicated time points. Anti-HA and anti-Ub western blotting were used to detect cycB ubiquitylation and the overall Ub distribution in cell extract. **B.** P32-labeled unmodified (unmod.) or polyubiquitylated cycB-cpGFP was tested in a ubiquitylation reaction mediated by rHUWE1. The samples were separated by SDS-PAGE and analyzed by autoradiography. **C.** Purified MCL-1 and DDIT4 were used as substrates for ubiquitylation by rHUWE1 and results were analyzed by western blotting. **D.** Polyubiquitylated securin was tested in a ubiquitylation reaction with rHUWE1 and analyzed by anti-securin western blotting. **E.** As in B, but using purified Tom1 as the E3.

To identify the factor expanding the ubiquitylation on cycB, we sequentially fractionated HeLa extract using anion exchange and size-exclusion chromatography and discovered several fractions possessing this activity (Fig. S5A; methods). An E2 scan suggests that UbcH5a/b/c was the primary E2 involved in this ubiquitylation (Fig. S5B). We next applied quantitative mass spectrometry to these fractions and identified the E3 HUWE1 whose concentration profile among these fractions closely matches the activity profile (Table S1). Immunodepleting HUWE1 from the fractionated extract using an antibody abolished the further ubiquitylation of cycB, while this activity was retained on the beads conjugated with anti-HUWE1 antibody (Fig. S6A). Importantly, recognition by HUWE1 does not require the L73F mutation on Ub, as WT Ub-conjugated cycB showed a similar reactivity (Fig. S6B). In most of our subsequent experiments, we used WT Ub to prepare Ub conjugates for studying HUWE1.

Previous studies have suggested that HUWE1 ubiquitylates important substrates such as p53, MCL-1 and DDIT4, and regulates a number of key biological process(*25*). However, the activity of expanding the preexisting Ub modification on substrates has not been reported for HUWE1. To further establish HUWE1 as the factor ubiquitylating cycB-polyUb, we purified recombinant HUWE1 (rHUWE1) from mammalian cells through a strep-tag (Fig. S7A)(*35*). rHUWE1 preferentially reacted with cycB carrying more than four Ubs, and not unmodified cycB(Fig. 1B), similar to the results in the crude extract (Fig. 1A), suggesting that it has a previously uncharacterized ligase activity that depends on the priming ubiquitylation. HUWE1-mediated ubiquitylation was highly processive, conjugating 30∼100 Ubs in a few minutes on cycB, even at a low HUWE1 concentration of 5nM (Fig. 1B; Fig. S7B). This processivity on cycB was higher than that by the APC-specific chain-extension factor Ube2S (Fig. S7C), tested at Ube2S’s physiological concentration of ∼100nM(*30, 36*), and was also higher than the processivity on MCL-1 and DDIT4 which are conventional HUWE1’s substrates that do not require a priming ubiquitylation (Fig. 1C)(*35*). rHUWE1 also ubiquitylated securin and geminin in a highly processive manner, but only after a priming ubiquitylation by the APC (Fig. 1D; Fig. S7D). No Ub was incorporated on these substrates without prior ubiquitylation by the APC. These results demonstrate that HUWE1 has two distinct Ub ligase activities with different processivities and different substrate specificities.

Purified Tom1, a HUWE1 ortholog in budding yeast, also ubiquitylated cycB-polyUb, albeit with lower processivity (Fig. 1E). Therefore, the Ub-expanding activity of HUWE1 appears to be a conserved feature across distant species.

### HUWE1’s UDL activity conjugates Ubs directly to the substrate molecule

To probe whether the Ub-directed conjugating activity of HUWE1 resembles an elongation factor (or E4 (*37*)) that is dedicated to elongating Ub chains(*21, 38*), we tested HUWE1 on purified K48-linked Ub chains. Unlike an elongation factor, HUWE1 did not extend free Ub chains (Fig. S8A). Similarly, purified UbL73F chains were also not further ubiquitylated in cell extract (Fig. S8B). To further characterize this Ub-expanding activity of HUWE1, we created a fusion protein containing cycB and the fluorescent protein mNeonGreen, separated by a TEV-protease recognition site. This allowed us to monitor the ubiquitylation status of cycB and mNeonGreen separately after cleavage by TEV protease (Fig. 2A). The APC predominantly ubiquitylated the cycB part of this substrate, leaving the C-terminal mNeonGreen mostly unmodified (Fig. 2B). In contrast, when HUWE1 was employed the mNeonGreen domain was extensively ubiquitylated. Thus, HUWE1-mediated ubiquitylation is triggered by the prior Ub modification on a substrate and adds Ubs directly to available sites on the substrate molecule. We call this novel activity “Ubiquitin-Directed ubiquitin Ligase”, or UDL.

**Figure 2.**
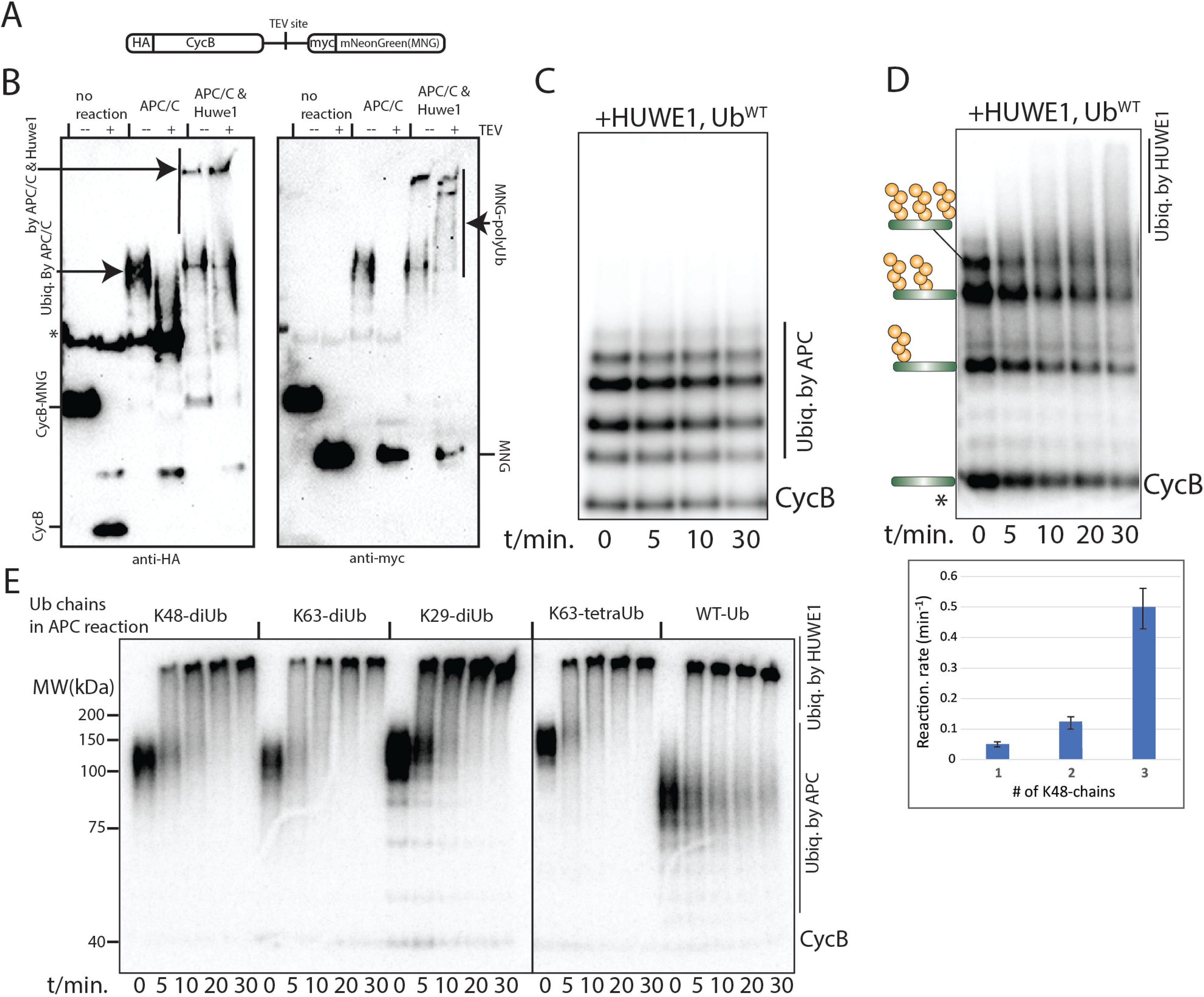
HUWE1 recognizes substrates carrying multiple Ub chains and conjugates additional Ubs directly on the substrate molecule. **A.** Design of the substrate. **B**. The substrate in A was ubiquitylated by APC-UbcH10, and subsequently by purified HUWE1. The products were cleaved with TEV protease and analyzed by anti-HA and anti-myc western blotting. **C**. P^32^-labeled cycB-cpGFP was multiple- monoubiquitylated by the APC plus methylated Ub (t=0) and was then reacted with purified HUWE1 in the presence of WT Ub. Products were separated by SDS electrophoresis and analyzed by autoradiography. **D**. K48-linked tetra-Ub chains were first methylated and conjugated to P^32^-labeled cycB-cpGFP via the APC. The product was then reacted with purified HUWE1. The decay constant (below) for each ubiquitylated species, separated on a gel, was measured from a time series. The error bars represent the standard deviation of three replicates. *Signal from unmodified cycB was used as a control for loading and nonspecific dephosphorylation (see methods). **E**. P^32^-labeled cycB-cpGFP was ubiquitylated by the APC with indicated Ub chains. The products were then reacted with purified HUWE1 plus WT Ub, and were analyzed by autoradiography.

We next examined which Ub configurations on substrate can trigger HUWE1’s UDL activity. HUWE1 recognition requires Ub chains, as multiple-monoubiquitylated cycB-cpGFP showed little reactivity (Fig. 2C). Using the APC we conjugated varying numbers of preformed K48-linked tetra-Ub chains on cycB(Fig. 2D)(*33*). These chains were methylated before conjugation to prevent chain elongation. The different ubiquitylated species were separated on a gel and their reaction rates were measured. This result suggests that HUWE1 preferentially reacts with substrates conjugated with more than one Ub chains.

To identify the Ub linkages formed by HUWE1’s UDL activity, we tested single-lysine variants of Ub in HUWE1 reactions. We found that both K48-only and K11-only Ub, and to a lesser extent K6- only Ub, support efficient ubiquitylation, consistent with a previous characterization of the Ub-chain linkages formed by HUWE1’s HECT domain (Fig. S9)(*39*). Since HUWE1 can conjugate 6∼15 methylated Ubs on cycB-polyUb (Fig. S9), it is likely to form several Ub chains on the substrate molecule in the presence of WT Ub. Because multiple Ub chains are also the recognition signal by HUWE1, more Ub chains may boost the substrate’s reactivity with HUWE1, resulting in a positive feedback to rapidly expand substrate’s ubiquitylation (see Discussion).

Because methylated Ub chains cannot be extended, the result in Fig. 2D provides further evidence that HUWE1 utilizes new sites on the substrate for conjugation, rather than extending pre-conjugated chains. Substrates carrying shorter Ub chains or chains with alternative linkages (K29 or K63-linked di- Ub chains) were also subject to HUWE1’s UDL activity (Fig. 2E), suggesting that the number of Ub chains, but not the length or linkage, is important for HUWE1’s UDL activity.

### HUWE1 recognizes the local density of Ub chains

Does recognition by HUWE1 require other features on substrate in addition to multiple Ub chains? To address this question, we constructed a synthetic substrate using a linear tri-Ub chain fused with a small tetramerization domain from the Vasodilator-stimulated phosphoprotein (VASP) and GFP (Fig. 3A)(*40*). HUWE1 efficiently ubiquitylated this synthetic substrate, with a similar processivity as we observed for cycB-polyUb, while a variant of this substrate with mono-Ub at the N-terminus showed little reactivity (Fig. 3A,B). Therefore, multiple Ub chains on a substrate are both necessary and sufficient for HUWE1’s UDL activity.

**Figure 3.**
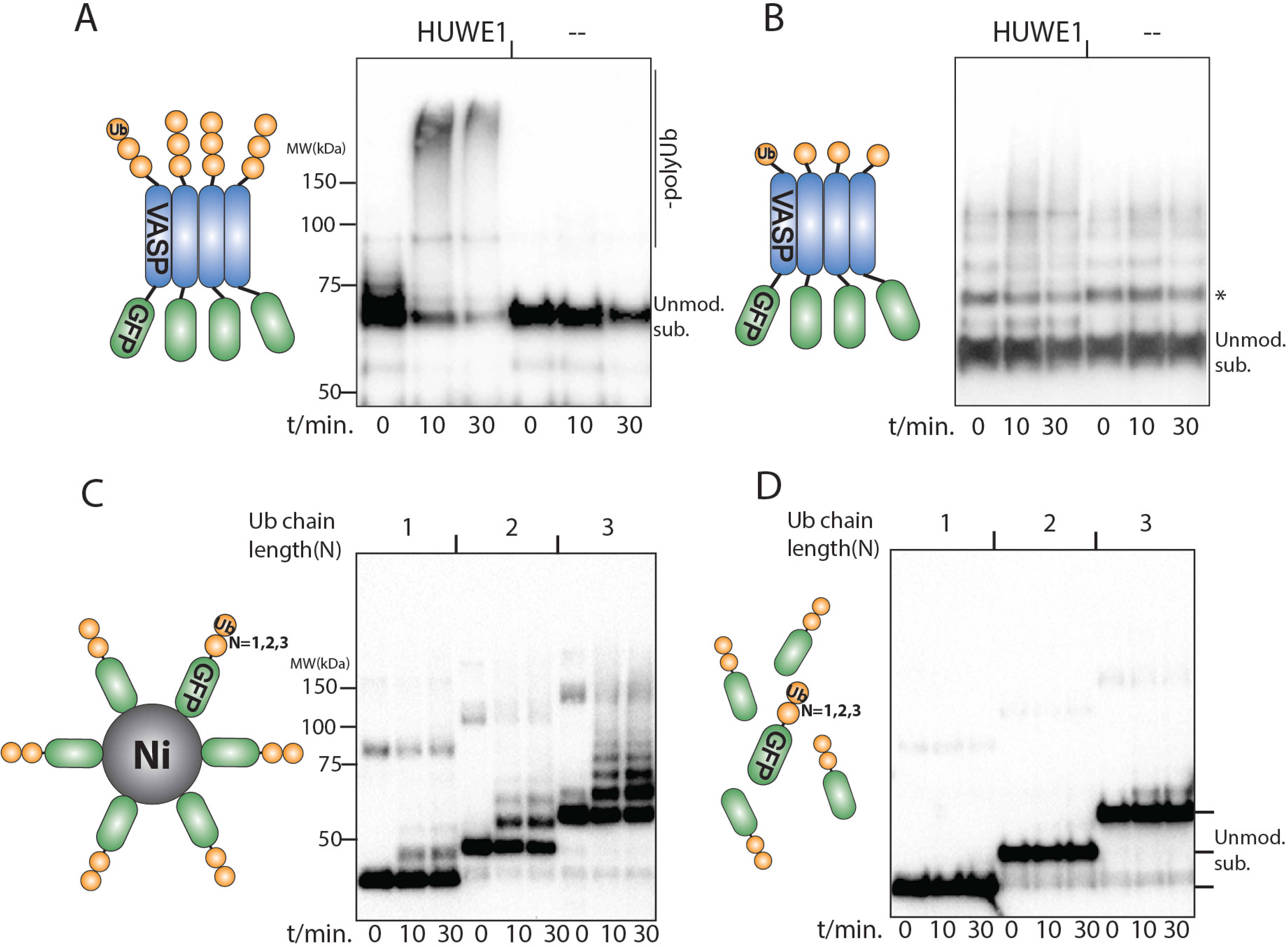
HUWE1 reacts with Ub-rich protein aggregates. **A**. The HA-tagged synthetic substrate (Left) was reacted with 50nM rHUWE1 plus WT Ub. The product was analyzed by anti-HA western blotting. **B**. As in A, but using an alternative substrate design (Left) with a monoUb at the N-terminus. “*” is a nonspecific band. **C**. HA-tagged GFP with 1,2 or 3 Ubs at the N-terminus and a polyHis-tag at the C-terminus was incubated in a rHUWE1 ubiquitylation reaction in the presence of Ni-NTA magnetic particles. Results were analyzed by anti-HA western blotting. **D**. As in C, but in the absence of Ni particles.

Based on these observations, we hypothesized that HUWE1’s specificity towards substrates with a high local density of Ub chains may target it to the often Ub-rich protein aggregates/condensates, such as those associated with neurodegenerative diseases, and expand their Ub modifications. To test this in a chemically-defined system, we purified GFP fused with 1∼3 Ubs at its N-terminus (Fig. 3C). rHUWE1 had little reactivity with these proteins in solution. However, when Nickel beads were added to condense these substrates via a 6xHis tag at their C-terminus, we detected efficient ubiquitylation by rHUWE1, except on Ub(1)-GFP, consistent with our hypothesis (Fig. 3C,D).

In conclusion, the UDL activity appears to target substrates with a high local density of Ub chains that can be formed by either poly-ubiquitylation or through protein condensation.

### The ubiquitin-binding domains on HUWE1 mediate the UDL activity

HUWE1 contains five ubiquitin-binding domains(UBDs) including one UBA, one UIM and three UBM domains(*35*). Mutating these domains lowers the rate of E2 discharging, but they are not required for ubiquitylating the UDL-independent substrates such as MCL-1 and DDIT4 (Fig. S10A)(*35*). By contrast, simultaneous mutation of the UBA, the UIM and one of the UBM domains (HUWE1^UBDmu^) severely compromised the UDL activity, even at high E3 concentrations, while mutating only the UBA and the UIM domains attenuated the processivity of ubiquitylation (Fig. S10B). Consistently, HUWE1^UBDmu^ was defective in ubiquitylating geminin-polyUb (Fig. S7D). Thus, the recognition of the multi-chain signal on substrates appears to require the UBDs on HUWE1, perhaps driven by these UBDs’ avidity with the substrate’s Ub chains.

Several mutations on HUWE1 have been linked to intellectual disability(*27, 35*). We purified and tested four patient variants of HUWE1 associated with neurodevelopmental disorders(*27*) and found that they showed defects in the UDL activity with differing severity (Fig. S10C). Since these mutations are outside HUWE1’s UBDs, this result indicates that regulation of the UDL activity involves other regions on HUWE1, which may underlie the pathogenicity of HUWE1 mutations.

### HUWE1 stimulates proteasomal degradation of soluble substrates *in vitro*

We next dissected the function of HUWE1’s UDL activity in controlling proteasome and p97/VCP-mediated substrate processing in reconstituted reactions. To study proteasomal degradation, we tested substrates that are resistant to processing by the 26S proteasome in a reconstituted ubiquitylation- degradation system (RUDS) comprising wtUb and other purified components. HUWE1 stimulated the initial degradation rate of cycB-cpGFP-polyUb^L73F^ as indicated by the loss of fluorescent signal (Fig. S11A). However the degradation rate trended lower at later time points when HUWE1 was present. We surmise that, although higher Ub stoichiometry generally favors faster proteasomal degradation, excessive ubiquitylation on substrates due to the unbridled processivity of HUWE1 in the *in vitro* reaction impede proteasomal degradation. Consistent with this hypothesis, when methylated Ub was added to the RUDS to limit the processivity of HUWE1, we observed a robust stimulation of the degradation of cycB-cpGFP by more than two-fold (Fig. S11A). This result indicates that a possible role of UDL activity is to rescue a defective degradation signal (see Discussion). The stimulation of degradation is not unique to Ub^L73F^ conjugates. Instead of deubiquitylation, proteasomal degradation can also be limited by unfolding of structurally-stable domains, such as EGFP. We observed a similar stimulation of degradation of cycB- EGFP-polyUb^WT^ in the RUDS when HUWE1 was involved (Fig. S11B).

To examine further the role of HUWE1 in protein degradation, we briefly reacted cycB-cpGFP- polyUb^L73F^ with purified HUWE1 in the presence of WT Ub and quenched the reaction using an E1 inhibitor. This procedure generated cycB substrates with a wide range of Ub stoichiometries, as apparent from SDS gel electrophoresis (Fig. S11C). We then determined the degradation rates of cycB, conjugated with either Ub and Ub^L73F^ by APC, and found that HUWE1-mediated further ubiquitylation markedly enhanced the breakdown of cycB by the 26S proteasome (Fig. S11C). At very high levels of ubiquitylation, cycB’s degradation rate decreased, consistent with the previous results.

The over-ubiquitylation in the *in vitro* reaction may be corrected by the deubiquitylating activity in the cell. Indeed, the extract of HeLa S3 cells contains strong deubiquitylating activity that completely deconjugated polyubiquitylated cyclinB and securin within five minutes, even in the absence of proteasome activity (Fig. S12A,B). We next studied whether HUWE1’s UDL activity enhances protein degradation in the presence of deubiquitylation. In a RUDS that contains the 26S proteasome, HUWE1 and the E3 APC, as well as their cognate E2s, we added unmodified cycB-cpGFP as the substrate and titrated the concentration of a deubiquitinase (DUB) Usp2 that has little specificity for Ub-chain linkages. In the absence of HUWE1, the degradation rate of cycB-cpGFP dropped steeply above 300nM Usp2, which was likely due to a decrease of substrate ubiquitylation (Fig. 4A). In the presence of HUWE1, the substrate’s degradation rate first increased with increasing Usp2 concentration, before the rate dropped at higher Usp2 levels (Fig. 4A; Fig. S12C). With HUWE1, the degradation rate also decreased less. Thus, HUWE1 appears to stimulate proteasomal degradation in the presence of active deubiquitylation and maintains the degradation rate at different DUB activities.

**Figure 4.**
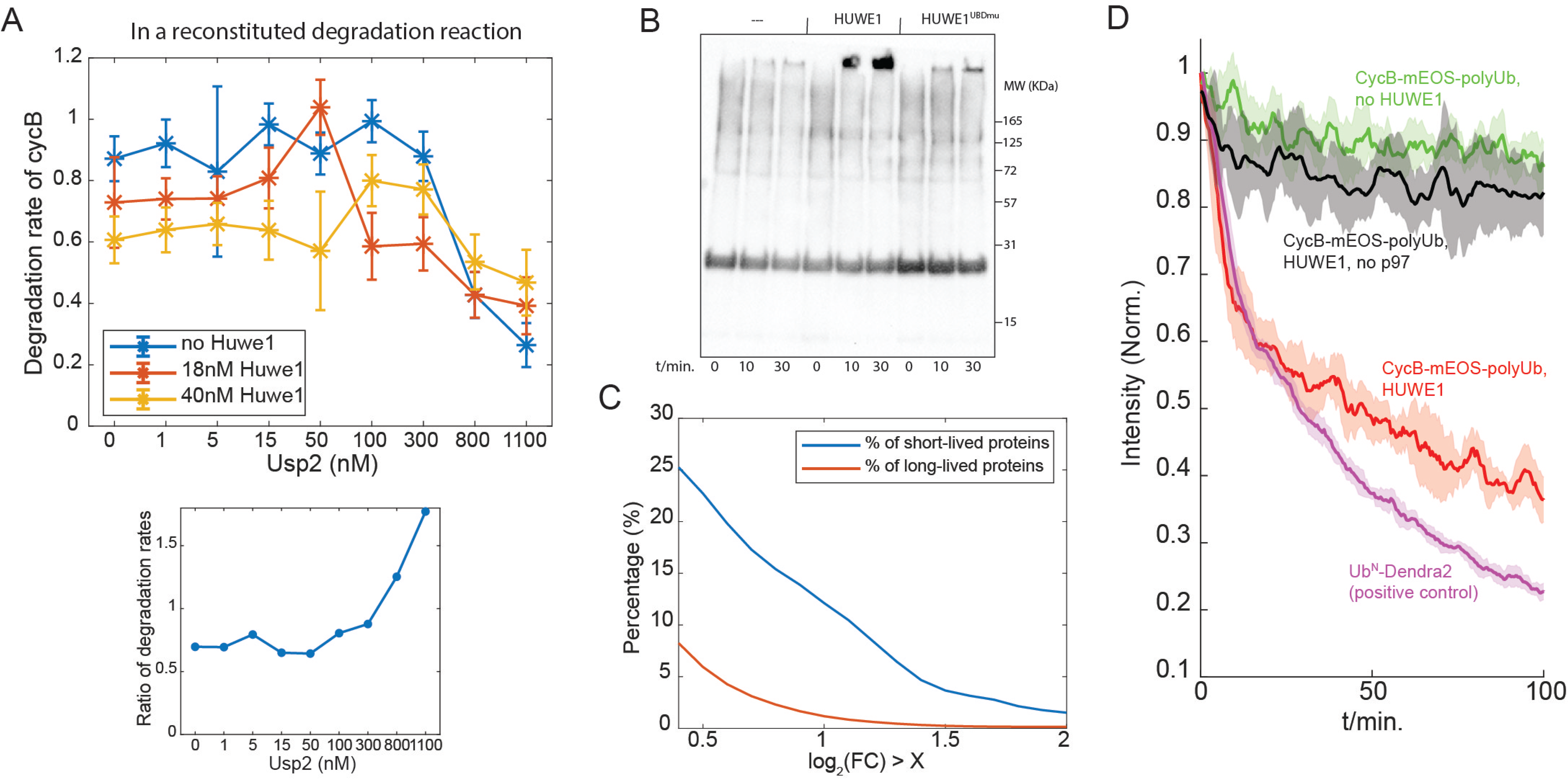
HUWE1’s UDL activity stimulates proteasomal degradation and p97-mediated substrate unfolding. **A.** Degradation rate of cycB, as indicated by the fluorescence intensity, was determined at different Usp2 concentrations in a RUDS containing the proteasome, HUWE1, the APC, WT Ub and unmodified cycB-cpGFP as the substrate. The ratio of the degradation rate with 40nM HUWE1 over that without HUWE1 is presented below at each Usp2 concentration. Error bars represent fitting uncertainty. **B.** Endogenous Ub conjugates were isolated from HEK293 cells expressing His6-HA-Ub using Ni- affinity chromatography (see methods). Eluted conjugates were incubated with purified HUWE1 or HUWE1^UBDmu^ in a ubiquitylation reaction with WT Ub and the samples were analyzed by anti-HA western blotting. **C.** The fold changes (FC) of protein abundance in HUWE1Δ cells were determined by quantitative mass spectrometry. The percentage of short-lived protein (N=264, defined in (*43*)) or long- lived proteins (N=5449, defined in (*36*)) whose level increase is larger than X (& p<0.05) was plotted. **D.** CycB-mEOS was first ubiquitylated by APC-UbcH10, and was incubated with recombinant human p97- UFD1-NPL4 complex, either without (green) or with (red) further ubiquitylation by rHUWE1. Ubiquitylated dendra was used as a positive control (*48*).

### HUWE1-mediated Ub signal amplification is a common step in protein degradation

The specificity for recognizing the Ub modification, instead of the substrate protein, suggests that HUWE1 reacts with a variety of cellular substrates. To test this possibility, we attempted to assay HUWE1 on a complex collection of natural substrates. We created a cell line stably expressing polyHis- HA-tagged Ub and purified Ub-modified endogenous proteins using Ni-affinity chromatography. Western blot analysis revealed a prominent retardation of migration of a fraction of these conjugates, suggesting further ubiquitylation, upon incubation with purified HUWE1 in a ubiquitylation reaction (Fig. 4B). In particular, the UDL, instead of the conventional E3 activity, of HUWE1 was mainly responsible for this ubiquitylation, as mutating HUWE1’s UBDs largely abolished this activity.

To evaluate the effect of HUWE1 on endogenous proteasome substrates, we performed deep global proteomic analysis comparing a HEK293 cell line in which the HUWE1 gene has been knocked out using CRISPR and WT HEK293(*41*). ∼9000 proteins were quantified (Table S2). Most short-lived proteins are degraded by the ubiquitin-proteasome system (UPS) (*42*). Our data suggests that the levels of ∼25% short-lived proteins (half-life shorter than 8hrs), as defined in a previous study(*43*), significantly increased, by their mass fractions, in HUWE1Δ cells (Fig. 4C; Fig. S13), while long-lived proteins (half-life longer than 24hrs(*36*)) were much less affected. HUWE1 knockout did not appear to cause a proteotoxic stress under a normal growth condition, as the level of Ub conjugates in HUWE1Δ was similar to that in WT cells (Fig. S14). Gene ontology analysis based on significantly up- or down- regulated proteins in HUWE1Δ cells also did not identify pathways involved in stress response or proteotoxicity (Fig. S15)(*44*). Stabilization of UPS substrates is neither caused by transcriptional changes because only ∼0.2% transcripts’ levels increase by more than two-fold due to HUWE1 knockout in different cell lines (Fig. S16) (*45*). Therefore, these results suggest that HUWE1-mediated Ub amplification is likely a general step acting on a fraction of endogenous substrates to promote their efficient degradation. Nonetheless, HUWE1 is not a nonselective activator of the UPS nor does it specifically promote the degradation of fluorescent proteins. Notably, the degradation of a N-end rule substrate Ub-R-YFP was not affected by knocking out HUWE1 (Fig. S17A). The obvious selectivity of UDL adds another layer of control to the degradation system, which may regulate important biological processes (see Discussion).

### HUWE1 promotes substrate unfolding by the p97/VCP complex

Along with the proteasome, the p97/VCP ATPase plays a key role in the Ub-mediated degradations. Previous studies suggest that the p97 complex preferentially recognizes substrates with a long Ub chain(*46, 47*). We therefore tested whether the UDL activity can promote substrate unfolding by p97 *in vitro.* Self-cleaved fluorescent proteins, such as mEOS and dendra, lose fluorescent signal upon unfolding by p97(*48*). Although dendra carrying a long Ub chain (Ub^N^-dendra) was unfolded efficiently by recombinant human p97 with its substrate receptors Npl4 and Ufd1, cycB-mEOS-polyUb was poorly unfolded (Fig. 4D), even though cycB contained a similar number of Ubs to the former (Fig. S18; S1A), likely because APC-UbcH10 preferentially forms multiple short chains (mostly shorter than three Ubs) on cycB(*49*). In contrast, further ubiquitylation by HUWE1 on cycB-mEOS dramatically enhanced its unfolding by p97 (Fig. 4D). A similar effect was observed for the tetramer substrate, Ub(3)-VASP-GFP, when GroEL(D87K) was added to prevent GFP refolding (Fig. S19A)(*50, 51*). Interestingly, Ub(3)- VASP-GFP was still poorly degraded by proteasome after HUWE1 reaction, even after a degradation initiation sequence was added to the substrate (Fig. S19B)(*29*). This may be due to the unhinged processivity of HUWE1 *in vitro* that compromised the reactivity with proteasome. Overall, HUWE1’s UDL activity consistently stimulates p97-mediated unfolding of protein substrates.

### HUWE1 facilitates the degradation of pharmacological targets

To investigate the connection between HUWE1 and the p97 in degrading soluble factors, we examined the substrates of Targeted Protein Degradation (TPD), induced by the Proteolysis Targeting Chimera (PROTAC) or molecular glues. TPD has become an important therapeutic strategy in eliminating disease-causing factors(*52*). Cullin RING ligase (SCF) E3s, such as VHL, DCAF15, and Cereblon, are common mediators for TPD, raising the possibility that HUWE1 may promote TPD via its UDL activity. To test this suggestion, we compared the changes of pharmacological targets in WT and HUWE1Δ cells after treating cells with pan-kinase degraders SK-3-91 or DB-0646 (*24*). These two compounds recruit Cereblon to degrade a panel of kinase targets, including AURKA, CDK5, CDK12 etc. Notably, degradation of these kinases by PROTAC commonly requires p97 activity(*24*). HUWE1 knockout stabilized most of the targets of SK-3-91 and DB-0646 in HEK293 cells to a similar degree, as inhibition of p97 activity, with Pearson correlation coefficients ≈ 0.4 and 0.8 (Fig. 5A,B; Table S3). This result further supports a functional connection of HUWE1’s UDL activity with the p97-mediated unfolding and degradation.

**Figure 5.**
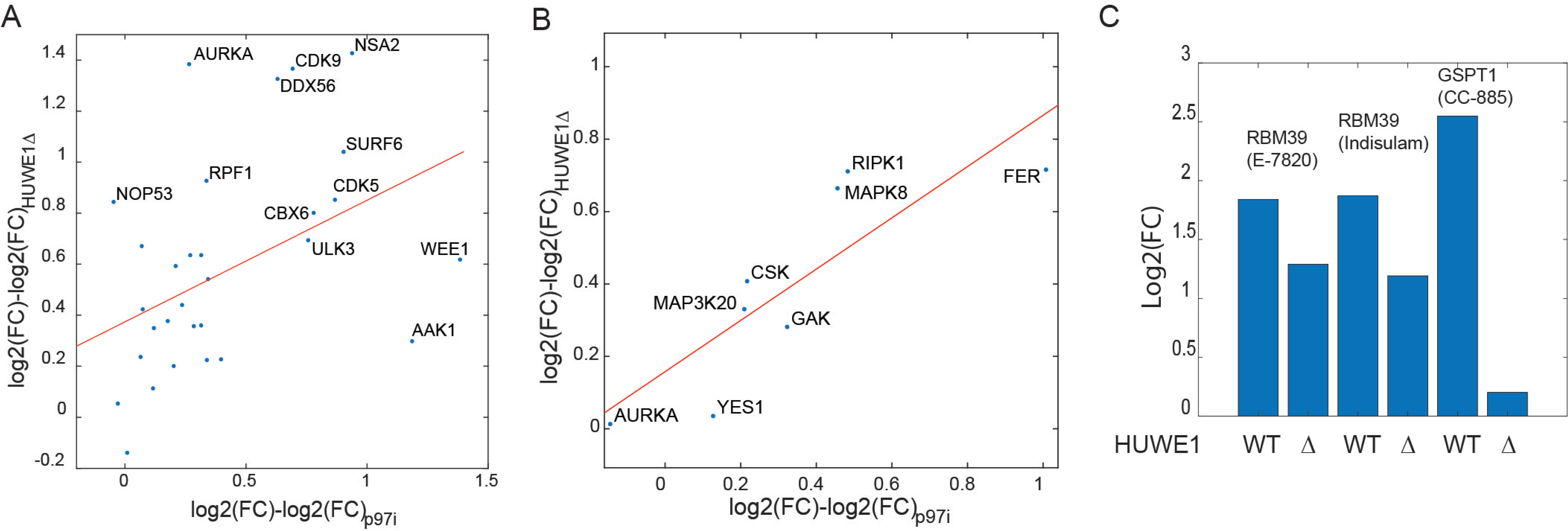
HUWE1 mediates efficient Targeted Protein Degradation. A, B. WT HEK293 and HUWE1Δ cells were treated with kinase degraders SK-3-91 (A) or DB-0646 (B). The fold-change (FC) of each kinase targeted by the compound (defined in Donovan KA, et al., *Cell* (2020)) was determined by mass spectrometry (Table S3) and the difference in WT and HUWE1Δ cells was correlated with the difference in the presence and absence of a p97 inhibitor. The p97 inhibitor result is from Donovan KA, et al., Cell (2020). **C.** As in A, but using indicated degrader compounds. The FC of each target in WT or HUWE1Δ cells was presented.

HUWE1 also facilitates the activities of molecular-glue degraders, such as E-7820, indisulam and CC-885 that degrade important therapeutic targets RBM39 and GSPT1 by recruiting DCAF15 and Cereblon respectively, as the induced degradations of these substrates were compromised in HUWE1Δ cells (Fig. 5C; Table S3).

### HUWE1’s UDL activity stimulates the degradation of cell-cycle regulators and is important for the sharp transitions between cell-cycle stages

We next explored the biological functions of the UDL activity *in vivo*.

The master regulator E3s of the cell cycle, the APC and the SCF, typically conjugate multiple Ub chains on their substrates(*49, 53–56*). Our findings raised the possibility that multiple Ub chains may be used to target these substrates to HUWE1’s UDL activity, suggesting HUWE1 as possible regulator of cell cycle progression. We tested this possibility in live cells using quantitative imaging.

The levels of geminin and Cdt1 oscillate during the cell cycle to regulate DNA replication. The degradation of geminin and Cdt1 are mediated respectively by the APC in M phase and by the SCF in S phase. The FUCCI cell-cycle markers are based on the ubiquitylation domains of geminin and Cdt1 coupled to fluorescent proteins, and have been widely used to study cell cycle progression in live cells(*57, 58*). We stably expressed these markers in a mutant HEK293 cell line in which the HUWE1 gene has been knocked out using CRISPR, and monitored their levels in unsynchronized cell cycles using timelapse microscopy (Fig. 6A,B)(*59*). Although both WT and HUWE1Δ cells showed a similar steady-state distribution of cell-cycle stages by DNA content, the HUWE1Δ cells grew slightly more slowly than WT HEK293 cells (Fig. S20A,B), suggesting a defect in cell cycle progression.

**Figure 6.**
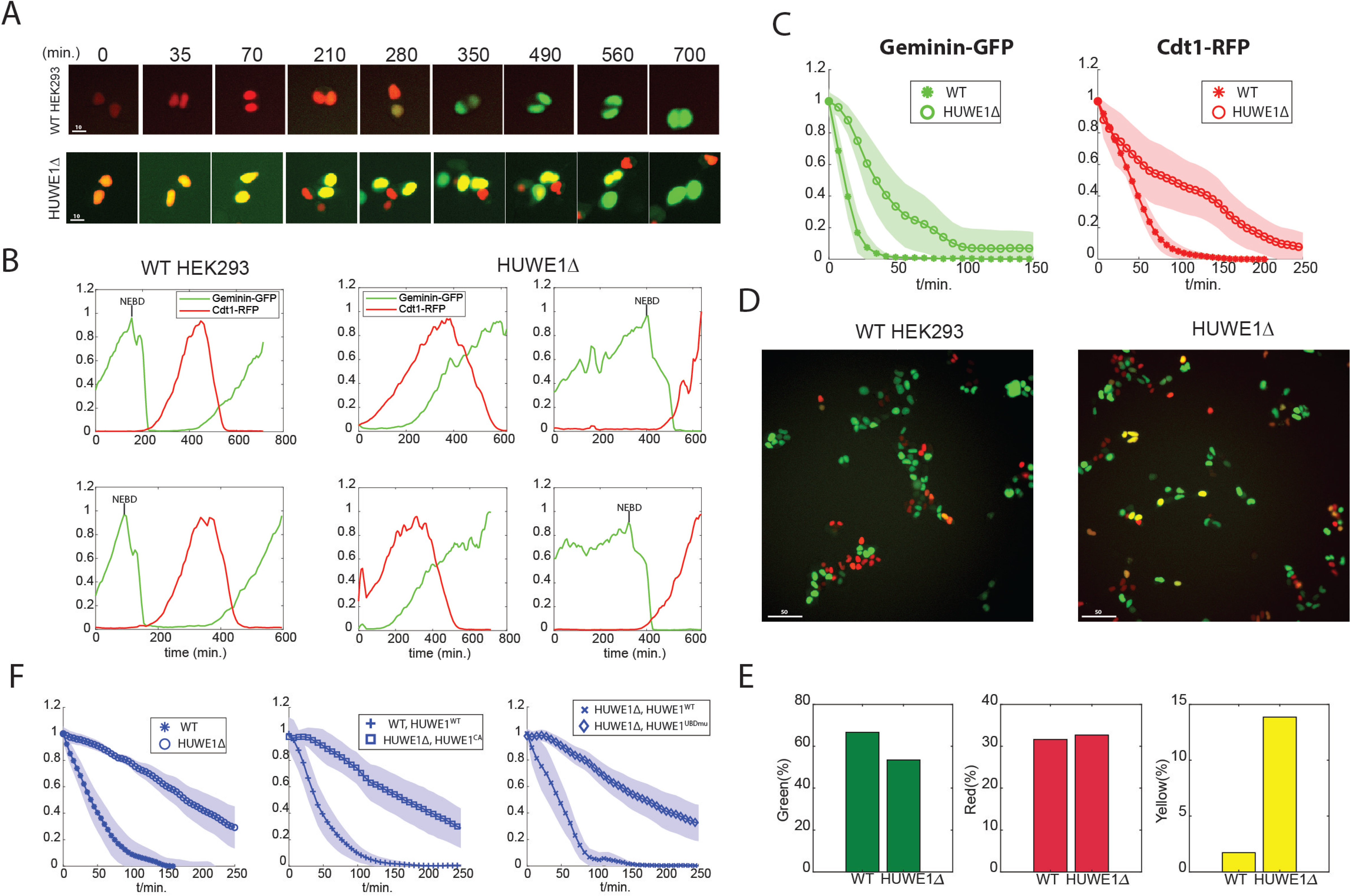
HUWE1’s UDL activity ensures the timely degradation of cell-cycle regulators and is important for the sharp transitions between cell-cycle stages. A. Image montage of exemplary WT HEK293 and HUWE1Δ cells expressing the FUCCI cell-cycle markers, captured by timelapse microscopy. Scale bar unit: μm. **B.** Example traces of geminin-GFP and Cdt1-RFP in WT or HUWE1Δ cells. **C.** The average decay kinetics of geminin and Cdt1. Geminin traces (N=32) in single cells were normalized and aligned by the moment of the nuclear envelope breakdown (NEBD); Cdt1 traces (N=35) were aligned by their peaks. The averaged intensities were plotted over time. The shaped regions represent the standard deviations. **D.** Example images of an asynchronous population of WT and HUWE1Δ cells expressing FUCCI markers. **E.** The fractions of geminin-expressing (green), Cdt1- expressing (red), and both-expressing (yellow) cells of either WT or HUWE1Δ. **F.** The average decay kinetics of securin-YFP in WT or HUWE1Δ cells. Single-cell securin traces were aligned by the moment of NEBD and processed as in C. Cells were co-transfected with HUWE1^WT^, or the catalytically-inactive HUWE1^CA^, or the UBD mutant HUWE1^UBDmu^ with a RFP tag at the N-terminus to track the expression level. Total N=114.

We analyzed the average kinetics of degradation for these FUCCI markers, aligning the geminin traces of individual cells by the timepoint of nuclear envelope breakdown. Individual Cdt1 traces were aligned at their intensity peaks. The averaged kinetics suggest that both geminin and Cdt1 were degraded significantly more slowly in HUWE1Δ cells, by a factor of two to three (Fig. 6C). These results suggest that HUWE1-mediated degradation may facilitate cell cycle progression in all stages.

In WT cells, geminin and Cdt1 are expressed in different phases of the cell cycle and only overlap briefly at the G1/S boundary(*58*). Interestingly, the temporal expression patterns of geminin-GFP and Cdt1-RFP overlapped significantly in HUWE1Δ cells, especially in the S phase, which was likely due to the sluggish degradation of these factors in the absence of HUWE1 (Fig. 6B). HUWE1 knockout thus led to a dramatic increase of the number of “yellow” cells that expressed both geminin and Cdt1 FUCCI markers in an unsynchronized population (Fig. 6D,E). Therefore, by ensuring timely degradation of cell-cycle regulators such as geminin and Cdt1, HUWE1 helps to set the boundary between cell-cycle stages and establishes rapid and concerted cell-cycle transitions.

To determine whether these effects on the degradation of cell-cycle regulators are due to HUWE1’s UDL or its conventional E3 activity, we transiently expressed securin-YFP in HUWE1Δ cells together with either WT HUWE1 or variants with mutations in its UBDs or at its catalytic site (Fig. S17B). The HUWE1 variants were tagged with a red fluorescent protein (RFP) at the N-terminus to track their expression levels (Fig. S17C). Similar to what we found with geminin and Cdt1, degradation of securin during M phase was significantly compromised in HUWE1Δ cells (Fig. 6F). As expected, co-expressing WT HUWE1 rescued the degradation of securin, but neither the catalytically-inactive (HUWE1^CS^) nor the UBD mutant (HUWE1^UBDmu^), was able to rescue the degradation of securin in HUWE1Δ cells (Fig. 6F). Notable, the HUWE1^UBDmu^ retained the conventional E3 activity (Fig. S10A), but not the UDL activity, we conclude that HUWE1’s UDL activity is essential for the timely degradation of cell-cycle substrates, consistent with our results in chemically-defined ubiquitylation/degradation systems.

### HUWE1 colocalizes with cellular protein aggregates and promotes their clearance

The *in vitro* observations that HUWE1 targets Ub-rich aggregates and promotes p97-mediated substrate unfolding (Fig. 3; Fig. S19) suggest the possibility that HUWE1 mediates the clearance of cellular protein aggregates. To test this, we employed a chemically-inducible aggregation system, agDD, which is based on fusing a FKBP domain with a short hydrophobic peptide(*60*). AgDD-GFP aggregates were visible in HEK293 cells within 15 minutes once shield-1, a stabilizing ligand for FKBP, was removed from the medium (Fig. S21A; Supp. movie 1), and further congregated at a perinuclear region, resembling the aggresome which is a common cellular response to the accumulation of misfolded proteins (Fig. S21B)(*61*). Consistent with their aggresome designation, formation of the perinuclear agDD aggregates depends on intact microtubule structure and HDAC6 activity(*62, 63*), both are known for their roles in aggresome formation (Fig. S21C). Importantly, unlike most conditions for inducing the formation of protein aggregates/condensates, agDD aggregation did not appear to affect the growth or division of WT HEK293 cells(*60*), which strongly simplifies data interpretation.

Cells constitutively expressing a low level of agDD did not form aggregates, while these aggregates were stable in high-expression cells (Fig. S22A,B). AgDD aggregates, at intermediate expression levels, tend to undergo spontaneous clearance within a few hours after induction (Fig. S22A), in a process that depends on active ubiquitylation and the p97 activity, as either inhibitor abolished agDD clearance (Fig. S22C). Adding cycloheximide to stop new protein synthesis promoted the clearance of most agDD aggregates, regardless of the expression level (Fig. S22D). AgDD clearance was not affected by the lysosome inhibitor chloroquine (Fig. S22C).

We tested the kinetics of agDD aggresome clearance using timelapse microscopy in WT and HUWE1Δ cells stably expressing agDD-GFP. Cycloheximide was added 30 minutes after removing shield-1 to facilitate clearance. By tracking the intensity of individual agDD aggregates in live cells, we found that HUWE1Δ dramatically reduced the rate of agDD dissolution (Fig. 7A,B; Fig. S23). We defined the clearance of agDD aggregates as when their intensities decreased by more than 50%, and found that HUWE1 knockout dramatically slowed down the clearance kinetics in both HUWE1Δ clones (Fig. 7C).

**Figure 7.**
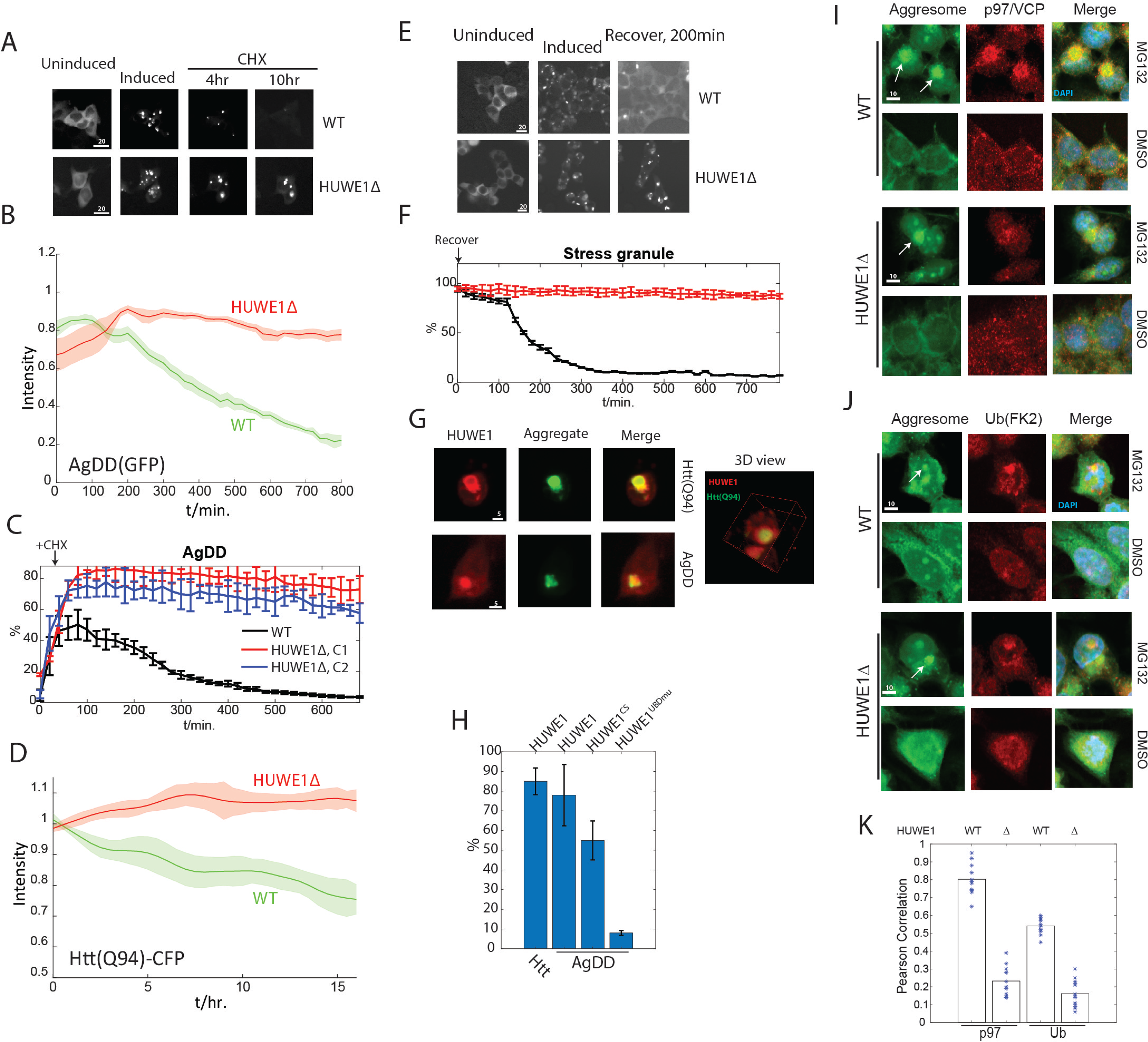
HUWE1 is required for the clearance of cellular aggregates/condensates. **A.** WT or HUWE1Δ cell lines stably expressing agDD(GFP) was induced to form aggregates. Cycloheximide(CHX) was added after 30 minutes to trigger clearance, and images were taken at indicated time points. Scale bar unit: μm. **B.** The intensities of individual agDD aggresomes in WT or HUWE1Δ cells (N=38) were tracked using timelapse microscopy. The average intensity traces were plotted vs time. Shade indicates standard error. **C.** The percentage of agDD-expressing cells with remaining agDD aggresomes were analyzed by timelapse microscopy after CHX was added to induce clearance (methods). C1, C2 are two clones of HUWE1 knockout. Error bars represent the standard deviation of three replicates. **D.** The average intensity traces of Htt(Q94) aggregates in WT or HUWE1Δ cells, tracked using timelapse microscopy (N=42). Htt expression was turned off at t=0 by withdrawing doxycycline to induce clearance. **E.** Stress granules, marked by stably-expressing GFP-G3BP1 in WT or HUWE1Δ cells, were induced by Arsenite treatment. Arsenite was then washed off to induce stress granule clearance in the recovery phase. The percentage of cells with remaining stress granules were analyzed by timelapse microscopy and ImageJ (methods), presented in **F.** Errorbars represent the standard deviation of three replicates. **G.** RFP-HUWE1 was transiently expressed in either AgDD or Htt(Q94)-CFP cell lines. Images were taken using confocal microscopy, 6hr (agDD) or 20hr (Htt) after inducing aggregation (methods). **H.** As in G. The percentage of colocalization of RFP-HUWE1 or RFP- HUWE1 variants with indicated aggregates were counted, 6hr (agDD) or 20hr (Htt) after inducing aggregation. **I, J.** Immunofluorescence images, by confocal microscopy, analyzing the colocalization of aggresome with p97 or Ub conjugates in WT or HUWE1Δ cells. Aggresome was induced by MG132 treatment and was stained by ProteoStat, marked by white arrows. **K.** Quantification of the colocalization of aggresome with the immunofluorescence signal of p97 or Ub conjugates in WT or HUWE1Δ cells, as suggested Pearson Correlation coefficient. Cells were treated as in I, J. Images were analyzed by ImageJ (methods). Marks represent individual cells.

We next studied HUWE1 in clearing disease-associated protein aggregates. We stably expressed synphilin-1, which is associated with Parkinson’s disease, and triggered its aggresome formation by treating the cells with a proteasome inhibitor MG132(*64*). MG132 was then removed to initiate synphilin aggresome clearance. We found, in a timelapse study, that HUWE1Δ significantly slowed the clearance rate of synphilin aggresomes (Fig. S24A,B). Furthermore, overexpression of HUWE1, but not its catalytically-inactive HUWE1^CS^ variant or the UBD mutant, significantly suppressed the accumulation of the high expression of synphilin (Fig. S24C).

Aggregates formed by poly-glutamine proteins, such as Huntingtin (Htt), are involved in the etiology of Huntington’s disease and ataxias. Clearance of the aggresome-like inclusions by Htt(Q94)- CFP was slow in HEK293 cells after the promoter was turned off. Most cells, either WT or HUWE1Δ, still contained Htt aggregates after 24 hours, but by tracking individual Htt aggregates, we found a much faster reduction in Htt aggregate’s intensity in WT cells after Htt expression was terminated (Fig. 7D).

Besides the aggresomes, HUWE1 also appeared important for the timely dissolution of G3BP1- GFP-marked stress granules, triggered by either Arsenite treatment or heat shock, after the stressors were relieved (Fig. 7E,F; Fig. S25).

To study whether HUWE1 directly interacts with protein aggregates via their Ub modifications, we transiently expressed RFP-HUWE1 in cells harboring the aggregates of either agDD or Htt(Q94). RFP-HUWE1 colocalized with both aggregates and formed a “shell-like” structure that encapsulated the Htt aggregates (Fig. 7G). This colocalization is mediated by the Ub signal on protein aggregates, as mutating HUWE1’s UBDs abolished the colocalization (Fig. 7H; Fig. S26). Overexpressing the catalytically-inactive HUWE1^CS^ partially compromised its localization with the aggregates (Fig. 7H; Fig. S26).

To further investigate whether HUWE1 targets protein aggregates in animals, we examined the localization of HUWE1 in the cerebral cortex of an Alzheimer’s disease mouse model (THY-Tau22) expressing disease-causing tau variants (G272V and P301S)(*65*). Immunohistochemical staining revealed the expression of HUWE1 in cortical neurons in both WT and mutant mice. There was a substantial colocalization of HUWE1 with tau aggregates, by Manders’ overlap coefficient, in neurons at the early stage of tau aggregate formation; such colocalization increased as the aggregates matured but completely disappeared at the end stage when tau formed stable fibrils accompanied by neuronal loss (Fig. S27).

To examine whether HUWE1 is important for the ubiquitylation of protein aggregates and their recognition by the segregase p97/VCP, aggresomes were stained with the dye “ProteoStat” in cells after MG132 treatment (Fig. 7I∼K). In WT cells aggresomes exhibited clear staining by both p97 and Ub conjugate antibodies as in previous studies(*9*). However, in HUWE1Δ cells the association of both markers with the aggresome was significantly reduced, as demonstrated by the Pearson correlations with the aggresome channel. Overall, our results suggest that HUWE1 promotes the timely clearance of several forms of protein aggregates and this activity requires the segregase p97, which most likely recognizes the Ub conjugation mediated by HUWE1 on protein aggregates.

### HUWE1 mitigates the toxicity of protein aggregates

To understand the overfall cellular consequences of the action of HUWE1 on protein aggregation, we studied whether it had an effect on cellular morphology and viability. Although agDD aggregates are apparently benign in WT HEK293 cells(*60*), HUWE1Δ cells responded dramatically by rounding up and detaching from the surface six hours after shield-1 removal to trigger agDD aggregation (Fig. 8A∼C; Fig. S28A,B). A cell-death assay also reported a higher death rate for the cells that remained on plate, only in the presence of agDD aggregation in HUWE1Δ cells (Fig. S28C). A similar phenotype was also observed in the Huntington-disease model where HUWE1Δ led to a hypersensitivity to Htt(Q94) aggregate formation, cell rounding and detachment, as suggested by a dramatic reduction of cell density on surface after Htt(Q94) aggregation was induced (Fig. 8D; Fig. S29).

**Figure 8.**
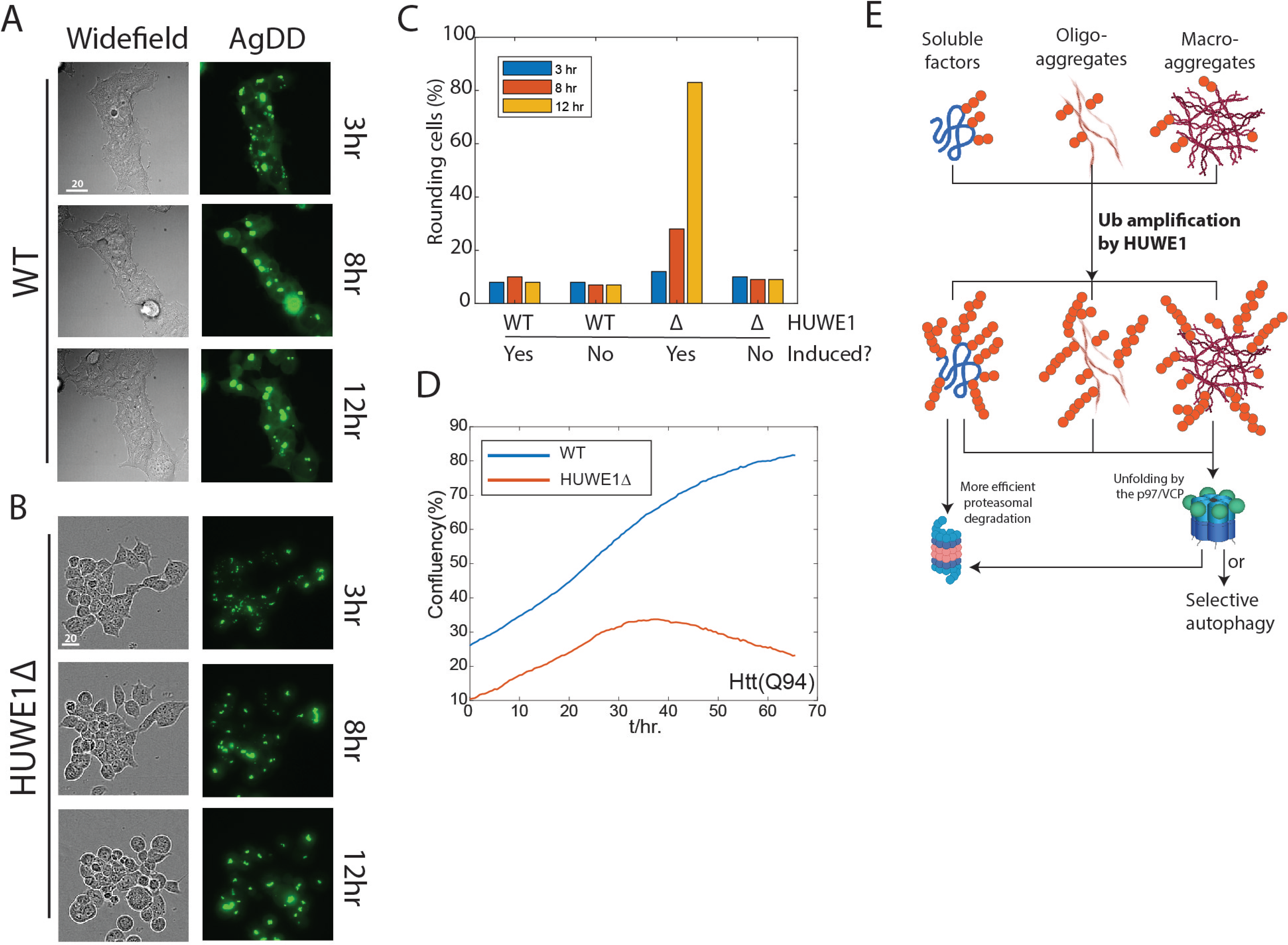
HUWE1 mitigates the cytotoxicity of protein aggregates. **A.** WT or HUWE1Δ (**B**) cells stably expressing agDD were induced to form aggregates and were imaged at indicated time points. Scale bar unit: μm. **C.** The percentage of rounding cells that remained attached to the surface in the experiment described in A,B. **D.** Expression of Htt(Q94) was induced at t=0 by adding doxycycline in stable WT or HUWE1Δ cell line. Cell confluency was monitored using an Incucyte imager. **E.** The proposed model for HUWE1-mediated Ub signal amplification and its biological functions.

## DISCUSSION

The eukaryotic protein degradation system encounters substrates with diverse structural features, stabilities, and different condensation states. Inefficient processing of these targets results in a variety of human diseases. In this study, we identified the HECT-family Ub ligase HUWE1 as a key regulator of cellular proteostasis through broadly facilitating the degradations of many soluble substrates and protein aggregates/condensates. These activities of HUWE1 regulate key biological processes, such as cell division, mediate the degradation of therapeutic targets and control the cellular response to protein aggregation.

### Ub amplification by HUWE1 generates a “try-hard-to-degrade” signal

The versatility of HUWE1 in facilitating the metabolism of proteins and aggregates depends on the exquisite features of a novel class of Ub ligase activity, UDL, associated with HUWE1, that mediates a previously-unrecognized molecular process – Ub signal amplification on protein targets (Fig. 8E). The ubiquitylation state generated by HUWE1 appears to constitute a “try-hard-to-degrade” signal that recruits the potent segregase p97/VCP to process difficult substrates such as aggregates and factors that lack features permitting efficient degradation by the proteasome, such as certain TPD targets. Remarkably, many kinases depend on HUWE1, to a similar extent as their depending on the p97/VCP, for targeted degradation (Fig. 5A,B), which strongly supports a close connection between the activities of HUWE1 and p97. HUWE1 knockout desensitizes a leukemia cell line (KBM-7) to killing by TPD drugs, Indisulam and CC-885, in a CRISPR screening (*66*), which may result from a compromised degradation of their targets.

### The HUWE1-p97 axis clears protein aggregates

The dynamic formation and clearance of protein aggregates/condensates play an important role in normal cell physiology and are associated with various diseases, such as neurodegeneration. The p97/VCP complexes are broadly involved in this process(*7, 13, 67*). However, the pathway that provides the Ub signal to recruit p97/VCP to these aggregates was not well understood. Targeting by p97 has been suggested to require a long Ub chain, although the exact requirement may vary with the substrate adaptors(*68*). UDL-mediated ubiquitylation is highly processive, which may be key to establishing the long Ub chains for targeting by the p97, especially in an intracellular milieu with active deubiquitylation. Besides the length of Ub chains, Ub-chain branching may also promote p97 recruitment, as HUWE1 can generate several different Ub linkages and forms branched chains (*69*).

Both the proteasome and autophagy may be involved in degrading the fragmented protein aggregates after HUWE1-p97. Their respective roles remain to be determined. Although we focused on the macroscopic aggregates in this study, HUWE1 also recognizes ubiquitylated, oligomeric aggregates which may be much more toxic to the cell than macroscopic aggregates(*70*). A buildup of smaller aggregates may underlie the toxicity of aggregation-prone proteins in the absence of HUWE1.

### HUWE1’s UDL activity is commonly involved in the degradation of short-lived proteins by the proteasome

HUWE1-mediated Ub amplification can directly stimulate proteasomal degradation. Short-lived proteins differ in their dependence on HUWE1 and a stronger effect was recorded for substrates with difficult features for processing by the proteasome. Although the direct *in vivo* equivalence of the Ub^L73F^ conjugates is still ambiguous, the proteasome degrades a substantial fraction of endogenous Ub together with substrates, especially when the proteasome-associated DUB Usp14/Ubp6 is compromised (*71, 72*). UDL activity may promote substrate-proteasome interaction and ensure efficient degradation under these conditions. Additionally, UDL might prioritize the degradation of factors that carry multiple Ub chains, such as the substrates of APC and SCF. This might be especially important when the capacity of the UPS becomes limited under stress or pathological conditions. Consistent with this notion, knocking out HUWE1 strongly increases the sensitivity of the HAP1 cell to a variety of mitotic inhibitors(*73*).

### Ub chain density on substrate is a separate dimension of the ubiquitin code, detected by HUWE1

HUWE1 does not amplify the ubiquitylation of all proteins but requires a high local-density of Ub chains on its substrates. Proteomic studies have identified more than 13,000 ubiquitylation sites on about 4000 ubiquitylation substrates in human cells, averaging about three sites per substrate(*74*). Our study supports the generality of HUWE1’s UDL activity in ubiquitylating and degrading endogenous Ub conjugates involved in diverse processes (Fig. 4C). Taken together, these results suggest that a wide variety of proteins may undergo amplification of their Ub signal by UDL(s), and that Ub amplification may be a general step in the ubiquitylation pathway. This study suggests the copy number of Ub chains on a substrate may be an important dimension of the “ubiquitin code”, involved in regulating the processes, such as cell division, that require efficient protein breakdown(*3*).

### HUWE1’s UDL represents a novel class of Ub ligase activity

Ub-protein ligases (E3s) often recognize substrates based on structural features or sequence motifs on substrates(*37*). For certain substrates, specialized Ub-chain elongation factors (E4s) are involved to collaborate with the E3 enzymes in establishing polyubiquitin chains on substrates(*37*). An E4 enzyme may recognize the Ub modification on its substrate or the Ub-substrate conjugate(*75*), but does not substitute for the E3 enzyme in ubiquitylation. Substrate recognition by HUWE1 resembles an E4, as it strictly depends on the substrate’s prior ubiquitylation by a substrate-specific E3, such as the APC. Instead of engaging in elongating the existing Ub chains, HUWE1 transfers Ubs directly to available sites on the substrate molecules, as does an E3. E4s often extend free Ub chains(*21, 38*). Unlike these factors, HUWE1 does not elongate free Ub chains at a physiological HUWE1 concentration. Therefore, HUWE1’s UDL activity integrates the features of both E3 and E4 enzymes and represents a novel class of Ub ligase activity. The mechanistic features of UDL are likely to be important for HUWE1’s substrate selectivity and for rapidly expanding the Ub modifications in, perhaps, a positive feedback fashion.

There are likely to be other factors containing UDL activity and features recognizing previous rounds of ubiquitylation. Our study supports this possibility (Fig. S5A). These new enzymatic activities, represented by HUWE1, greatly increase the substrate variants generated by modification by Ub and extend the regulatory potential of the ubiquitylation system.

## Acknowledgements

We are grateful for fundings from the National Institute of Health (R01 GM134064-01 to Y.L.; R01 CA214607 to E.S.F.; P30 AG062421 & R21 AG072516 to Y. S.) and the Mathers Foundation (grant to E.S.F.). We thank Dr. Yihong Ye at National Institute of Diabetes and Digestive and Kidney Diseases for sharing the HUWE1Δ and control cell lines. We want to thank Ms Meghan Van Orden for assistance in administration. We also want to thank Dr. Tim Mitchison, Dr. Daniel Finley, Dr. Marc Kirschner and Dr. Alfred Goldberg for valuable discussions and for commenting on the manuscript.

## Author contributions

Y. L. and M. Z. designed the project; M. H., M. Z. and L. C. purified the proteins; M. Z., L.C., R. F., Y. S., and C. Z. performed the study; G. B., R. E., M. K. and K.A.D. analyzed mass spec samples. Y. L., G. B., M. K., M. Z., L. C, Y. S. and R. F. analyzed the data. All authors participated in manuscript preparation.

## SUPPLEMENTARY FIGURE LEGENDS

**Figure S1. Ub^L73F^-conjugated cycB is degraded in cell extract but not by the purified 26S proteasome**. **A.** Ubiquitylation of HA-cycB-cpGFP or HA-cycB-EGFP by APC-UbcH10 in the presence of Ub^WT^ or Ub^L73F^. Samples were analyzed by anti-HA western blotting. **B.** CycB-cpGFP, unmodified or poly-ubiquitylated by recombinant APC-UbcH10 with the indicated Ub variant, was incubated with either purified human 26S proteasome (PTSM) or in asynchronized HeLa Cell extract. GFP fluorescence was detected by a plate reader.

**Figure S2. Degradation of Ub^L73F^-conjugates cycB substrates in cell extract depends on the proteasome and the Ub E1 activity in extract. A, B.** CycB-cpGFP conjugated with poly-Ub^WT^ or poly- Ub^L73F^ by recombinant APC-UbcH10 was incubated in asynchronous HeLa cell extract in the presence of a proteasome inhibitor (Bortezomib), an Ub E1 inhibitor (MLN7243) or DMSO. Fluorescent signal was detected on a plate reader. **C.** Polyubiquitylated cycB-EGFP or cycB-mNeonGreen(mNG) was incubated in HeLa cell extract, with or without the Ub E1 inhibitor (E1inh). The initial degradation rate of these substrates was calculated from the fluorescent signal that was detected on a plate reader.

**Figure S3. The L73F mutation on Ub does not affect substrate interaction with proteasome. A.** Schematic of the single-molecule binding assay (see methods)(*33*). **B.** Dwell time distribution of securin, conjugated with poly-wtUb or poly-Ub^L73F^ that is also fluorescently labeled, on immobilized 26S proteasome. The dwell time is registered as a function of the copy number of Ubs on a securin molecule. Error bars represent the standard error of dwell time.

**Figure S4. Further ubiquitylation of cycB-cpGFP-polyUb^L73F^ in extract is not due to any E3 or E4 activity associated with the APC, proteasome or p97/VCP. A∼C.** HeLa cell extract, whose APC, p97 or proteasome had been immunodepleted using their respective antibodies(methods), was incubated with HA-cycB-cpGFP that has been ubiquitylated with Ub^L73F^ by APC-UbcH10. The time-series samples were analyzed by anti-HA western blotting.

**Figure S5. Cell extract contains factors that can further ubiquitylate cycB-cpGFP-polyUb^L73F^. A.** Hela extract was fractioned by anion exchange (monoQ) chromatography. Each fraction was tested for ubiquitylating HA-cycB-cpGFP-polyUb^L73F^ in the presence of E2 UbcH5a. Fractions with highest activities (indicated) were grouped and fractionated again by size-exclusion (Superose 6) chromatography. The resulting fractions were tested likewise. **B.** E2 scanning. 34 E2s were tested for their activity in mediating the ubiquitylation of HA-cycB-cpGFP-polyUb^L73F^ by the active FPLC fraction from A.

**Figure S6. HUWE1 is responsible for the further ubiquitylation of cycB-cpGFP-polyUb^L73F^ in an active fraction and can react with wtUb-conjugated cycB with a similar efficiency. A.** HUWE1 was immunodepleted from the active fractions (grouped) in Fig. S5A. The depleted fraction or the anti- HUWE1 beads were tested for ubiquitylating HA-cycB-cpGFP-polyUb^L73F^. **B.** Ubiquitylation of HA-cycB-cpGFP-polyUb^L73F^ or HA-cycB-cpGFP-polyUb^WT^ by the gel-filtration fractions described in Fig. S1A.

**Figure S7. HUWE1’s UDL activity maintains high processivity at low nM concentrations. A.** Purification of recombinant HUWE1. Coomassie stain of SDS-PAGE. **B.** Titration of rHUWE1 concentration in ubiquitylating p^32^-labeled cycB-cpGFP-polyUb. Samples were analyzed by autoradiography. **C.** P^32^-labeled cycB-cpGFP was ubiquitylated by purified APC with E2 UbcH10 (left), or UbcH10+Ube2S (middle), or UbcH10 then Ube2S (right). **D.** P32-labeled geminin was ubiquitylated by the APC plus UbcH10, and then was tested for ubiquitylation by HUWE1 or HUWE1^UBDmu^. Samples were resolved by SDS-PAGE and autoradiography.

**Figure S8. HUWE1 does not react with free Ub chains. A.** K48-linked, Dy550-labeled Ub chains were purified and separated as described in methods. These chains were incubated with 30nM HUWE1 in a ubiquitylation reaction. Samples were separated by SDS electrophoresis and analyzed by a fluorescence imager (Bio-Rad). A standard Ub-chain formation reaction by the E2, E2-25K, involving these chains were used as a positive control (see methods). **B.** Preformed K48-linked Ub^L73F^ chains were incubated with HeLa extract. Ub is also biotinylated at the N-terminus. The samples were analyzed by anti-biotin western blotting which was intentionally overexposed to reveal the further ubiquitylation (if any).

**Figure S9. HUWE1 reaction forms different Ub linkages on substrates.** CycB-cpGFP-polyUb (p^32^ labeled) was ubiquitylated by rHUWE1 in the presence of indicated Ub variants. Samples were analyzed by autoradiography.

**Figure S10. The UBDs of HUWE1 mediate its UDL activity. A.** HUWE1^UBDmu^ is competent for ubiquitylating MCL1-1 and DDIT4. Ubiquitylation of purified MCL-1 and DDIT4 by HUWE1^UBDmu^. Samples were processed as in Figure 1C. **B.** WT HUWE1 or HUWE1 variants with indicated UBD mutations were tested for ubiquitylation of cycB-cpGFP-polyUb, at indicated concentrations. **C.** As in B, but using purified HUWE1 with indicated mutations(*35*).

**Figure S11. UDL stimulates proteasomal degradation but inhibits it at very Ub stoichiometry. A.** GFP fluorescence intensity in a RUDS containing the 26S proteasome, HUWE1, cycB-cpGFP- polyUb^L73F^(pre-ubiquitylated by APC-UbcH10) and either WT Ub or methylated Ub (see methods). The initial rate of fluorescence decay, normalized by the value in the absence of HUWE1, is presented on the right. Error bars represent the standard deviation of three replicates. **B.** As in A, but using cycB-EGFP- polyUb^WT^ as the substrate in a RUDS containing either WT Ub or methylated Ub. **C.** P^32^-labeled cycB- cpGFP was reacted with the APC in the presence of Ub^WT^ or Ub^L73F^ to form polyubiquitylation. The products were then briefly reacted with HUWE1 in the presence of Ub^WT^ or no Ub, and were quenched with an E1 inhibitor before the 26S proteasome was mixed in (t=0). The degradation rate of each ubiquitylated cycB species, separated into six zones (z1∼z6), was determined from a time series and presented on the right.

**Figure S12. Cell extract contains high deubiquitylating activity. A,B.** Deubiquitylation of HA-cycB- cpGFP-polyUb^WT^ and securin-mcherry-polyUb^WT^ in HeLa S3 extract. Substrates were first ubiquitylated by purified APC plus UbcH10 and then were incubated with the cell extract which had been treated with a proteasome inhibitor (Bortezomib) and an E1 inhibitor (MLN7243). 25uM DUB inhibitor Ub-vinyl sulfone was added as a control. Samples were analyzed by anti-HA and anti-securin western blotting. **C.** Raw data for figure 5A.

**Figure S13. Volcano plot of the fold change (FC) and the p values of proteins in HUWE1Δ vs. WT cells.** The whole cell lysate of HUWE1Δ and WT HEK293 cells was analyzed by quantitative mass spectrometry (Fig. 4C). The FC (X) of identified proteins was plotted with the p values (Y). Known APC and SCF substrates(*76*) were highlighted with green and red asterisks.

**Figure S14. HUWE1 knockout decreased the accumulation of K48-chain conjugates by proteasome inhibition, without affecting the overall ubiquitylation level. A.** WT HEK293 or HUWE1Δ cells were treated with MG132 or DMSO for 8hrs. The cells were lysed, and the soluble fraction(SN) and the pellet were separated by centrifugation and were probed by anti-Ub (**A**) and anti-K48 chain (**B**) western blotting. GAPDH was used as a loading control.

**Figure S15. Gene Ontology(GO) analysis of significantly up or down regulated proteins in HUWE1Δ cells.** Protein abundance in WT and HUWE1Δ HEK293 cells were compared using label-free quantitative mass spectrometry (methods). Proteins that are significantly (P<0.05) up or down regulated in HUWE1Δ (log2(fold change) >1 or < -1) are identified and analyzed by shinyGO (*44*). The gene names associated with each GO term can be found in Table S2.

**Figure S16. HUWE1 knockout does not significantly affect most transcripts’ levels.** The fold changes of ∼8600 transcripts were extracted from a dataset generated by Perturb-seq(*45*) in either K562 and RPE1 cells, and the distribution is plotted. There are 11 and 29 transcripts whose levels increase by more than 2X in K562 and RPE1 cells respectively.

**Figure S17. Expression of RFP-HUWE1. A.** Ub-R-YFP was transiently expressed in WT HEK293 or *HUWE1Δ.* Cycloheximide was added at t=0 to stop protein synthesis, and the cells were imaged once every 7 minutes. The averaged degradation kinetics of these reporters were calculated from single-cell traces (Ub-R-YFP N=31). The shaped area represents the standard deviation. **B.** Anti-HUWE1 and Anti-streptag western blotting of cells expressing strepTag-RFP-HUWE1 or strepTag-RFP-HUWE1^UBDmu^. **C.** Sample images of both securin-YFP and RFP-HUWE1 channels for WT or *HUWE1Δ* transiently transfected with these constructs.

**Figure S18. Ubiquitylation of dendra substrate.** Recombinant SUMO-dendra2 was purified and photo-cleaved as in a previous study(*48*). After de-sumoylation, the substrate was incubated with 1x or 2x Ubr1 (E3), and was analyzed by anti-dendra western blotting.

**Figure S19. UDL stimulates p97-mediated unfolding of a tetramer substrate. A.** HA-tagged synthetic substrate Ub(3)-VASP-cpGFP was ubiquitylated by rHUWE1 as in Figure 3A, and then was incubated with recombinant p97-UFD1-NPL4 complex in the presence of 1uM GroEL(D87K). The fluorescence intensity was monitored using a plate reader. **B.** the same substrate, after HUWE1 reaction, was incubated with purified human 26S proteasome.

**Figure S20. Characterization of HUWE1Δ HEK293 cells. A.** Growth curve of HUWE1Δ cells, determined by an Incucyte imager. **B.** DNA content of WT and HUWE1Δ, determined by FACS and Propidium Iodide staining.

**Figure S21. AgDD forms an aggresome-like aggregates upon induction. A.** Kymograph images of agDD(GFP)-expressing HEK293 cells. Shield-1 was removed at t=0 to induce aggregation. **B.** Fluorescence image of fixed HEK293 cells containing agDD aggregates. **C.** Cells were treated as in A, but with colchicine or Nexturastat A (a HDAC6 inhibitor) added right after shield-1 removal. Images were taken 4 or 8hrs after the induction.

**Figure S22. Clearance of agDD aggregates in cells. A.** Example images of agDD-expressing cells exhibiting three different phenotypes: no aggregates, unstable aggregates that undergo spontaneous clearance, stable aggregates without obvious clearance. Shield-1 was removed at t=0 to induce aggregation. **B.** AgDD levels, as determined by the total fluorescence intensity in individual cells exhibiting the three phenotypes. **C.** The percentage of cells that have spontaneously cleared agDD aggregates at 4 or 8hrs after shield-1 removal, in the presence of indicated compounds. MLN7243: Ub E1 inhibitor; NMS-873: p97/VCP inhibitor; Chloroquine: autophagy inhibitor. **D.** As in the C, but cycloheximide (CHX) was added 30 minutes after shield-1 removal.

**Figure S23.** Raw data for figure 7B.

**Figure S24. HUWE1 mediates the clearance of synphilin aggresome and promotes the degradation of over-expressed synphilin. A.** Synphilin aggresomes were induced in WT or HUWE1Δ cells stably expressing synphilin-GFP by incubating the cells with MG132 for 8hrs (*64*). MG132 was washed off at t=0 to initiate the clearance of synphilin aggresome. The clearance process was monitored by timelapse microscopy (methods). Error bars represent the standard deviation of three replicates. **B.** Sample images**. C.** HUWE1 (WT, HUWE1^CS^ or HUWE1^UBDmu^) and synphilin-GFP plasmids were co-transfected into HEK293 cells. 1d or 2d after transfection, cells were harvested and the whole-cell lysate was analyzed by western blotting. Two anti-GFP antibodies were used to quantify the synphilin-GFP levels. Numbers are quantification of the relative synphilin concentrations (an average of the two anti-GFP blots).

**Figure S25. Clearance of heat-shocked induced stress granules in WT or HUWE1Δ.** Experiment and analysis was performed as described in Figure 7E,F, except that stress granules were induced by heat shock at 43 degree for 60 minutes, and cells were moved to 37 degree for recovery. Error bars represent the standard deviation of three replicates.

**Figure S26. Colocalization of RFP-HUWE1 with agDD aggregates requires its Ub-binding domains.** Experiment was performed as described in Figure 7G, but using either RFP-HUWE1^CS^ or RFP- HUWE1^UBDmu^.

**Figure S27. Expression and distribution of HUWE1 in WT and THY-Tau22 mice. A.** Confocal images showed that HUWE1 was highly expressed in cortical neurons in both WT mice and THY-Tau22 mutant mice. **B.** Structured-Illumination-Microscopy (SIM) demonstrated neuronal colocalization of HUWE1 and Tau in the symptomatic THY-Tau22 mice. **C.** Quantification of colocalization using Manders’ overlap coefficient showed that ∼50% Tau is associated with HUWE1 in early-stage tau aggregates, ∼ 75% in mid-stage tau aggregates, and 0% in late-stage tau aggregates.

**Figure S28. HUWE1 controls the cytotoxicity of agDD aggregates (controls). A, B.** as described in Figure 8A,B, but without inducing agDD aggregates. **C.** As described in Figure 8A,B. The viability of the cells were determined using CytoTox-Glo(Promega).

**Figure S29. HUWE1 controls the cytotoxicity of Htt(Q94) aggregates.** Sample images for the experiment described in fig. 8D. The percentage of cells that have rounded at indicate time points, in the population with or without a clear Htt(Q94) aggregate (instead showing diffusive Htt-CFP signal), was counted.

## SUPPLEMENTARY TABLE LEGENDS

**Table S1. Identification of factors for further ubiquitylation of polyubiquitylated cycB**. Asynchronous Hela S3 extract was first fractionated by anion exchange chromatography. The eluent at around 350mM NaCl was collected and fractionated again by size-exclusion chromatography into 10 fractions. Protein abundance across these fractions was detected by 10-plex TMT mass spectrometry. Proteins were then clustered according to their elution profiles, and were compared with the ubiquitylation activity profile of these size-exclusion fractions (tab: Western_Blot) using polyubiquitylated cycB-cpGFP as the substrate.

**Table S2. Global proteomics comparing WT and HUWE1Δ HEK cells** (tab: KO vs WT). GO term analysis was performed using shinyGO on proteins whose levels went up or down by more than two folds in HUWE1Δ cells.

**Table S3. Proteomic profiling of WT and HUWE1Δ cells treated with different degrader compounds.** WT or HUWE1**Δ** cells were treated with each compound at indicated concentration (tab: description) for 5 hours, in three replicates. Cell pellets were analyzed by quantitative mass spectrometry (see methods).

## SUPPLEMENTARY MOVIE LEGENDS

**Movie S1. Induction of agDD aggregation.** HEK293 cells stably expressing agDD(GFP) were grown in the presence of shield-1. FKBP was added at the beginning to induce the aggregation of agDD.

## Materials and Methods

### Resource table

| Reagent or Resource | Source | Identifier |
| --- | --- | --- |
| **Plasmids** |  |  |
| pT7-His6-Cys-Ubiquitin(Human) | Lu et al., Science 2015 | pLM40 |
| pT7-His6-Cys-Ubiquitin(L73F) | This study | pLM202 |
| pT7-His6-HA-cyclinB(1~88)-<br>cpGFP | This study | pLM460 |
| pT7-His6-HA-cyclinB(1~88)-<br>mNeonGreen | This study | pLM444 |
| pT7-His6-HA-cyclinB(1~88)-<br>EGFP | This study | pLM446 |
| pT7-His6-securin | Lu et al., Science 2015 | pLM74 |
| pT7-His6-Geminin | Lu et al., Science 2015 | pLM75 |
| pT7-His6-HA-cyclinB(1~88)-<br>[TEV]-myc-mNeonGreen | This study | pLM468 |
| pT7-His6-HA-Ub(1)-VASP-cpGFP-tail | This study | pLM510 |
| pT7-His6-HA-Ub(3)-VASP-cpGFP-tail | This study | pLM525 |
| pT7-HA-Ub(1)-mNeonGreen-His6 | This study | pLM563 |
| pT7-HA-Ub(2)-mNeonGreen-His6 | This study | pLM564 |
| pT7-HA-Ub(3)-mNeonGreen-His6 | This study | pLM565 |
| pT7-SUMO-Dendra2 | Ji Z. et al., Mol Cell, 2022 |  |
| pT7-His6-HA-cyclinB(1~88)-<br>mEOS | This study | pLM454 |
| pTetOn-HA-streptag-KO2-HUWE1 | This study | pLM551 |
| pTetOn-HA-streptag-KO2-HUWE1_CS | This study | pLM552 |
| pTetOn-HA-streptag-KO2-HUWE1_UBDmu<br>(M1328A/F1330A/<br>A1376G/S1382A/L4976A/P2977A) | This study | pLM554 |
| pT7-his6-securin-mcherry | A gift from Qiong Yang | Psec-mcherry |
| pBOB-EF1-FastFUCCI, lentiviral | Koh SB et al., J. Cell Sci.<br>2017 | Addgene: 86849 |
| pCMV-EGFP-G3BP1, lentiviral | This study | pLM580 |
| pTetOn-Sph1(synphilin)-EGFP,<br>lentiviral | This study | pLM579 |
| pTetOn-HttQ94-CFP, lentiviral | This study | pLM582 |
| pCMV-AgDD, lentiviral | Miyazaki Y. et al., Nature<br>Comm. 2016 | pF57; addgene 78289 |
| pCMV-Ub-R-YFP | Menendez-Benito et al.,<br>Mol. Genet. 2005 | pF73; addgene 11948 |
| pCMV-securin-YFP | Hagting et al., J. Cell Biol.<br>2002 | Addgene 39847 |
| pTetOn-His6-HA-UBA52; pTetOn-His6-HA-RPS27A | This study | pLM403 |
| Constructs for purifying<br>recombinant human p97, NPL4,<br>UFD1 | Ji Z. et al., Mol Cell, 2022 |  |
| **Purified proteins** |  |  |
| Various substrates, APC, HUWE1,<br>26S proteasome | This study | See methods |
| Ube2S, UbcH10, UbcH5a,<br>Usp2(CD), Human E1 | Boston Biochem |  |
| Methylated Ubiquitin | Boston Biochem |  |
| Single Lysine Ubiquitin (K48, K63,<br>K11 or K6) | Boston Biochem |  |
| Di-ubiquitin chains (K48, K63 or<br>K29) | Boston Biochem |  |
| Tetra-ubiquitin chains (K48 or K63) | Boston Biochem |  |
| HUWE1 patient mutants (D169-189, H669Q, F3194S, R4187C) | Hunkeler M et al., Mol Cell. 2021 |  |
| Tom1 | This study |  |
| Recombinant human p97, Ufd1, Npl4, Ubr1, Ubc4 | This study; Ji Z. et al., Mol Cell, 2022 |  |
| GroEL(D87K) | Lu et al., Mol. Cell. 2017 |  |
| **Cell lines** |  |  |
| HEK293 |  |  |
| HEK293, TetR | GenTarget Inc |  |
| HEK293, pTetOn-His6-HA-Ubiquitin(UBA52, RPS27A) | This study |  |
| HEK293, FastFUCCI | This study |  |
| HEK293, HUWE1Δ, clone 1 and clone 2 | Xu Y. et al., Cell Discov. 2016 |  |
| HUWE1Δ, FastFUCCI | This study |  |
| pCMV-GFP-G3BP1 (WT HEK293 or HUWE1Δ) | This study |  |
| pTetOn-Sph1-GFP (WT HEK293 or HUWE1Δ) | This study |  |
| pTetOn-HttQ94-CFP (WT HEK293 or HUWE1Δ) | This study |  |
| pCMV-agDD, HUWE1Δ, clone 1, clone 2 | This study |  |
| pCMV-agDD, WT HEK293 | This study |  |
| **Key chemicals, antibodies and kits** |  |  |
| TAK-243 (MLN7243) | Cayman |  |
| NMS-873 | Tocris |  |
| E-7820 | Tocris |  |
| Indisulam | Tocris |  |
| CC-885 | Cayman |  |
| SK-3-91 | Donovan KA et al., *Cell* 2020 |  |
| DB-0646 | Donovan KA et al., *Cell* 2020 |  |
| Shield-1 | Takara Bio |  |
| Anti-ubiquitin conjugates (FK2) antibody | Millipore | FK2 |
| Anti-ubiquitin antibody | Cell signaling | E4I2J |
| Anti-K48 chain antibody | BostonBiochem | A-101 |
| Anti-p97 antibody | Proteintech | 10736-1-AP |
| Anti-huwe1 antibody 1 | Bethyl | A300-486A |
| Anti-huwe1 antibody 2 | Bethyl | A300-700A |
| Anti-GFP antibody | Proteintech | 3H9 |
| Anti-dendra antibody | Thermo Scientific | OTI1G6 |
| Anti-MCL1 antibody | Cell signaling | Cat#5453 |
| Anti-DDIT4 antibody | Cell signaling | Cat#2516 |
| Anti-securin antibody | MBL | K0090-3 |
| Anti-tau aggregates (AT8) antibody | Thermo Scientific | MN1020 |
| Anti-Tubb3 | Biolegend | 657406 |
| Anti-streptag | Thermo Scientific | PA5-119772 |
| E2 scan kit, version 2 | Ubiquigent |  |
| CytoTox-Glo™ Cytotoxicity Assay | Promega |  |
| ProteoStat Aggresome Detection kit | Enzo Life Sciences |  |
| **Software** |  |  |
| Fuji (v.1.53), plugins: MiToBo,<br>Coloc 2 |  |  |
| MATLAB | Mathworks |  |
| Biorender |  |  |
| Cellprofiler |  |  |
| Micromanager2 |  |  |

### Methods

#### 1. Construction of plasmids

Plasmids for expressing cys-Ub/Ub^L73F^, HA-cyclinB(1∼88)-cpGFP/mNeonGreen/EGFP/mEOS, HA- cyclinB(1∼88)-[TEV]-myc-mNeonGreen, Ub(1,3)-VASP-cpGFP and Ub(1,2,3)-mNeonGreen-His6 were constructed in a pET28 vector under a T7 promoter. The VASP domain was based on the VASP-4M sequence(*40*). For Ub(1,3)-VASP-cpGFP-tail constructs, a 35-AA peptide from S. cerevisiae cytochromeB2 was added as the proteasome initiation sequence (*29*). A PKA site (RRASV) was added to both the N- and C-termini of substrates used for radioactive labeling.

Plasmids expressing RFP-HUWE1/HUWE1^CA^/HUWE1^UBDmu^ were constructed in a pBluescript vector (addgene: 72835) containing a tetracycline-inducible promoter (TetOn). A KO2(RFP) sequence, followed by a Gateway destination cassette, was inserted after the TetOn promoter. Human HUWE1 sequences were subcloned using Gateway reaction. HUWE1^UBDmu^ contains the following mutations: UBA(M1328A, F1330A), UIM(A1376G, S1382A), UBM(L4976A, P2977A). HUWE1^CS^ contains the following mutation: C4341S. Plasmids used for expressing recombinant HUWE1 have been described in a previous study.

For the plasmid expressing synphilin-GFP or Htt(exon1)Q94-CFP, the respective coding sequence was inserted into a lentiviral vector under a TetOn promoter. EGFP-G3BP1 was cloned under a CMV promoter in a lentiviral vector. The His6-HA-Ub(UBA52, RPS27A) sequence was cloned into a pCDNA5/TO vector under the control of a TetOn promoter.

#### 2. Construction of cell lines

The cell line overexpressing His6-HA-Ub was constructed by stably integrating the expression plasmid into a HEK293(TetR) cell line. Stable clones were selected by isolating single colonies.

Cell lines stably expressing FUCCI markers were constructed by integrating fastFUCCI (addgene: 86849) into WT HEK293 or HUWE1Δ cell lines using lentivirus and selected in the presence of puromycin for one week.

Cell lines stably expressing agDD or GFP-G3BP1 were constructed by integrating the expression constructs into WT or HUWE1Δ through lentivirus, followed by puromycin selection. 1uM shield-1 was added to agDD culture to prevent aggregation.

Cell lines expressing GFP-synphilin or HttQ94-CFP under TetOn promoter were constructed by integrating the expression constructs into WT or HUWE1Δ through lentivirus, following by puromycin selection.

#### 3. Protein purification and labeling

Recombinant Ub and substrates (Ub, UbL73F, cycB-FPs, securin, geminin, Ub(1,2,3)-VASP-cpGFP etc) were purified from E. coli cells through a poly-His tag and Ni-NTA agarose. All proteins were buffer-exchanged and stored in UBAB buffer (25mM Tris-HCL[pH7.5], 50mM NaCl, 5mM MgCl2). Protein concentrations were determined using Bio-Rad protein assay; ubiquitin concentration was determined by UV A280 absorption.

To fluorescently label Ub, a cysteine residue was placed between the N-terminal His tag and Ub. Ub or UbL73F was further purified using cation exchange chromatography and was labeled with Dylight550- maleimide. After removing unreacted dyes, labeled Ub was subjected to anion exchange chromatography to separate labeled and unlabeled species. Finally, the N-terminal His tag was cleaved off using thrombin and thrombin was later removed using aminobenzamidine beads.

Anti-20S antibody (MCP21) was biotinylated using biotin-NHS, and was purified using a desalting column.

To biotinylate UbL73F, His6-cys-UbL73F was reacted with biotin-maleimide and was coupled to Ni-NTA beads to remove unreacted biotin. Labeled UbL73F was then cleaved off the beads using thrombin, followed by cation exchange chromatography.

Recombinant human MCL-1-ΔTM and DDIT4 were purified from sf9 insect cells via a Twin-Strep tag as previously described(*77*).

Purification of recombinant human p97, NPL4, UFD1 and yeast Ubr1 were performed as described in a previous study (*48*).

Radioactive P32-ATP was used to label substrates with a PKA site at the N-terminus for in vitro ubiquitylation or degradation assays.

#### 4. Purification of recombinant human APC-Cdh1

Purification of recombinant APC-Cdh1 from insect cells has been described elsewhere(*31*). Briefly, viruses expressing 14 APC components were generated by transfecting Sf9 insect cells with the recombinant baculoviral genome based on a biGBac system using Fugene 6 reagent. Amplified viruses were added to HighFive insect cell culture for protein expression.

APC was expressed with a Twin-Step(II)-tag on the C-terminus of APC4, and was isolated from cell lysate using Strep-Tactin sepharose, and then was polished by ion exchange chromatography and gel filtration. Myc-6xHis-Cdh1 was purified from sf9 insect cells using Ni2+ agarose and polished by gel filtration (Superdex 200) chromatography.

#### 5. Purification of endogenous or recombinant HUWE1

Endogenous HUWE1 was isolated from HeLa S3 cells. A 2L spinner culture of asynchronous HeLa S3 cells was collected and homogenized using nitrogen cavitation in swelling buffer (20mM Tris-HCl [pH 7.5], 5mM KCl, 1.5mM MgCl2, 1mM DTT, 1x protease-inhibitor tablet(Roche)). The S100 extract was prepared by centrifugation at 10,000g for 30 minutes and again at 100,000g for 60 minutes at 4C. The S100 extract was then equilibrated with buffer A (20mM Bis-Tris[pH7.0], 50mM NaCl, 2mM MgCl2, 1mM DTT, 10% glycerol) and separated on a monoQ column using a linear gradient from 50mM to 600mM NaCl. HUWE1 was eluted at approximately 350mM NaCl. Two fractions containing the highest HUWE1 activity were combined and further fractionated on a superose 6 column in buffer C (15mM Hepes[pH7.5], 50mM NaCl, 2mM MgCl2, 10% glycerol, 1mM DTT). The active fractions were combined and further separated on a Phenyl Sepharose column equilibrated with buffer D (20mM Hepes [pH7.5], 100mM NaCl, 5mM MgCl2, 1mM DTT) + 10% ammonium sulfate, and eluted with a linear gradient from buffer D + 5% ammonium sulfate to buffer D.

Recombinant human HUWE1 and *S. cerevisiae* Tom1 was purified from Expi293 cells via a N-terminal twin-strep tag(*35*). Briefly, plasmids carrying HUWE1 (or Tom1) or mutants were transiently transfected into Expi293 cells according to the manufacture’s instruction. Cells were harvested after 72 hours post transfection and lysed by sonication in lysis buffer (50 mM HEPES/KOH [pH 7.4], 200 mM NaCl, 2 mM MgCl2, 5% glycerol) supplemented with protease inhibitors. The lysate was cleared by ultracentrifugation (45 min, 120,000 g) and incubated with strep-tactin sepharose. Beads were washed with lysis buffer and protein was eluted with 5 column volumes of lysis buffer supplemented with 50mM biotin. The protein was concentrated using centrifugal concentrators and polished by size exclusion chromatography (superose 6) in gel filtration buffer (30 mM HEPES/KOH [pH 7.4], 150 mM NaCl, 2 mM TCEP).

#### 6. Purification of human 26S proteasome

Human proteasomes were affinity-purified on a large scale from a stable HEK293 cell line harboring HTBH tagged hRPN11(a gift from L.Huang). The cells were Dounce-homogenized in lysis buffer (50mM NaH2PO4 [pH7.5], 100mM NaCl, 10% glycerol, 5mM MgCl2, 0.5% NP-40, 5mM ATP and 1mM DTT) containing protease inhibitors. Lysates were cleared, then incubated with NeutrAvidin agarose resin overnight at 4°C. The beads were then washed with excess lysis buffer followed by the wash buffer (50mMTris-HCl[pH7.5], 1mM MgCl2 and 1mMATP). Usp14 was removed from proteasomes using wash buffer+100mM NaCl for 30 min. 26S proteasomes were eluted from the beads by cleavage, using TEV protease (Invitrogen).

#### 7. Purification of endogenous ubiquitin conjugates

The HEK293 cell line stably expressing His6-HA-Ub under TetOn promoter was cultured in 10 x 150mm dishes. 1μg/ml doxycycline was added to the culture for 3 days before harvesting to induce His6- HA-Ub expression. 500nM bortezomib was added to the culture 1 hour before harvesting. Cell pellet was dounce-homogenized in lysis buffer in lysis buffer (50mM NaH2PO4 [pH7.5], 100mM NaCl, 10% glycerol, 5mM MgCl2, 0.5% NP-40). The lysate was then centrifuged at 30,000g for 30 minutes at 4C. The supernatant was incubated with Ni-NTA agarose and the agarose was washed with 60x column volume of wash buffer (25mM Tris-HCL [pH7.5], 100mM NaCl, 10% glycerol, 5mM MgCl2) + 20mM imidazole and eluted with wash buffer + 250mM imidazole. The eluent was buffer exchanged into wash buffer +1mM DTT and concentrated using ultrafiltration (Amicon, 10kD).

#### 8. Synthesis, purification and methylation of ubiquitin chains

Synthesis of K48-linked UbL73F chains: 500μM UbL73F, 50μM biotin-UbL73F, 100nM E1, 20μM E2-25K was incubated overnight at 37C in buffer (50mM Tris-HCL, [pH7.5], 10mM MgCl2, 0.4mM DTT, 10mM ATP, 0.1mM PMSF). The reaction was then separated using cation exchange chromatography (monoS) as previously described(*78*).

Synthesis of K48-linked fluorescent Ub chains: 500μM wtUb, 50μM Dy550-Ub, 100nM E1, 20μM E2-25K was incubated overnight at 37C in a buffer (50mM Tris-HCL, [pH7.5], 10mM MgCl2, 0.4mM DTT, 10mM ATP, 0.1mM PMSF). The reaction was then separated using gel filtration (superdex 200) in TBS buffer (20mM Tris-HCL[pH 7.5], 150mM NaCl).

For preparing methylated Ub chains, chain samples were buffer-exchanged into (50mM Hepes-Na[pH 7.5], 150mM NaCl, 2% glycerol). Dimethylamine borane and formaldehyde were added to the sample to final concentrations of 20mM and 40mM respectively. After 2-hour incubation on ice, another 20mM dimethylamine borane and 40mM formaldehyde were again added, bringing final concentrations to 40mM and 80mM respectively, and incubation was continued for 16 hours on ice. Methylated proteins were buffer-exchanged to TBS buffer.

#### 9. Preparation and fractionation of cell extract

A 2L spinner culture of asynchronous HeLa S3 cells was collected and homogenized using nitrogen cavitation in swelling buffer (20mM Tris-HCl[pH 7.5], 5mM KCl, 1.5mM MgCl2, 1mM DTT, 1x protease-inhibitor tablet). The S100 extract was prepared by centrifugation at 10,000g for 30 minutes and again at 100,000g for 60 minutes at 4C. The S100 extract was then equilibrated with buffer A (20mM Bis-Tris[pH7.0], 50mM NaCl, 2mM MgCl2, 1mM DTT, 0.2mM ATP, 10% glycerol) and separated on a monoQ column using a linear gradient from 50mM to 600mM NaCl. Each fraction was collected, concentrated and buffer exchange into gel filtration buffer (15mM Hepes[pH7.5], 50mM NaCl, 2mM MgCl2, 10% glycerol, 1mM DTT, 0.2mM ATP) using Amicon filters. Selected fractions from anion exchange were further separated by gel filtration on a superose 6 column in gel filtration buffer. Fractions were then concentrated using Amicon filters.

#### 10. Quantitative mass spectrometry and data analysis

Wild type and HUWE1Δ HEK cells were treated with DMSO or small molecule degraders for 5 hrs. Cells were harvested by centrifugation and washed with phosphate buffered saline (PBS) before snap freezing in liquid nitrogen. Concentrations used: SK-3-91, 1μM; DB-0646, 1μM; Indisulam, 10μM; CC- 885, 10nM; E-7820, 10μM.

Cells were lysed by addition of lysis buffer (8 M Urea, 50 mM NaCl, 50 mM 4-(2- hydroxyethyl)-1-piperazineethanesulfonic acid (EPPS) pH 8.5, Protease and Phosphatase inhibitors) and homogenization by bead beating (BioSpec) for three repeats of 30 seconds at 2400. Bradford assay was used to determine the final protein concentration in the clarified cell lysate. 50 µg of protein for each sample was reduced, alkylated, tryptic digested and processed as previously described(*79*). Desalted tryptic peptides were dried in a vacuum-centrifuged and reconstituted in 0.1% formic acid for LC-MS analysis.

Data were collected using a TimsTOF Pro2 (Bruker Daltonics, Bremen, Germany) coupled to a nanoElute LC pump (Bruker Daltonics, Bremen, Germany) via a CaptiveSpray nano-electrospray source. Peptides were separated on a reversed-phase C18 column (25 cm x 75 µm ID, 1.6 µM, IonOpticks, Australia) containing an integrated captive spray emitter. Peptides were separated using a 50 min gradient of 2 - 30% buffer B (acetonitrile in 0.1% formic acid) with a flow rate of 250 nL/min and column temperature maintained at 50 °C.

DDA was performed in Parallel Accumulation-Serial Fragmentation (PASEF) mode to determine effective ion mobility windows for downstream diaPASEF data collection (*80*). The ddaPASEF parameters included: 100% duty cycle using accumulation and ramp times of 50 ms each, 1 TIMS-MS scan and 10 PASEF ramps per acquisition cycle. The TIMS-MS survey scan was acquired between 100 – 1700 m/z and 1/k0 of 0.7 - 1.3 V.s/cm2. Precursors with 1 – 5 charges were selected and those that reached an intensity threshold of 20,000 arbitrary units were actively excluded for 0.4 min. The quadrupole isolation width was set to 2 m/z for m/z <700 and 3 m/z for m/z >800, with the m/z between 700-800 m/z being interpolated linearly. The TIMS elution voltages were calibrated linearly with three points (Agilent ESI-L Tuning Mix Ions; 622, 922, 1,222 m/z) to determine the reduced ion mobility coefficients (1/K0). To perform diaPASEF, the precursor distribution in the DDA m/z-ion mobility plane was used to design an acquisition scheme for DIA data collection which included two windows in each 50 ms diaPASEF scan. Data was acquired using sixteen of these 25 Da precursor double window scans (creating 32 windows) which covered the diagonal scan line for doubly and triply charged precursors, with singly charged precursors able to be excluded by their position in the m/z-ion mobility plane. These precursor isolation windows were defined between 400 - 1200 m/z and 1/k0 of 0.7 - 1.3 V.s/cm2.

##### LC-MS data analysis

The diaPASEF raw file processing and controlling peptide and protein level false discovery rates, assembling proteins from peptides, and protein quantification from peptides was performed by analysis in DIA-NN 1.8 using targeted cell line specific spectral libraries generated by searching offline fractionated DDApasef data against a Swissprot human database (January 2021) (*81*). Library free mode performs an in silico digestion of a given protein sequence database alongside deep learning-based predictions to extract the DIA precursor data into a collection of MS2 spectra. The search results are then used to generate a spectral library which is then employed for the targeted analysis of the DIA data searched against a Swissprot human database (January 2021). Database search criteria largely followed the default settings for directDIA including: tryptic with two missed cleavages, carbomidomethylation of cysteine, and oxidation of methionine and precursor Q-value (FDR) cut-off of 0.01. Precursor quantification strategy was set to Robust LC (high accuracy) with RT-dependent cross run normalization. Proteins with missing values in any of the treatments and with poor quality data were excluded from further analysis (mean number of precursors used for quantification <2) and proteins with missing values were imputed by random selection from a Gaussian distribution either with a mean of the non-missing values for that treatment group or with a mean equal to the median of the background (in cases when all values for a treatment group are missing). Protein abundances were scaled using in-house scripts in the R framework

(R Development Core Team, 2014) and statistical analysis was carried out using the limma package within the R framework (*82*).

#### 11. In vitro ubiquitylation reactions

##### a. APC-mediated ubiquitylation

The APC ubiquitylation reactions were carried out in UBAB buffer + 0.5mM DTT and contain 30nM recombinant APC, 1μM recombinant Cdh1, 50nM E1, 1μM UbcH10, an energy regenerating system, and either ubiquitin or ubiquitin chains: 50μM ubiquitin; 50μM methylated ubiquitin, 10μM methylated K48- tetraubiquitin chain; 10μM methylated K63-tetraubiquitin chain; 20μM K48-diubiquitin chain; 20μM K29-diubiquitin chain; 20μM K63-diubiquitin chain. Reactions were incubated at 30°C.

To compare APC’s and HUWE1’s reaction processivity, the same concentration of 200nM substrate was used. To prepare ubiquitylated substrates for HUWE1 reaction, 50nM APC and 1μM substrate was used and the reaction was incubated for 90 minutes at 30C. The ubiquitylated substrates were then isolated on Ni-NTA magnetic beads, eluted and buffer exchanged into UBAB buffer using Amicon filters.

For reaction involving Ube2S, 100nM Ube2S was added together with 1μM UbcH10.

For PKA-labeled substrates, calyculin A, a broad-spectrum phosphatase inhibitor, was added at 10μg/ml. Results were quantified using a phosphorImager.

All samples for SDS electrophoresis in this study were prepared in Laemmli sample buffer supplemented with 50mM DTT and 6M Urea.

##### b. HUWE1-mediated ubiquitylation

The HUWE1 ubiquitylation reactions were carried out in UBAB buffer + 0.5mM DTT and contain 30nM(unless indicated) purified endogenous or recombinant HUWE1, 50nM E1, 1μM UbcH5a, an energy regenerating system, 50μM ubiquitin (or 50μM methylated ubiquitin), 200nM substrate (unmodified or preubiquitylated). Reactions were incubated at 30C. The 500kD molecular weight marker in figure 1B is based on HUWE1 detected by western blot.

For the reaction involving fluorescent Ub chains, 200nM purified Dy550-Ub chains were used as substrates. For the reaction involving endogenous Ub conjugates, a final concentration of 0.1mg/ml Ub conjugates were added to HUWE1 reactions.

For the reaction involving aggregates (Fig. 3C,D), 100nM rHUWE1 and 1μM substrate were used; 1ul Ni-NTA dynabeads was added to 40ul reaction to crosslink the substrate. Non-his6 tagged E1 and E2s are used in this reaction to avoid interference. The reaction was incubated at 30C with shaking at 900rpm.

#### 12. *In extract* ubiquitylation and deubiquitylation reactions

5ul fractionated cell extract was added to a 15ul reaction (total volume) containing 50nM E1, 1μM UbcH5a, an energy regenerating system, 50μM Ub or UbL73F, 100nM substrate, in UBAB buffer + 0.5mM DTT. Reactions were incubated at 30C and results were analyzed by western blotting.

To deplete factors from cell extract (HUWE1, APC, p97/VCP), the antibody (anti-HUWE1, anti- Cdc27, anti-p97) was conjugated to proteinA/G agarose. The beads were then incubated with cell extract for 90 minutes at 4C and depleted afterwards by centrifugation.

For testing deubiquitylation, asynchronous HeLa S3 extract was prepared as described above. The extract was treated 50μM bortezomib and 40μM MLN7243 at room temperature for 15 minutes. HA-cycB-cpGFP or securin-mcherry, which had been ubiquitylated by purified APC plus UbcH10, was then incubated with the treated extract. Samples were analyzed by anti-HA or anti-securin western blotting.

All samples for SDS electrophoresis in this study were prepared in Laemmli sample buffer supplemented with 50mM DTT and 6M Urea.

#### 13. Fluorescent degradation assay in cell extract

Hela S3 extract was supplemented with 1x energy-mix and 1x protease inhibitor cocktail (Sigma). 200nM substrates (unmodified or preubiquitylated) was mixed in and the extract was incubated at 30C on a plate reader (Biotek Synergy H1). Signal was detected every minute by a build-in monochromator (ex: 470nM/em: 510nM).

For reactions involving proteasome or E1 inhibitor, 50μM bortezomib or 40μM MLN7243 was added, and extract was incubated at room temperature for 15 minutes before adding substrates.

#### 14. Reconstituted ubiquitylation degradation system

Reactions involving preubiquitylated substrates: Substrates were first ubiquitylated by purified APC + UbcH10 in the presence of either Ub or UbL73F and then added to the reaction at a final concentration of 150nM. The reaction contains purified HUWE1, 5nM purified 26S proteasome, 50nM E1, 1μM UbcH5a, an energy regenerating system, 50μM ubiquitin (or 50μM methylated ubiquitin) in buffer R (25mM Tris-HCL [pH7.5], 50mM NaCl, 5mM MgCl2, 5% glycerol, 0.05% NP-40, 0.5mM DTT). The reaction was incubated at 30C and the fluorescence signal was collected every minute on a plate reader (Biotek Synergy H1).

Reactions involving unmodified substrate: 150nM unmodified substrate (cycB-cpGFP) was added to a reaction containing 30nM purified APC, 1μM purified Cdh1, purified HUWE1, 5nM purified 26S proteasome, 50nM E1, 500nM UbcH5a, 500nM UbcH10, an energy regenerating system, 50μM ubiquitin, and various concentrations of Usp2 in buffer R. The reaction was incubated at 30C and the fluorescence signal was collected every minute on a plate reader (Biotek Synergy H1).

#### 15. P97-mediated unfolding assay

Purification of recombinant human p97, NPL4 and UFD1, and preparing the ubiquitylated positive control substrate (Ub^N^-Dendra2) was detailed in a previous study (*48*).

To setup the unfolding assay, purified p97, UFD1 and NPL4 was first combined to form the complex by incubating in buffer P (50mM HEPES[pH 7.5], 130mM KCl, 10mM MgCl2, 0.5mM TCEP, 0.5 mg/mL protease-free BSA, 1x protease inhibitor cocktail) at 35C for 10 minutes, in the absence of ATP.

HUWE1 substrates were briefly purified using Ni-NTA dynabeads after the HUWE1 reaction. Substrates were added to the reaction mix at a final concentration of 100nM together with 5mM ATP- Mg2+. The final concentration of p97 (monomer), NPL4 and UFD1 was also 100nM. For reactions involving Ub(3)-VASP-cpGFP, 1μM recombinant GroEL(D87K) was added to the reaction before incubation.

Reactions were read on a platereader at 550nm/585nm at 35C, sampled once every 30 seconds.

#### 16. Single-molecule imaging and data analysis

The procedure for single-molecule binding assay with immobilized proteasome has been described in our previous study(*33*). Briefly, purified securin was ubiquitylated by APC plus UbcH10 with dy550-labeled Ub. Purified 26S proteasomes and biotinylated MCP21 antibody were mixed to a final concentration of 20nM and 12.5nM respectively. The mixture was incubated at room temperature for 15 minutes then kept on ice until the experiment. 50μM proteasome inhibitor epoxomicin was added during the incubation. For the imaging experiments, the temperature in the microscope room was set to 29°C. The proteasome-antibody mix was loaded onto PEG-passivated slides coated with streptavidin and incubated for 3 minutes.

For measuring dwell time, unbound proteasomes were washed off and replaced with imaging buffer (UBAB, 3mM ATP, 40mM imidazole, 5mg/ml BSA; * Imidazole in the imaging buffer serves to reduce the nonspecific binding of fluorescently labeled proteins) containing diluted 5nM securin- polyUbdy550. Image acquisition was started immediately with <15 s delay. Time series were acquired at 200 ms per frame for 3 minutes using a custom fluorescence TIRF microscope equipped with an sCMOS (pco) camera which was controlled by Micromanager.

#### 17. Formation and clearance of aggresomes and stress granules AgDD

WT or HUWE1Δ cells stably expressing AgDD(GFP) were seeded on a poly-lysine-passivated glass-bottom 12- or 24- well culture plate in DMEM medium (without phenol red) plus 10% FBS supplemented with 1uM shield-1, at about 30% confluency. To induce agDD aggregation, purified recombinant FKBP was added to the culture to a final concentration of 100μM, right before imaging started. 1μg/ml colchicine or 20μM NexturastatA was added to the culture to study their effect on agDD aggresome formation.

For studying the spontaneous clearance, cells were imaged continuously after induction. 10μM MLN7243, 10μM NMS-873 or 100μM chloroquine was added to the culture, 30 minutes before adding FKBP, to study the dependence of the clearance on different pathways. For studying induced clearance, 10μg/ml cycloheximide(CHX) was added to the culture 30 minutes after adding FKBP.

##### Htt

WT or HUWE1Δ cells stably integrated with pTetOn-HttQ94-CFP were seeded on a poly-lysine- passivated glass-bottom 12- or 24- well culture plate in DMEM medium (without phenol red) plus 10% FBS, at 10%∼30% confluency. To induce Htt aggregation, 1μg/ml doxycycline was added to the culture. To study the clearance kinetics, the culture was washed for 4x with the growth medium without doxycycline (with a 30-minute incubation after each wash).

##### Synphilin-1

WT or HUWE1Δ cells stably integrated with pTetOn-synphilin_1-GFP were seeded on a poly-lysine- passivated glass-bottom 12- or 24- well culture plate in DMEM medium (without phenol red) plus 10% FBS, at ∼15% confluency. 1μg/ml doxycycline was added to induce synphilin expression 48 hours before the induction. To induce aggregation, 5μg/ml MG132 was added to the culture for 8 hours, as in a previous study (*64*). To study the clearance kinetics, cells were washed for 3x with a growth medium without MG132.

##### Native aggresome by proteasome inhibition

WT or HUWE1Δ cells were seeded on a poly-lysine-passivated glass-bottom 12- or 24- well culture plate in DMEM medium (without phenol red) plus 10% FBS, at ∼40% confluency. 5μg/ml MG132 was added to the culture to induce aggresome formation. Cells were harvested 10∼12 hours after adding MG132.

##### Stress granule

WT or HUWE1Δ cells stably expressing GFP-G3BP1 were seeded on a poly-lysine-passivated glass-bottom 12- or 24- well culture plate in DMEM medium (without phenol red) plus 10% FBS, at ∼50% confluency. Cells were treated with either 0.5mM sodium arsenite for 45 minutes or incubated at 42C for 60 minutes to induce stress granule. To induce clearance, cells were washed with a growth medium without arsenite for 2x (for arsenite), or were incubated at 37C (for heat shock).

#### 18. Immunocytochemistry and colocalization analysis

After the removal of the media and brief washes, the cells were fixed with 4% formaldehyde for 30 minutes at room temperature (RT). Permeabilization and blocking were performed for 1 hour at RT with HBBS buffer containing 0.3% Triton X-100 and 2% BSA. Anti-Ub conjugates antibody (FK2) or anti-p97/VCP (Proteintech #10736-1-AP) antibodies were added and incubated for 1 hour at RT. The cells were washed and incubated with Alexa 647 secondary antibodies for 1 hour in dark at RT. After removal of secondary antibodies and brief washes, the cells were stained with PROTEOSTAT® Aggresome detection kit (Enzo ENZ-51035-K100) for both aggresome and nuclear and further washed twice. One drop of the mounting medium was added on top of the cells and 18mm round coverslips were used to cover the cells and sealed. All images were collected using Yokagawa CSU-X1 spinning disk confocal on an inverted Nikon Ti fluorescence microscope.

For colocalization analysis of KO2-HUWE1 (or variants) with aggregates, cells were first transected with pTetOn-KO2-HUWE1 plasmids, and the expression of HUWE1 was induced for 24 hours by adding 1µg/ml doxycycline. Cells were then fixed with paraformaldehyde and imaged using confocal microscopy. The degree of colocalization of aggregates with HUWE1, Ub(FK2) or p97 was quantified using Fiji and Coloc 2.

For long-term storage of the sample, the cells were fixed with 4% formaldehyde for 30 minutes at RT. One drop of the mounting medium was added on top of the cells and 18mm round coverslips were used to cover the cells and sealed. All images were collected using Yokagawa CSU-X1 spinning disk confocal on an inverted Nikon Ti fluorescence microscope.

#### 19. Immunohistochemistry on mouse brain samples and data analysis

Paraformaldehyde-fixed and gelatin-embedded mouse brain sections from 9mo WT and Thy- Tau22 mice were prepared at 35µm thickness and immunostained with AT8 antibody (Thermo MN1020, 1:2000) recognizing pathological Tau, HUWE1 antibody (Bethyl A300-486A, 1:1000), Tubb3-647 (Biolegend 657406, 1:1000), and DAPI. SIM images were taken on Elyra 7 (Zeiss) with Z stacks and processed using SIM2 algorithm. Tau aggregates were characterized as: 1. Early-stage, where AT8 positive staining was obvious yet diffuse, accompanied by normal neuronal structures as determined by Tubb3 staining; 2. Mid-stage, where AT8 staining showed a clear fibril pattern with relatively organized neuronal structures as defined by Tubb3 staining; and 3. late-stage, where AT8 staining showed an exclusive aggregation pattern with severe disruption in the cytoskeletal organization based on Tubb3 staining. Tau-HUWE1 association was quantified by colocalizing AT8 channel and HUWE1 channel using Mander’s coefficient of Tau: M(tau) and of HUWE1: M(Hw1), and all data were plotted using Prism.

#### 20. Quantitative live cell imaging Cell cycle analysis

A log-phase culture of WT or HUWE1Δ cells stably expressing mAG-Geminin and mKO2-Cdt1 was grown and was split into an 8-well TEK culture dish coated with fibronectin in DMEM medium without phenol red. After 8 hours, the chamber was mounted on a Nikon Ti2 epi-fluorescence microscope equipped with an environmental chamber for controlling temperature, humidity and CO2 level. Images were acquired once every 7 minutes using a 20X objective. Live-cell microscopy was performed in the Nikon Imaging Center at Harvard Medical School.

For the experiments testing securin degradation, cells were transfected with a plasmid mix containing one plasmid carrying RFP-HUWE1 (or variant) under a tetracycline-inducible promoter and the other with securin-YFP under a CMV promoter at a ratio of 4:1, using TransIT-293 reagent 48 hours before imaging. 24 hours after transfection, 1μg/ml doxycycline was added to induce HUWE1 expression.

##### Reporter degradation

For the experiments testing the degradation of YFP reporters, WT or HUWE1Δ cells were transfected with a plasmid expressing Ub-R-YFP, diluted by 10X with an empty vector, 24 hours before imagining. 100μg/ml cycloheximide was added to the culture right before imaging to inhibit protein translation.

##### Clearance of aggregates

For studying the clearance of agDD aggresome, synphilin aggresome and stress granules, cells were mounted on a Nikon Ti2 epi-fluorescence microscope equipped with an environmental chamber for controlling temperature, humidity and CO2 level. Images were acquired once every 7∼30 minutes using a 20X or 40X objective.

For long-term imaging of Htt aggregates, cells were imaged in an Incucyte fluorescence imagers (GFP and widefield channels) inside an incubator, once every 30 minutes for 48 hours.

#### 21. Quantification of protein aggregates

Quantification of stress granules was carried out as described in a previous study (*6*). Briefly, automatic cell segmentation was performed using CellProfiler based on the fluorescence channels. Segmentation results were then validated manually to exclude or correct obvious segmentation errors. Subcellular features (stress granule) identification was performed using Fiji and MiToBo(*83*). Cells with at least two G3BP1 aggregates were counted as containing stress granules.

For quantifying aggresomes (agDD, Htt, synphilin), the intensity of each aggresome in a time series was tracked automatically using p53Cinema, an automatic feature tracking and image quantification software(*59*), with an averaging disk diameter of 3μm. We define aggresome clearance as when its intensity decreases to less than 50% of the initial intensity or disappearance.

#### 22. Cell viability assay

Cell viability assay was performed using CytoTox-Glo™ Cytotoxicity Assay (Promega) as suggested by the manual.

## References

1. J. Labbadia, R. I. Morimoto, The biology of proteostasis in aging and disease. Annu. Rev. Biochem. 84, 435–464 (2015).

2. I. Dikic, B. A. Schulman, An expanded lexicon for the ubiquitin code. Nat. Rev. Mol. Cell Biol., 1– 15 (2022).

3. D. Komander, M. Rape, The ubiquitin code. Annu. Rev. Biochem. 81, 203–229 (2012).

4. J. Lowe, A. Blanchard, K. Morrell, G. Lennox, L. Reynolds, M. Billett, M. Landon, R. J. Mayer, Ubiquitin is a common factor in intermediate filament inclusion bodies of diverse type in man, including those of Parkinson’s disease, Pick’s disease, and Alzheimer’s disease, as well as Rosenthal fibres in cerebellar astrocytomas, cytoplasmic bodies in muscle, and mallory bodies in alcoholic liver disease. J. Pathol. 155, 9–15 (1988).

5. A. Alves-Rodrigues, L. Gregori, M. E. Figueiredo-Pereira, Ubiquitin, cellular inclusions and their role in neurodegeneration. Trends Neurosci. 21, 516–520 (1998).

6. B. A. Maxwell, Y. Gwon, A. Mishra, J. Peng, H. Nakamura, K. Zhang, H. J. Kim, J. P. Taylor, Ubiquitination is essential for recovery of cellular activities after heat shock. Science. 372, eabc3593 (2021).

7. N. F. Darwich, J. M. Phan, B. Kim, E. Suh, J. D. Papatriantafyllou, L. Changolkar, A. T. Nguyen, C. M. O’Rourke, Z. He, S. Porta, G. S. Gibbons, K. C. Luk, S. G. Papageorgiou, M. Grossman, L. Massimo, D. J. Irwin, C. T. McMillan, I. M. Nasrallah, C. Toro, G. K. Aguirre, V. M. Van Deerlin, E. B. Lee, Autosomal dominant VCP hypomorph mutation impairs disaggregation of PHF-tau. Science. 370 (2020), doi:10.1126/science.aay8826.

8. J. R. Buchan, R.-M. Kolaitis, J. P. Taylor, R. Parker, Eukaryotic stress granules are cleared by autophagy and Cdc48/VCP function. Cell. 153, 1461–1474 (2013).

9. S. Mukkavalli, J. A. Klickstein, B. Ortiz, P. Juo, M. Raman, The p97-UBXN1 complex regulates aggresome formation. J. Cell Sci. 134 (2021), doi:10.1242/jcs.254201.

10. D. K. Ghosh, A. Roy, A. Ranjan, The ATPase VCP/p97 functions as a disaggregase against toxic Huntingtin-exon1 aggregates. FEBS Lett. 592, 2680–2692 (2018).

11. J. van den Boom, H. Meyer, VCP/p97-Mediated Unfolding as a Principle in Protein Homeostasis and Signaling. Mol. Cell. 69, 182–194 (2018).

12. N. Tolay, A. Buchberger, Role of the Ubiquitin System in Stress Granule Metabolism. Int. J. Mol. Sci. 23 (2022), doi:10.3390/ijms23073624.

13. I. Saha, P. Yuste-Checa, M. Da Silva Padilha, Q. Guo, R. Körner, H. Holthusen, V. A. Trinkaus, I. Dudanova, R. Fernández-Busnadiego, W. Baumeister, D. W. Sanders, S. Gautam, M. I. Diamond, F.U. Hartl, M. S. Hipp, The AAA+ chaperone VCP disaggregates Tau fibrils and generates aggregate seeds in a cellular system. Nat. Commun. 14, 560 (2023).

14. J.-S. Ju, S. E. Miller, P. I. Hanson, C. C. Weihl, Impaired protein aggregate handling and clearance underlie the pathogenesis of p97/VCP-associated disease. J. Biol. Chem. 283, 30289–30299 (2008).

15. G. Papiani, A. Ruggiano, M. Fossati, A. Raimondi, G. Bertoni, M. Francolini, R. Benfante, F. Navone, N. Borgese, Restructured endoplasmic reticulum generated by mutant amyotrophic lateral sclerosis-linked VAPB is cleared by the proteasome. J. Cell Sci. 125, 3601–3611 (2012).

16. M. Słabicki, H. Yoon, J. Koeppel, L. Nitsch, S. S. Roy Burman, C. Di Genua, K. A. Donovan, A. S. Sperling, M. Hunkeler, J. M. Tsai, R. Sharma, A. Guirguis, C. Zou, P. Chudasama, J. A. Gasser, P. G. Miller, C. Scholl, S. Fröhling, R. P. Nowak, E. S. Fischer, B. L. Ebert, Small-molecule-induced polymerization triggers degradation of BCL6. Nature. 588, 164–168 (2020).

17. E. Mee Hayes, L. Sirvio, Y. Ye, A Potential Mechanism for Targeting Aggregates With Proteasomes and Disaggregases in Liquid Droplets. Front. Aging Neurosci. 14, 854380 (2022).

18. F. Limanaqi, F. Biagioni, S. Gambardella, P. Familiari, A. Frati, F. Fornai, Promiscuous Roles of Autophagy and Proteasome in Neurodegenerative Proteinopathies. Int. J. Mol. Sci. 21 (2020), doi:10.3390/ijms21083028.

19. R. Hjerpe, J. S. Bett, M. J. Keuss, A. Solovyova, T. G. McWilliams, C. Johnson, I. Sahu, J. Varghese, N. Wood, M. Wightman, G. Osborne, G. P. Bates, M. H. Glickman, M. Trost, A. Knebel, F. Marchesi, T. Kurz, UBQLN2 Mediates Autophagy-Independent Protein Aggregate Clearance by the Proteasome. Cell. 166, 935–949 (2016).

20. A. Hershko, A. Ciechanover, The ubiquitin system. Annu. Rev. Biochem. 67, 425–479 (1998).

21. B. Crosas, J. Hanna, D. S. Kirkpatrick, D. P. Zhang, Y. Tone, N. A. Hathaway, C. Buecker, D. S. Leggett, M. Schmidt, R. W. King, S. P. Gygi, D. Finley, Ubiquitin chains are remodeled at the proteasome by opposing ubiquitin ligase and deubiquitinating activities. Cell. 127, 1401–1413 (2006).

22. S. Aviram, D. Kornitzer, The ubiquitin ligase Hul5 promotes proteasomal processivity. Mol. Cell. Biol. 30, 985–994 (2010).

23. T. Van Nguyen, J. Li, C.-C. J. Lu, J. L. Mamrosh, G. Lu, B. E. Cathers, R. J. Deshaies, p97/VCP promotes degradation of CRBN substrate glutamine synthetase and neosubstrates. Proc. Natl. Acad. Sci. U. S. A. 114, 3565–3571 (2017).

24. K. A. Donovan, F. M. Ferguson, J. W. Bushman, N. A. Eleuteri, D. Bhunia, S. Ryu, L. Tan, K. Shi, H. Yue, X. Liu, D. Dobrovolsky, B. Jiang, J. Wang, M. Hao, I. You, M. Teng, Y. Liang, J. Hatcher, Z. Li, T. D. Manz, B. Groendyke, W. Hu, Y. Nam, S. Sengupta, H. Cho, I. Shin, M. P. Agius, I. M. Ghobrial, M. W. Ma, J. Che, S. J. Buhrlage, T. Sim, N. S. Gray, E. S. Fischer, Mapping the Degradable Kinome Provides a Resource for Expedited Degrader Development. Cell. 183, 1714–1731.e10 (2020).

25. X. Gong, D. Du, Y. Deng, Y. Zhou, L. Sun, S. Yuan, The structure and regulation of the E3 ubiquitin ligase HUWE1 and its biological functions in cancer. Invest. New Drugs. 38, 515–524 (2020).

26. S.-H. Kao, H.-T. Wu, K.-J. Wu, Ubiquitination by HUWE1 in tumorigenesis and beyond. J. Biomed. Sci. 25, 67 (2018).

27. A. C. Giles, B. Grill, Roles of the HUWE1 ubiquitin ligase in nervous system development, function and disease. Neural Dev. 15, 6 (2020).

28. M. Békés, K. Okamoto, S. B. Crist, M. J. Jones, J. R. Chapman, B. B. Brasher, F. D. Melandri, B. M. Ueberheide, E. L. Denchi, T. T. Huang, DUB-resistant ubiquitin to survey ubiquitination switches in mammalian cells. Cell Rep. 5, 826–838 (2013).

29. K. Martinez-Fonts, C. Davis, T. Tomita, S. Elsasser, A. R. Nager, Y. Shi, D. Finley, A. Matouschek, The proteasome 19S cap and its ubiquitin receptors provide a versatile recognition platform for substrates. Nat. Commun. 11, 477 (2020).

30. Y. Lu, W. Wang, M. W. Kirschner, Specificity of the anaphase-promoting complex: a single- molecule study. Science. 348, 1248737 (2015).

31. N. G. Brown, R. VanderLinden, E. R. Watson, F. Weissmann, A. Ordureau, K.-P. Wu, W. Zhang, S. Yu, P. Y. Mercredi, J. S. Harrison, I. F. Davidson, R. Qiao, Y. Lu, P. Dube, M. R. Brunner, C. R. R. Grace, D. J. Miller, D. Haselbach, M. A. Jarvis, M. Yamaguchi, D. Yanishevski, G. Petzold, S. S. Sidhu, B. Kuhlman, M. W. Kirschner, J. W. Harper, J.-M. Peters, H. Stark, B. A. Schulman, Dual RING E3 Architectures Regulate Multiubiquitination and Ubiquitin Chain Elongation by APC/C. Cell. 165, 1440–1453 (2016).

32. S. A. H. de Poot, G. Tian, D. Finley, Meddling with Fate: The Proteasomal Deubiquitinating Enzymes. J. Mol. Biol. 429, 3525–3545 (2017).

33. Y. Lu, B.-H. Lee, R. W. King, D. Finley, M. W. Kirschner, Substrate degradation by the proteasome: a single-molecule kinetic analysis. Science. 348, 1250834 (2015).

34. J. Hon, Y. Lu, “Chapter Ten - Single-molecule methods for measuring ubiquitination and protein stability” in Methods in Enzymology, M. Hochstrasser, Ed. (Academic Press, 2019), vol. 619, pp. 225–247.

35. M. Hunkeler, C. Y. Jin, M. W. Ma, J. K. Monda, D. Overwijn, E. J. Bennett, E. S. Fischer, Solenoid architecture of HUWE1 contributes to ligase activity and substrate recognition. Mol. Cell (2021), doi:10.1016/j.molcel.2021.06.032.

36. B. Schwanhäusser, D. Busse, N. Li, G. Dittmar, J. Schuchhardt, J. Wolf, W. Chen, M. Selbach, Global quantification of mammalian gene expression control. Nature. 473, 337–342 (2011).

37. T. Hoppe, Multiubiquitylation by E4 enzymes: “one size” doesn’t fit all. Trends Biochem. Sci. 30, 183–187 (2005).

38. A. Bremm, S. M. V. Freund, D. Komander, Lys11-linked ubiquitin chains adopt compact conformations and are preferentially hydrolyzed by the deubiquitinase Cezanne. Nat. Struct. Mol. Biol. 17, 939–947 (2010).

39. M. A. Michel, K. N. Swatek, M. K. Hospenthal, D. Komander, Ubiquitin Linkage-Specific Affimers Reveal Insights into K6-Linked Ubiquitin Signaling. Mol. Cell. 68, 233–246.e5 (2017).

40. S. Brühmann, D. S. Ushakov, M. Winterhoff, R. B. Dickinson, U. Curth, J. Faix, Distinct VASP tetramers synergize in the processive elongation of individual actin filaments from clustered arrays. Proc. Natl. Acad. Sci. U. S. A. 114, E5815–E5824 (2017).

41. Y. Xu, D. E. Anderson, Y. Ye, The HECT domain ubiquitin ligase HUWE1 targets unassembled soluble proteins for degradation. Cell Discov. 2, 16040 (2016).

42. K. L. Rock, C. Gramm, L. Rothstein, K. Clark, R. Stein, L. Dick, D. Hwang, A. L. Goldberg, Inhibitors of the proteasome block the degradation of most cell proteins and the generation of peptides presented on MHC class I molecules. Cell. 78, 761–771 (1994).

43. J. Li, Z. Cai, L. P. Vaites, N. Shen, D. C. Mitchell, E. L. Huttlin, J. A. Paulo, B. L. Harry, S. P. Gygi, Proteome-wide mapping of short-lived proteins in human cells. Mol. Cell. 81, 4722–4735.e5 (2021).

44. S. X. Ge, D. Jung, R. Yao, ShinyGO: a graphical gene-set enrichment tool for animals and plants. Bioinformatics. 36, 2628–2629 (2020).

45. J. M. Replogle, R. A. Saunders, A. N. Pogson, J. A. Hussmann, A. Lenail, A. Guna, L. Mascibroda, E. J. Wagner, K. Adelman, G. Lithwick-Yanai, N. Iremadze, F. Oberstrass, D. Lipson, J. L. Bonnar, M. Jost, T. M. Norman, J. S. Weissman, Mapping information-rich genotype-phenotype landscapes with genome-scale Perturb-seq. Cell (2022), doi:10.1016/j.cell.2022.05.013.

46. T. D. Deegan, P. P. Mukherjee, R. Fujisawa, C. Polo Rivera, K. Labib, CMG helicase disassembly is controlled by replication fork DNA, replisome components and a ubiquitin threshold. Elife. 9 (2020), doi:10.7554/eLife.60371.

47. H. Tsuchiya, F. Ohtake, N. Arai, A. Kaiho, S. Yasuda, K. Tanaka, Y. Saeki, In Vivo Ubiquitin Linkage-type Analysis Reveals that the Cdc48-Rad23/Dsk2 Axis Contributes to K48-Linked Chain Specificity of the Proteasome. Mol. Cell. 66, 488–502.e7 (2017).

48. Z. Ji, H. Li, D. Peterle, J. A. Paulo, S. B. Ficarro, T. E. Wales, J. A. Marto, S. P. Gygi, J. R. Engen, T. A. Rapoport, Translocation of polyubiquitinated protein substrates by the hexameric Cdc48 ATPase. Mol. Cell. 82, 570–584.e8 (2022).

49. D. S. Kirkpatrick, N. A. Hathaway, J. Hanna, S. Elsasser, J. Rush, D. Finley, R. W. King, S. P. Gygi, Quantitative analysis of in vitro ubiquitinated cyclin B1 reveals complex chain topology. Nat. Cell Biol. 8, 700–710 (2006).

50. Y. Lu, J. Wu, Y. Dong, S. Chen, S. Sun, Y.-B. Ma, Q. Ouyang, D. Finley, M. W. Kirschner, Y. Mao, Conformational Landscape of the p28-Bound Human Proteasome Regulatory Particle. Mol. Cell. 67, 322–333.e6 (2017).

51. E. U. Weber-Ban, B. G. Reid, A. D. Miranker, A. L. Horwich, Global unfolding of a substrate protein by the Hsp100 chaperone ClpA. Nature. 401, 90–93 (1999).

52. M. Békés, D. R. Langley, C. M. Crews, PROTAC targeted protein degraders: the past is prologue. Nat. Rev. Drug Discov. 21, 181–200 (2022).

53. N. V. Dimova, N. A. Hathaway, B.-H. Lee, D. S. Kirkpatrick, M. L. Berkowitz, S. P. Gygi, D. Finley, R. W. King, APC/C-mediated multiple monoubiquitylation provides an alternative degradation signal for cyclin B1. Nat. Cell Biol. 14, 168–176 (2012).

54. M. Rape, S. K. Reddy, M. W. Kirschner, The processivity of multiubiquitination by the APC determines the order of substrate degradation. Cell. 124, 89–103 (2006).

55. O. Cohen-Fix, J. M. Peters, M. W. Kirschner, D. Koshland, Anaphase initiation in Saccharomyces cerevisiae is controlled by the APC-dependent degradation of the anaphase inhibitor Pds1p. Genes Dev. 10, 3081–3093 (1996).

56. E. E. Arias, J. C. Walter, Replication-dependent destruction of Cdt1 limits DNA replication to a single round per cell cycle in Xenopus egg extracts. Genes Dev. 19, 114–126 (2005).

57. S.-B. Koh, P. Mascalchi, E. Rodriguez, Y. Lin, D. I. Jodrell, F. M. Richards, S. K. Lyons, A quantitative FastFUCCI assay defines cell cycle dynamics at a single-cell level. J. Cell Sci. 130, 512–520 (2017).

58. A. Sakaue-Sawano, H. Kurokawa, T. Morimura, A. Hanyu, H. Hama, H. Osawa, S. Kashiwagi, K. Fukami, T. Miyata, H. Miyoshi, T. Imamura, M. Ogawa, H. Masai, A. Miyawaki, Visualizing spatiotemporal dynamics of multicellular cell-cycle progression. Cell. 132, 487–498 (2008).

59. J. Reyes, J.-Y. Chen, J. Stewart-Ornstein, K. W. Karhohs, C. S. Mock, G. Lahav, Fluctuations in p53 Signaling Allow Escape from Cell-Cycle Arrest. Mol. Cell. 71, 581–591.e5 (2018).

60. Y. Miyazaki, K. Mizumoto, G. Dey, T. Kudo, J. Perrino, L.-C. Chen, T. Meyer, T. J. Wandless, A method to rapidly create protein aggregates in living cells. Nat. Commun. 7, 11689 (2016).

61. R. R. Kopito, Aggresomes, inclusion bodies and protein aggregation. Trends Cell Biol. 10, 524–530 (2000).

62. Y. Kawaguchi, J. J. Kovacs, A. McLaurin, J. M. Vance, A. Ito, T. P. Yao, The deacetylase HDAC6 regulates aggresome formation and cell viability in response to misfolded protein stress. Cell. 115, 727–738 (2003).

63. J. A. Johnston, C. L. Ward, R. R. Kopito, Aggresomes: a cellular response to misfolded proteins. J. Cell Biol. 143, 1883–1898 (1998).

64. N. Zaarur, A. B. Meriin, V. L. Gabai, M. Y. Sherman, Triggering Aggresome Formation: DISSECTING AGGRESOME-TARGETING AND AGGREGATION SIGNALS IN SYNPHILIN 1. J. Biol. Chem. 283, 27575–27584 (2008).

65. K. Schindowski, A. Bretteville, K. Leroy, S. Bégard, J.-P. Brion, M. Hamdane, L. Buée, Alzheimer’s disease-like tau neuropathology leads to memory deficits and loss of functional synapses in a novel mutated tau transgenic mouse without any motor deficits. Am. J. Pathol. 169, 599–616 (2006).

66. C. Mayor-Ruiz, M. G. Jaeger, S. Bauer, M. Brand, C. Sin, A. Hanzl, A. C. Mueller, J. Menche, G. E. Winter, Plasticity of the Cullin-RING Ligase Repertoire Shapes Sensitivity to Ligand-Induced Protein Degradation. Mol. Cell. 75, 849–858.e8 (2019).

67. T. Kobayashi, A. Manno, A. Kakizuka, Involvement of valosin-containing protein (VCP)/p97 in the formation and clearance of abnormal protein aggregates. Genes Cells. 12, 889–901 (2007).

68. R. Fujisawa, C. Polo Rivera, K. P. M. Labib, Multiple UBX proteins reduce the ubiquitin threshold of the mammalian p97-UFD1-NPL4 unfoldase. Elife. 11 (2022), doi:10.7554/eLife.76763.

69. R. G. Yau, K. Doerner, E. R. Castellanos, D. L. Haakonsen, A. Werner, N. Wang, X. W. Yang, N. Martinez-Martin, M. L. Matsumoto, V. M. Dixit, M. Rape, Assembly and Function of Heterotypic Ubiquitin Chains in Cell-Cycle and Protein Quality Control. Cell. 171, 918–933.e20 (2017).

70. R. P. Kreiser, A. K. Wright, N. R. Block, J. E. Hollows, L. T. Nguyen, K. LeForte, B. Mannini, M. Vendruscolo, R. Limbocker, Therapeutic Strategies to Reduce the Toxicity of Misfolded Protein Oligomers. Int. J. Mol. Sci. 21 (2020), doi:10.3390/ijms21228651.

71. J. Hanna, N. A. Hathaway, Y. Tone, B. Crosas, S. Elsasser, D. S. Kirkpatrick, D. S. Leggett, S. P. Gygi, R. W. King, D. Finley, Deubiquitinating enzyme Ubp6 functions noncatalytically to delay proteasomal degradation. Cell. 127, 99–111 (2006).

72. A. A. Kudriaeva, I. Livneh, M. S. Baranov, R. H. Ziganshin, A. E. Tupikin, S. O. Zaitseva, M. R. Kabilov, A. Ciechanover, A. A. Belogurov Jr, In-depth characterization of ubiquitin turnover in mammalian cells by fluorescence tracking. Cell Chem Biol (2021), doi:10.1016/j.chembiol.2021.02.009.

73. F. V. Hundley, N. Sanvisens Delgado, H. C. Marin, K. L. Carr, R. Tian, D. P. Toczyski, A comprehensive phenotypic CRISPR-Cas9 screen of the ubiquitin pathway uncovers roles of ubiquitin ligases in mitosis. Mol. Cell. 81, 1319–1336.e9 (2021).

74. N. D. Udeshi, D. C. Mani, S. Satpathy, S. Fereshetian, J. A. Gasser, T. Svinkina, M. E. Olive, B. L. Ebert, P. Mertins, S. A. Carr, Rapid and deep-scale ubiquitylation profiling for biology and translational research. Nat. Commun. 11, 359 (2020).

75. M. Koegl, T. Hoppe, S. Schlenker, H. D. Ulrich, T. U. Mayer, S. Jentsch, A novel ubiquitination factor, E4, is involved in multiubiquitin chain assembly. Cell. 96, 635–644 (1999).

76. Z. Li, S. Chen, J.-H. Jhong, Y. Pang, K.-Y. Huang, S. Li, T.-Y. Lee, UbiNet 2.0: a verified, classified, annotated and updated database of E3 ubiquitin ligase–substrate interactions. Database . 2021 (2021), doi:10.1093/database/baab010.

77. M. L. van de Weijer, L. Krshnan, S. Liberatori, E. N. Guerrero, J. Robson-Tull, L. Hahn, R. J. Lebbink, E. J. H. J. Wiertz, R. Fischer, D. Ebner, P. Carvalho, Quality Control of ER Membrane Proteins by the RNF185/Membralin Ubiquitin Ligase Complex. Mol. Cell. 79, 768–781.e7 (2020).

78. K. C. Dong, E. Helgason, C. Yu, L. Phu, D. P. Arnott, I. Bosanac, D. M. Compaan, O. W. Huang, A. V. Fedorova, D. S. Kirkpatrick, S. G. Hymowitz, E. C. Dueber, Preparation of distinct ubiquitin chain reagents of high purity and yield. Structure. 19, 1053–1063 (2011).

79. K. A. Donovan, J. An, R. P. Nowak, J. C. Yuan, E. C. Fink, B. C. Berry, B. L. Ebert, E. S. Fischer, Thalidomide promotes degradation of SALL4, a transcription factor implicated in Duane Radial Ray syndrome. Elife. 7 (2018), doi:10.7554/eLife.38430.

80. F. Meier, A.-D. Brunner, M. Frank, A. Ha, I. Bludau, E. Voytik, S. Kaspar-Schoenefeld, M. Lubeck, O. Raether, N. Bache, R. Aebersold, B. C. Collins, H. L. Röst, M. Mann, diaPASEF: parallel accumulation-serial fragmentation combined with data-independent acquisition. Nat. Methods. 17, 1229–1236 (2020).

81. V. Demichev, C. B. Messner, S. I. Vernardis, K. S. Lilley, M. Ralser, DIA-NN: neural networks and interference correction enable deep proteome coverage in high throughput. Nat. Methods. 17, 41–44 (2020).

82. M. E. Ritchie, B. Phipson, D. Wu, Y. Hu, C. W. Law, W. Shi, G. K. Smyth, limma powers differential expression analyses for RNA-sequencing and microarray studies. Nucleic Acids Res. 43, e47 (2015).

83. S. Posch, MiToBo - A Toolbox for Image Processing and Analysis. 4, e17 (2016).

