## Supplementary material for "HUWE1 Amplifies Ubiquitin Modifications to Broadly Stimulate Clearance of Proteins and Aggregates": Supp. figures

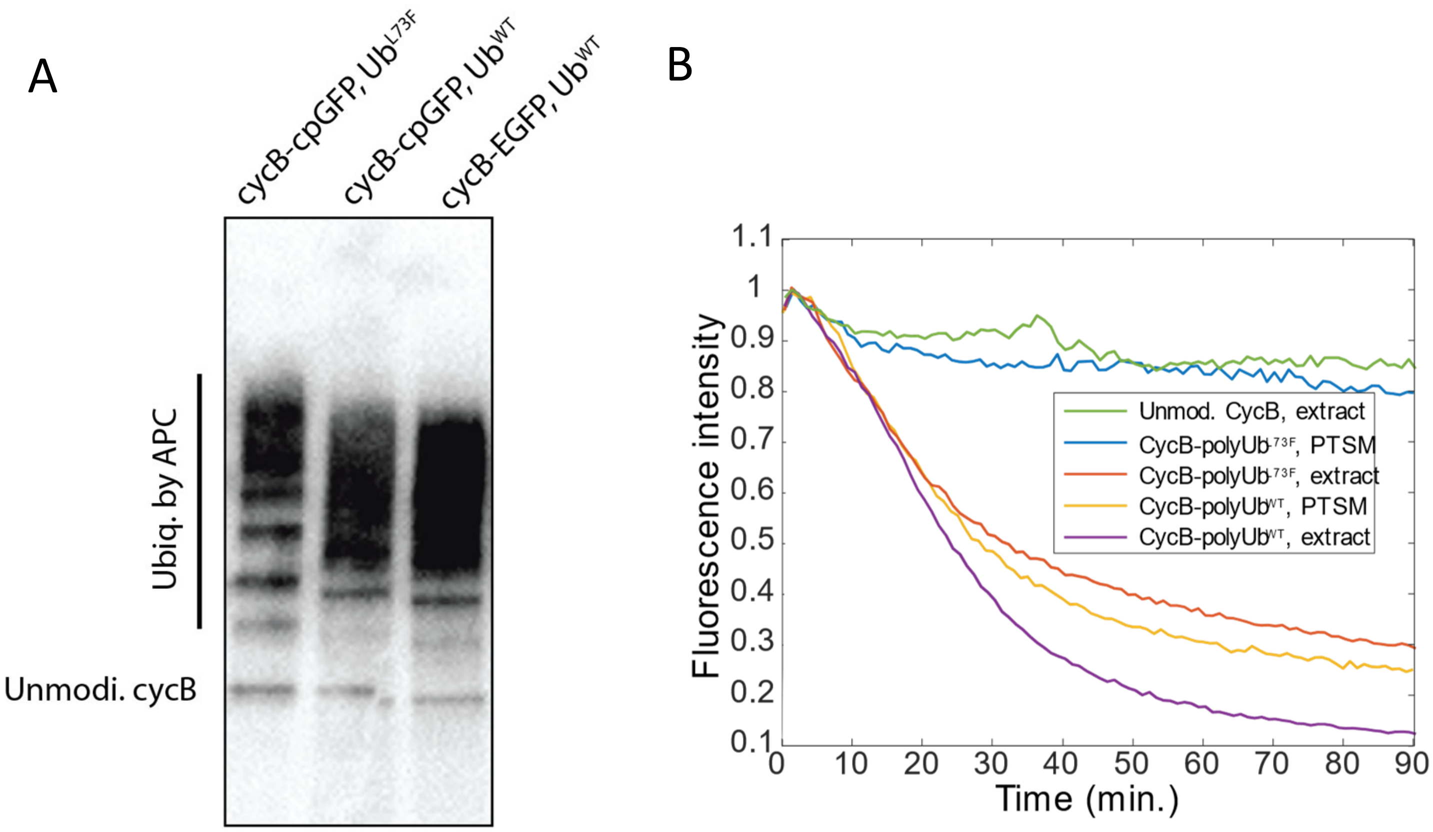


**Figure S1. Ub$^{L73F}$-conjugated cycB is degraded in cell extract but not by the purified 26S proteasome**.

**A.** Ubiquitylation of HA-cycB-cpGFP or HA-cycB-EGFP by APC-UbcH10 in the presence of Ub$^{WT}$ or Ub$^{L73F}$. Samples were analyzed by anti-HA western blotting. **B.** CycB-cpGFP, unmodified or poly-ubiquitylated by recombinant APC-UbcH10 with the indicated Ub variant, was incubated with either purified human 26S proteasome (PTSM) or in asynchronized HeLa Cell extract. GFP fluorescence was detected by a plate reader.

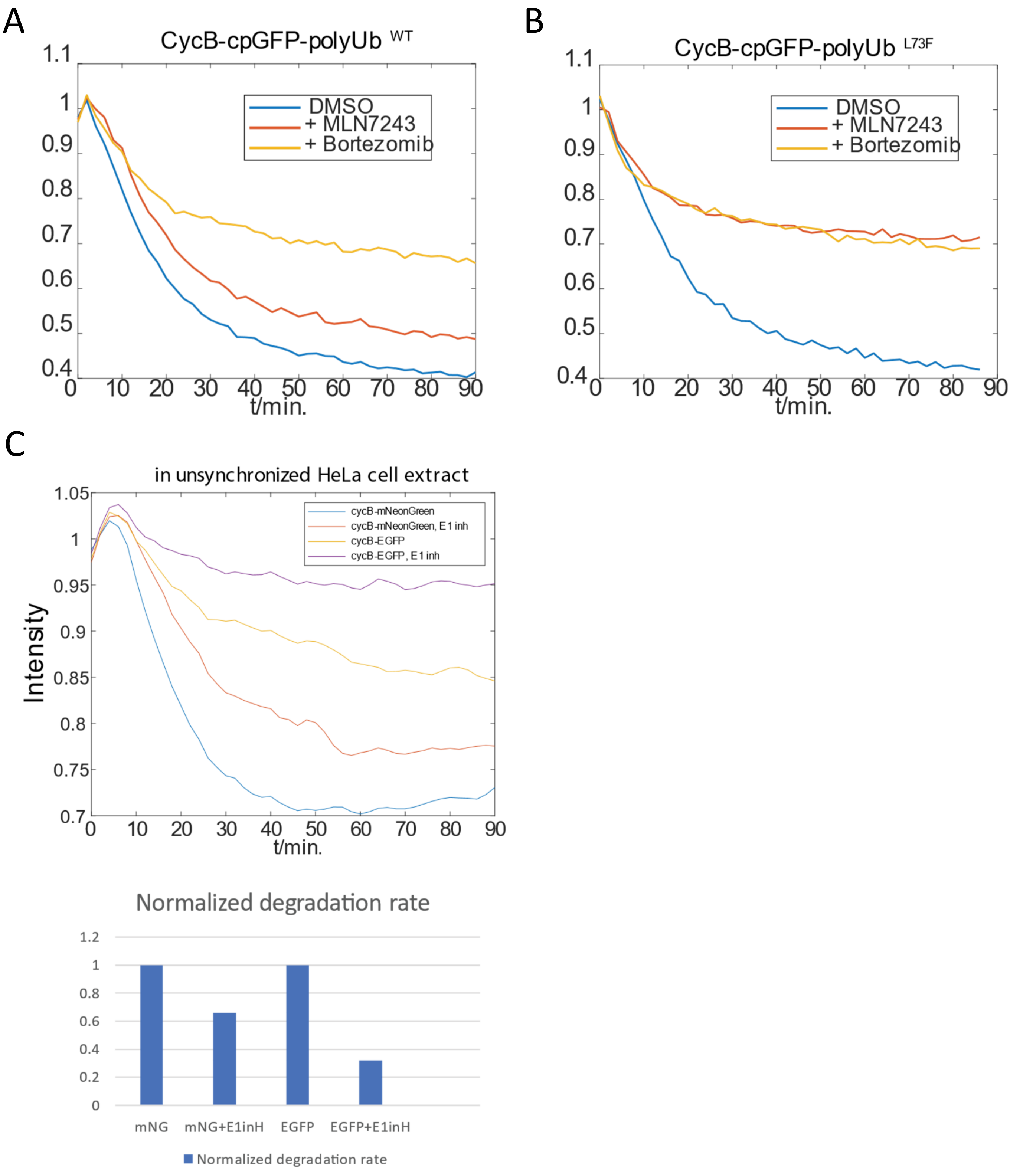


**Figure S2. Degradation of Ub$^{L73F}$-conjugates cycB substrates in cell extract depends on the proteasome and the Ub E1 activity in extract. A, B.** CycB-cpGFP conjugated with poly-Ub$^{WT}$ or poly-Ub$^{L73F}$ by recombinant APC-UbcH10 was incubated in asynchronous HeLa cell extract in the presence of a proteasome inhibitor (Bortezomib), an Ub E1 inhibitor (MLN7243) or DMSO. Fluorescent signal was detected on a plate reader. **C.** Polyubiquitylated cycB-EGFP or cycB-mNeonGreen(mNG) was incubated in HeLa cell extract, with or without the Ub E1 inhibitor (E1inh). The initial degradation rate of these substrates was calculated from the fluorescent signal that was detected on a plate reader.

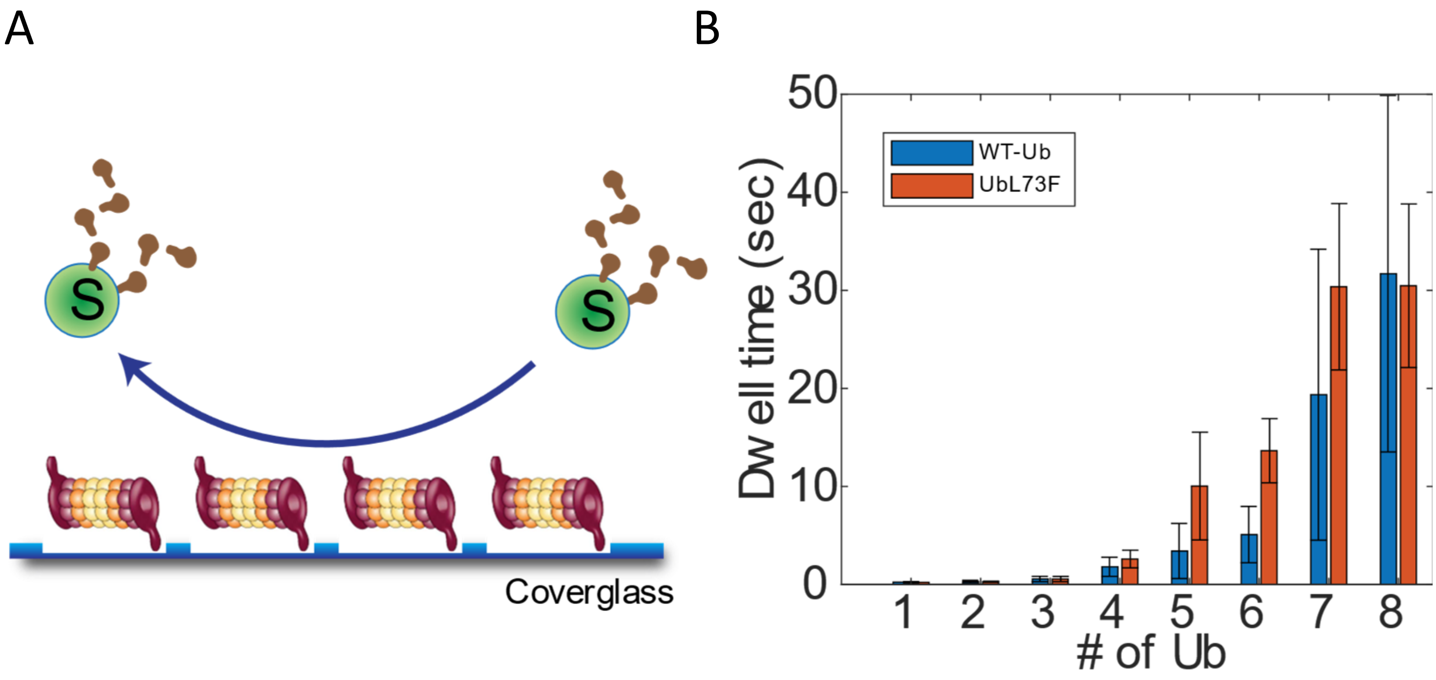


**Figure S3. The L73F mutation on Ub does not affect substrate interaction with proteasome. A.** Schematic of the single-molecule binding assay (see methods)(33). B. Dwell time distribution of securin, conjugated with poly-wtUb or poly-UbL73F that is also fluorescently labeled, on immobilized 26S proteasome. The dwell time is registered as a function of the copy number of Ubs on a securin molecule. Error bars represent the standard error of dwell time.

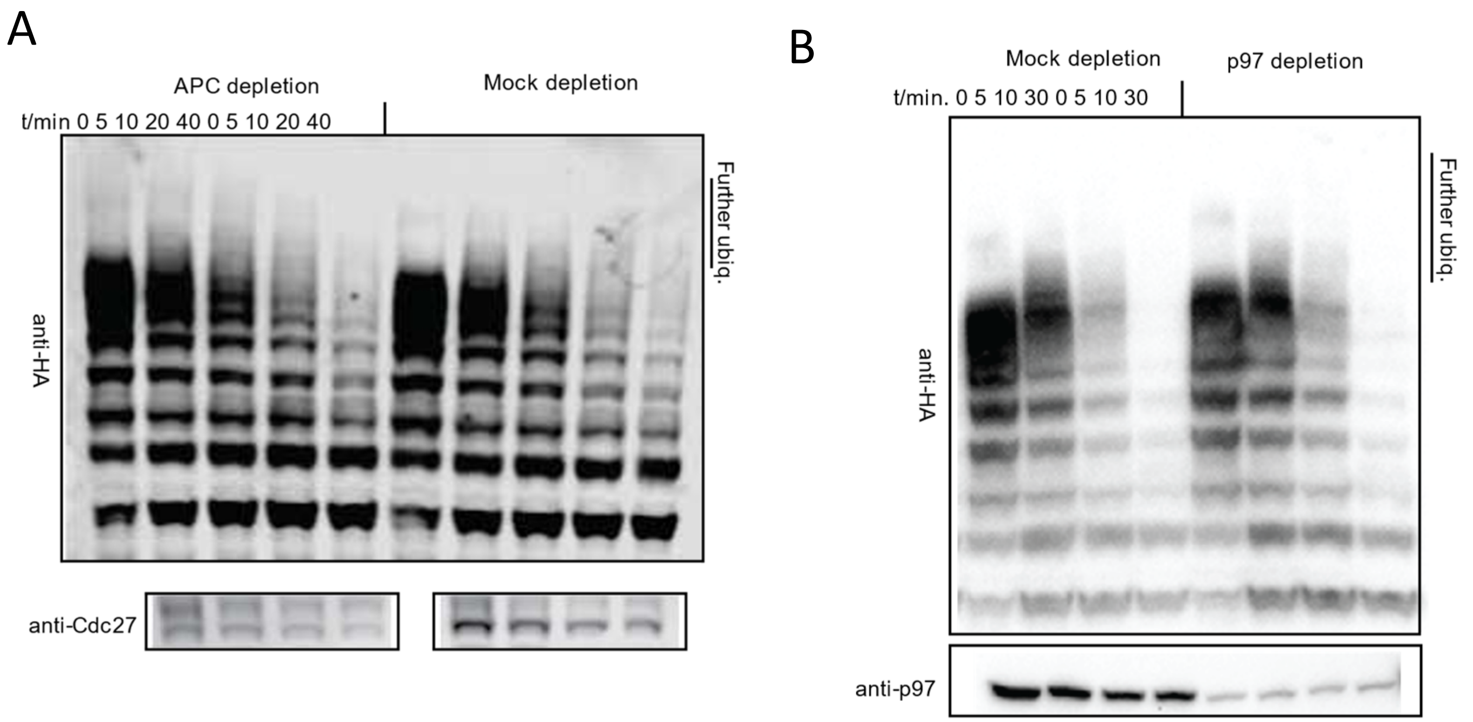


**Figure S4. Further ubiquitylation of cycB-cpGFP-polyUb$^{L73F}$ in extract is not due to any E3 or E4 activity associated with the APC or p97/VCP. A,B.** HeLa cell extract, whose APC or p97 had been immunodepleted using their respective antibodies(methods), was incubated with HA-cycB-cpGFP that has been ubiquitylated with Ub$^{L73F}$ by APC-UbcH10. The time-series samples were analyzed by anti-HA western blotting.

A

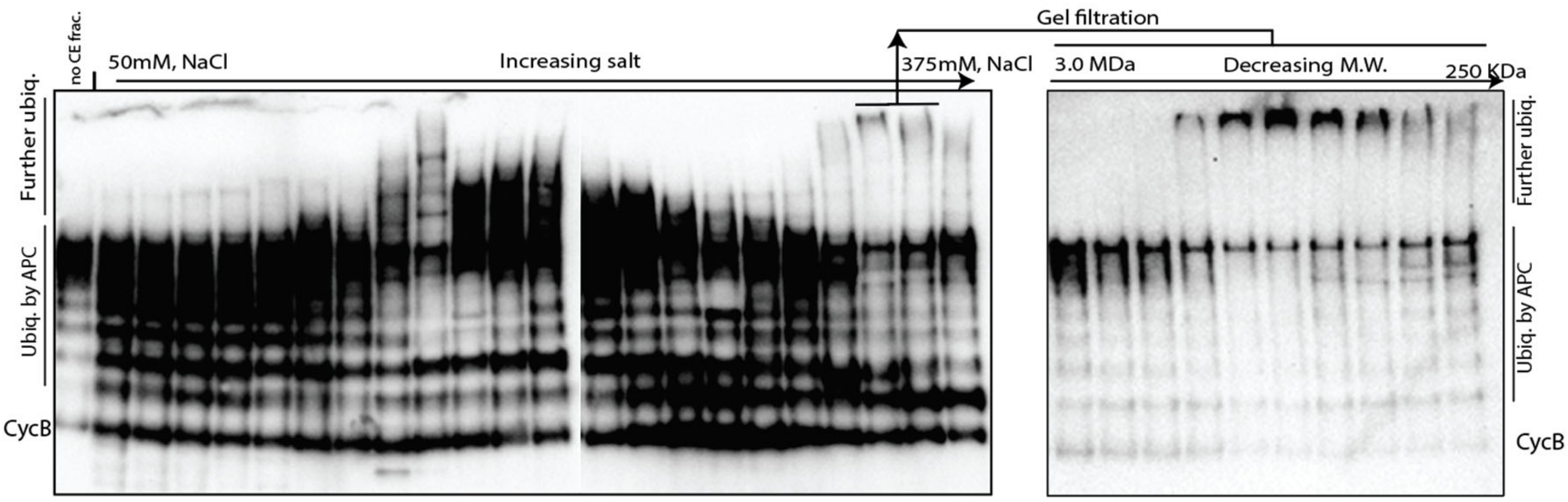


B

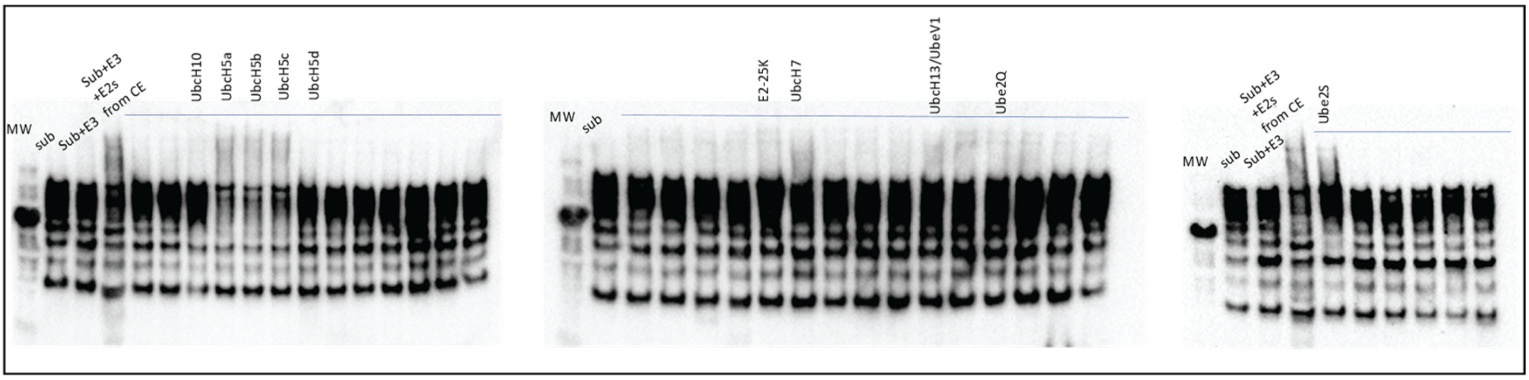


**Figure S5. Cell extract contains factors that can further ubiquitylate cycB-cpGFP-polyUb$^{L73F}$. A.** Hela extract was fractioned by anion exchange (monoQ) chromatography. Each fraction was tested for ubiquitylating HA-cycB-cpGFP-polyUb$^{L73F}$ in the presence of E2 UbcH5a. Fractions with highest activities (indicated) were grouped and fractionated again by size-exclusion (Superose 6) chromatography. The resulting fractions were tested likewise. **B.** E2 scanning. 34 E2s were tested for their activity in mediating the ubiquitylation of HA-cycB-cpGFP-polyUb$^{L73F}$ by the active FPLC fraction from A.

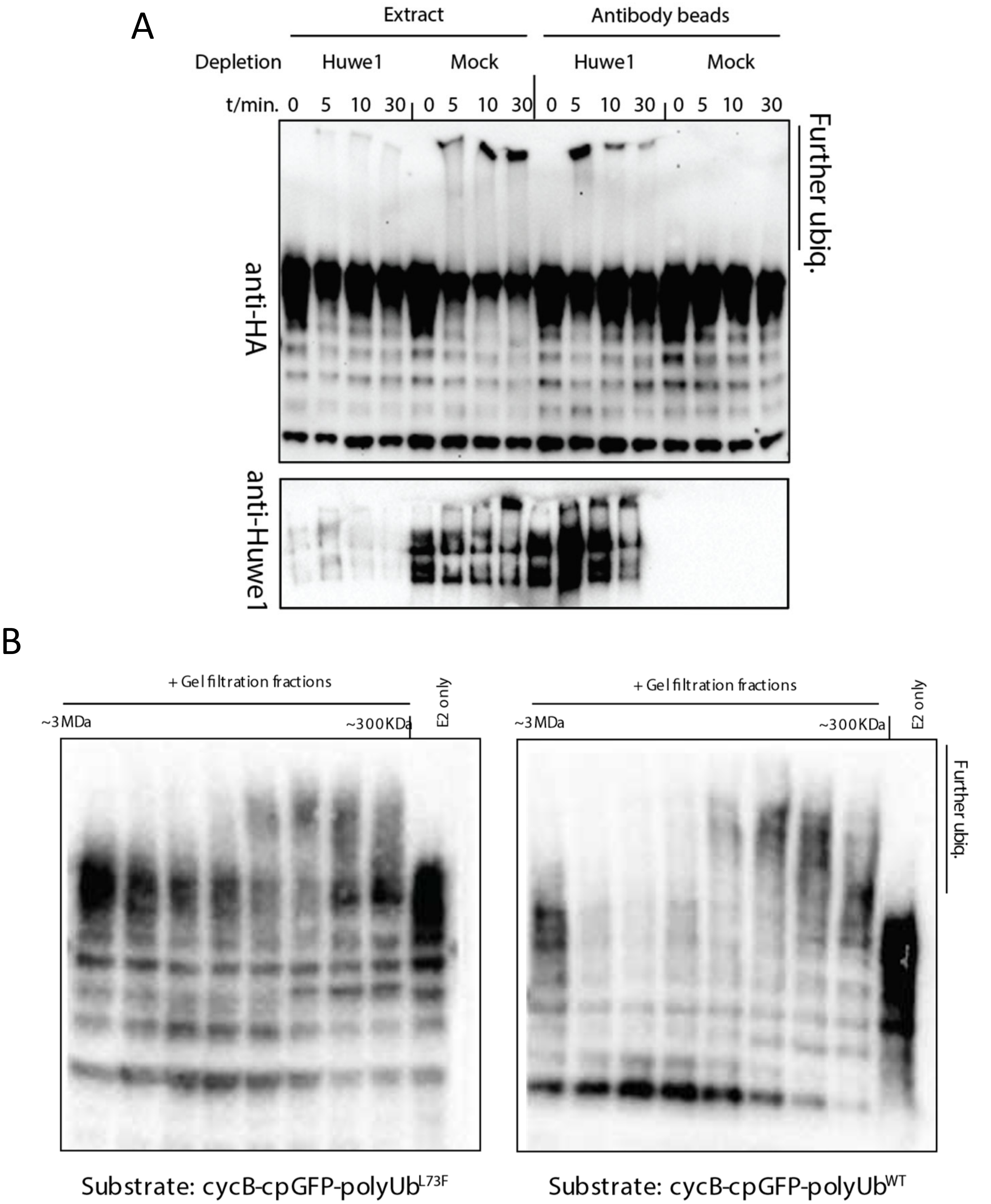


**Figure S6. HUWE1 is responsible for the further ubiquitylation of cycB-cpGFP-polyUb$^{L73F}$ in an active fraction and can react with wtUb-conjugated cycB with a similar efficiency. A.** HUWE1 was immunodepleted from the active fractions (grouped) in Fig. S5A. The depleted fraction or the anti-HUWE1 beads were tested for ubiquitylating HA-cycB-cpGFP-polyUb$^{L73F}$. **B.** Ubiquitylation of HA-cycB-cpGFP-polyUb$^{L73F}$ or HA-cycB-cpGFP-polyUb$^{WT}$ by the gel-filtration fractions described in Fig. S1A.

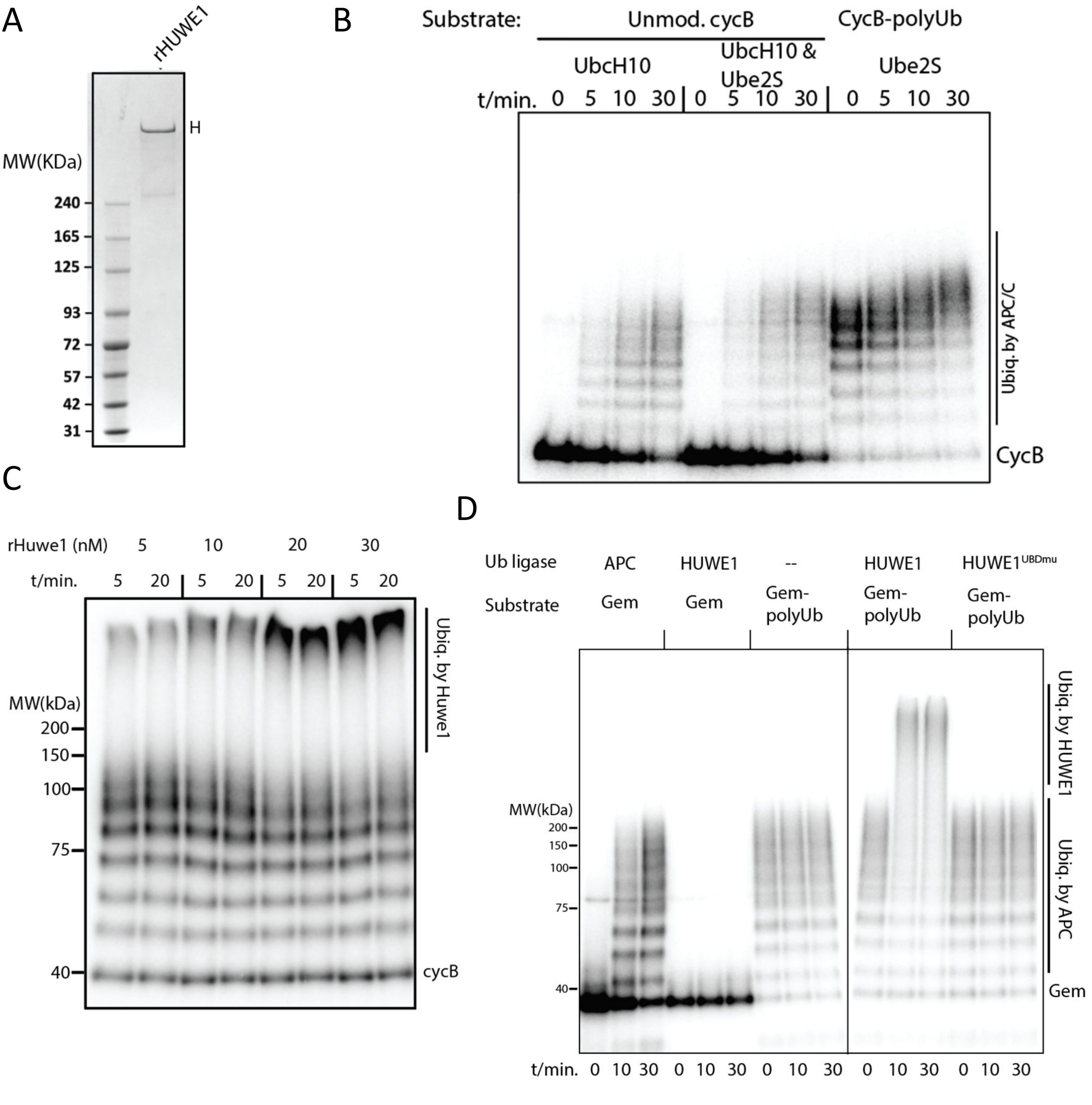


**Figure S7. HUWE1's UDL activity maintains high processivity at low nM concentrations. A.** Purification of recombinant HUWE1. Coomassie stain of SDS-PAGE. **B.** Titration of rHUWE1 concentration in ubiquitylating p$^{32}$-labeled cycB-cpGFP-polyUb. Samples were analyzed by autoradiography. **C.** P$^{32}$-labeled cycB-cpGFP was ubiquitylated by purified APC with E2 UbcH10 (left), or UbcH10+Ube2S (middle), or UbcH10 then Ube2S (right). **D.** P32-labeled geminin was ubiquitylated by the APC plus UbcH10, and then was tested for ubiquitylation by HUWE1 or HUWE1$^{UBDmu}$. Samples were resolved by SDS-PAGE and autoradiography.

A

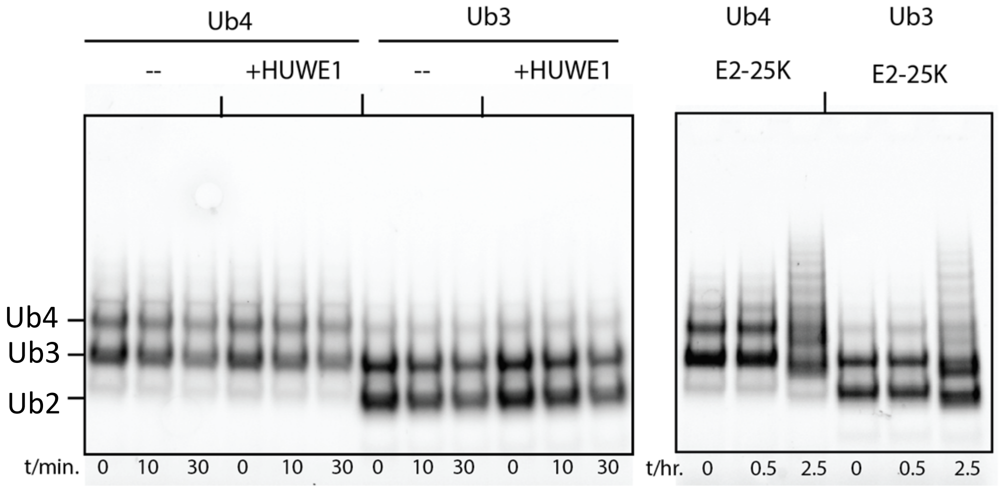


B

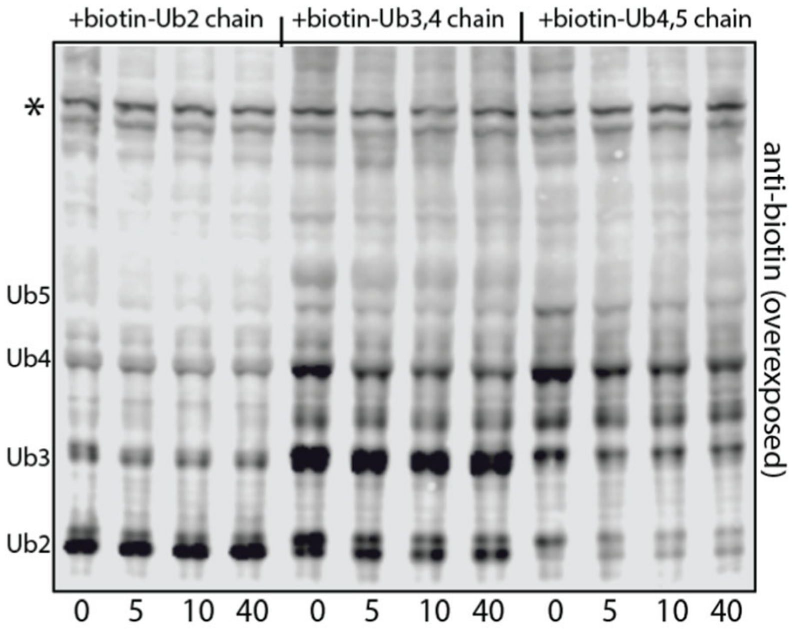


**Figure S8. HUWE1 does not react with free Ub chains. A.** K48-linked, Dy550-labeled Ub chains were synthesized and separated as described in methods. These chains were incubated with 30nM HUWE1 in a ubiquitylation reaction. Samples were separated by SDS electrophoresis and analyzed by a fluorescence imager (Bio-Rad). A standard Ub-chain formation reaction by the E2, E2-25K, involving these chains were used as a positive control (see methods). **B.** Preformed K48-linked $Ub^{L73F}$ chains were incubated with HeLa extract. Ub is also biotinylated at the N-terminus. The samples were analyzed by anti-biotin western blotting which was intentionally overexposed to reveal the further ubiquitylation (if any).

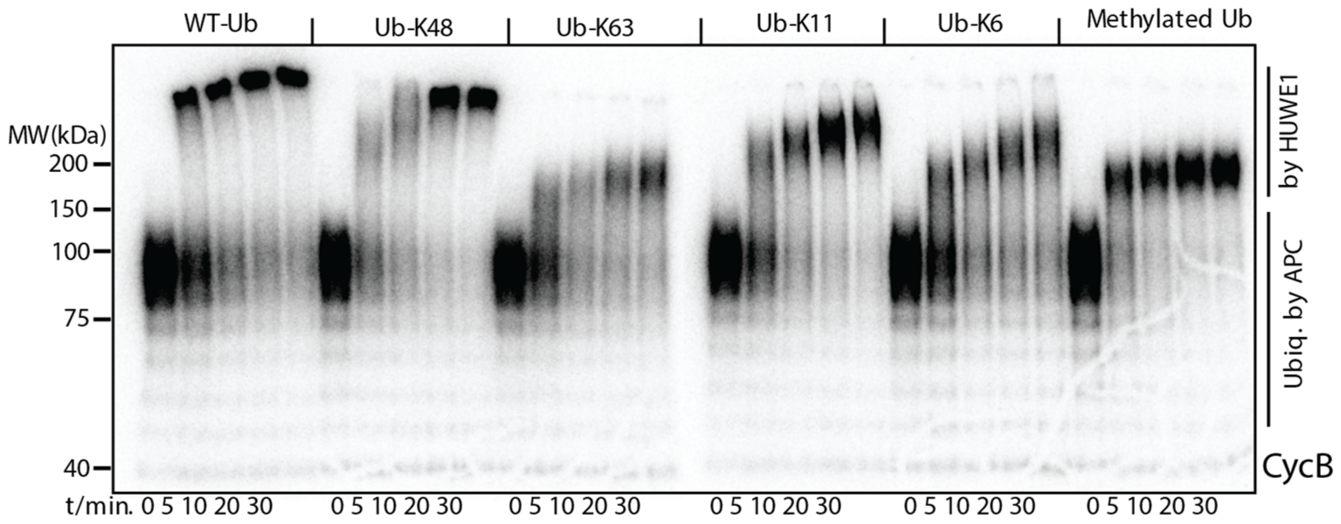


**Figure S9. HUWE1 reaction forms different Ub linkages on substrates.** CycB-cpGFP-polyUb ($p^{32}$ labeled) was ubiquitylated by rHUWE1 in the presence of indicated Ub variants. Samples were analyzed by autoradiography.

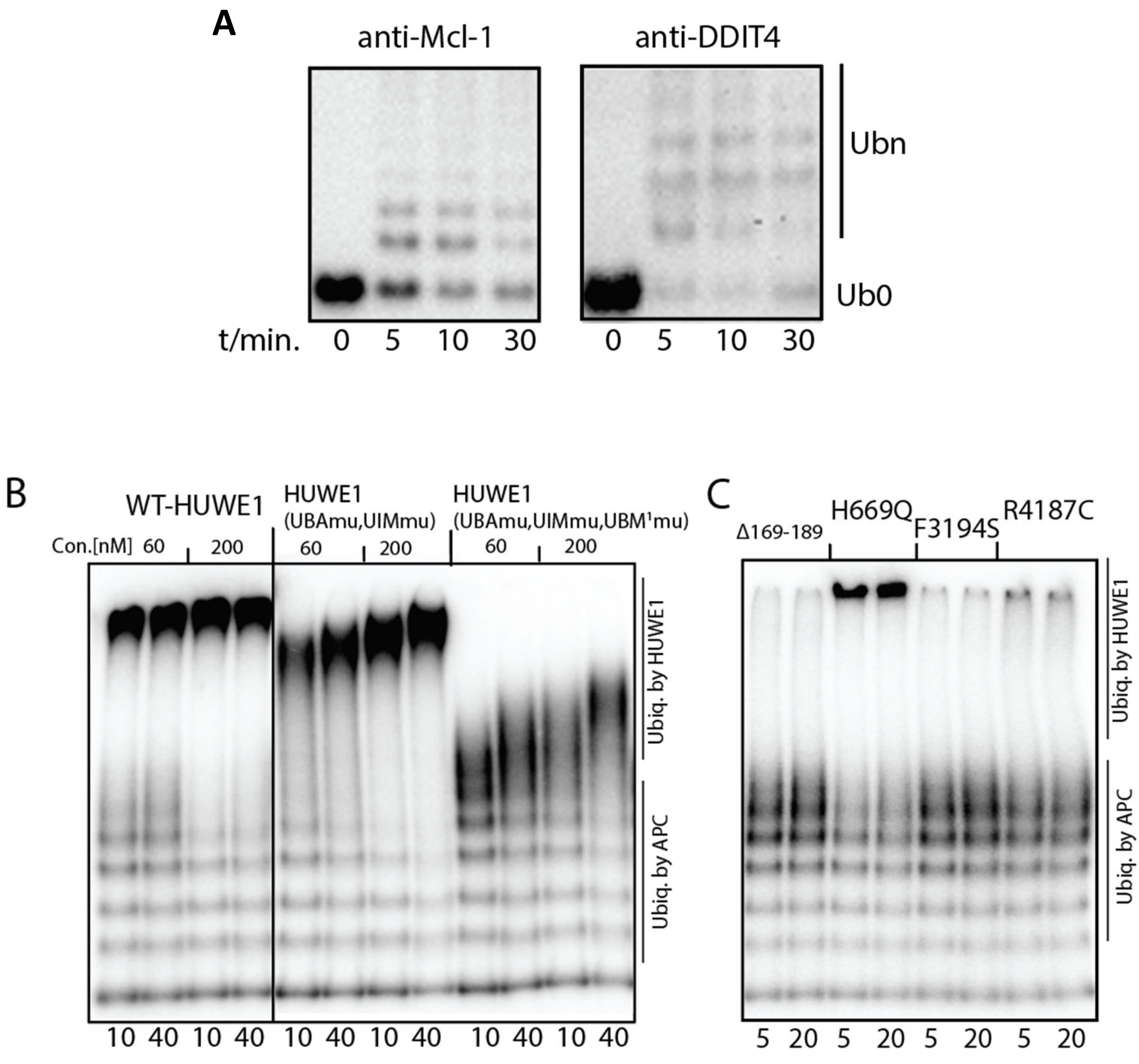


**Figure S10. The UBDs of HUWE1 mediate its UDL activity. A.** HUWE1$^{UBDmu}$ is competent for ubiquitylating MCL1-1 and DDIT4. Ubiquitylation of purified MCL-1 and DDIT4 by HUWE1$^{UBDmu}$. Samples were processed as in Figure 1D. **B.** WT HUWE1 or HUWE1 variants with indicated UBD mutations were tested for ubiquitylation of cycB-cpGFP-polyUb, at indicated concentrations. **C.** As in B, but using purified HUWE1 with indicated mutations (35).

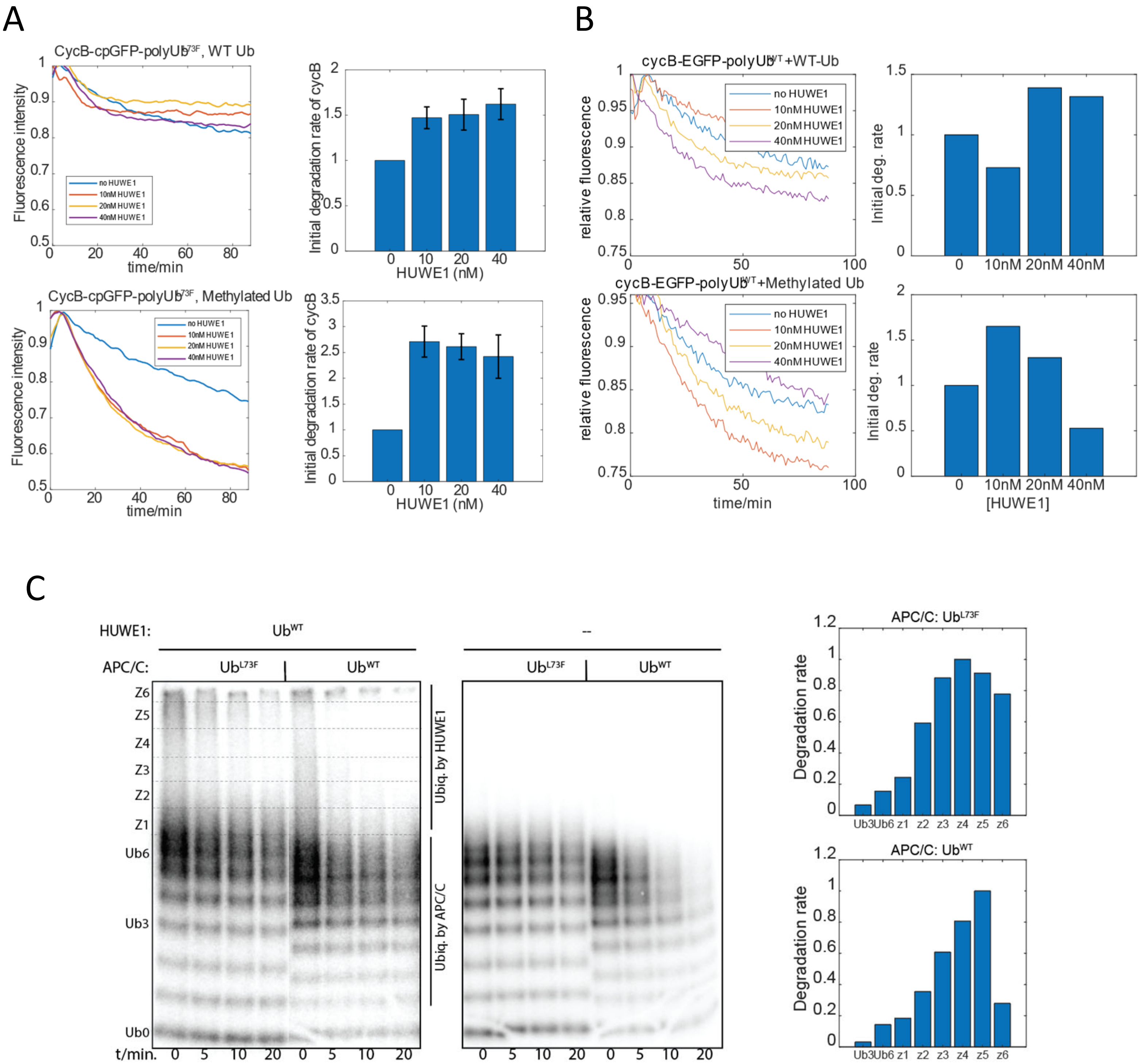


**Figure S11. UDL stimulates proteasomal degradation but inhibits it at very Ub stoichiometry. A.** GFP fluorescence intensity in a RUDS containing the 26S proteasome, HUWE1, cycB-cpGFP-polyUb$^{L73F}$(pre-ubiquitylated by APC-UbcH10) and either WT Ub or methylated Ub (see methods). The initial rate of fluorescence decay, normalized by the value in the absence of HUWE1, is presented on the right. Error bars represent the standard deviation of three replicates. **B.** As in A, but using cycB-EGFP-polyUb$^{WT}$ as the substrate in a RUDS containing either WT Ub or methylated Ub. **C.** P$^{32}$-labeled cycB-cpGFP was reacted with the APC in the presence of Ub$^{WT}$ or Ub$^{L73F}$ to form polyubiquitylation. The products were then briefly reacted with HUWE1 in the presence of Ub$^{WT}$ or no Ub, and were quenched with an E1 inhibitor before the 26S proteasome was mixed in (t=0). The degradation rate of each ubiquitylated cycB species, separated into six zones (z1~z6), was determined from a time series and presented on the right.

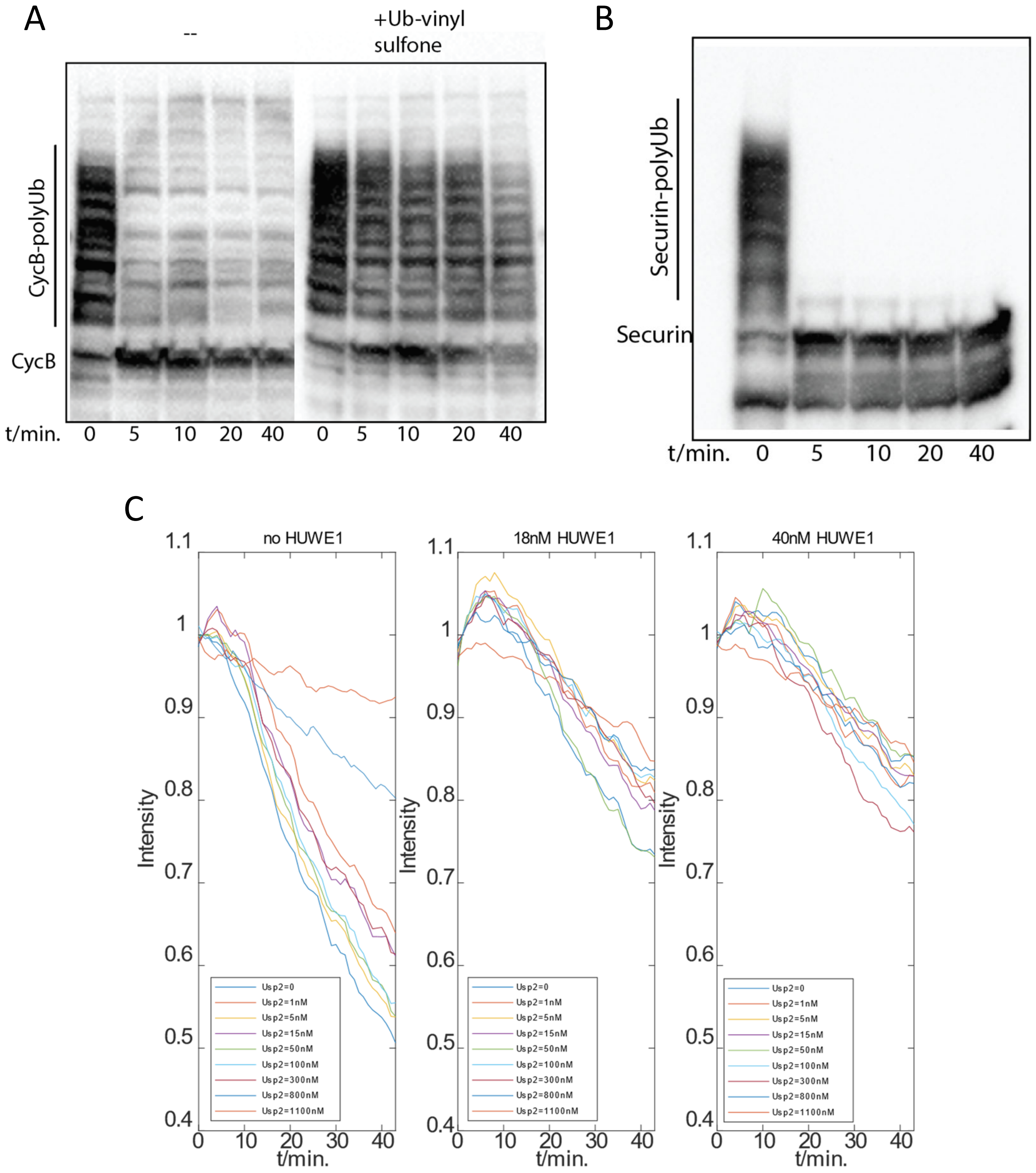


**Figure S12. Cell extract contains high deubiquitylating activity. A,B.** Deubiquitylation of HA-cycB-cpGFP-polyUb$^{WT}$ and securin-mcherry-polyUb$^{WT}$ in HeLa S3 extract. Substrates were first ubiquitylated by purified APC plus UbcH10 and then were incubated with the cell extract which had been treated with a proteasome inhibitor (Bortezomib) and an E1 inhibitor (MLN7243). 25uM DUB inhibitor Ub-vinyl sulfone was added as a control. Samples were analyzed by anti-HA and anti-securin western blotting. **C.** Raw data for figure 5A.

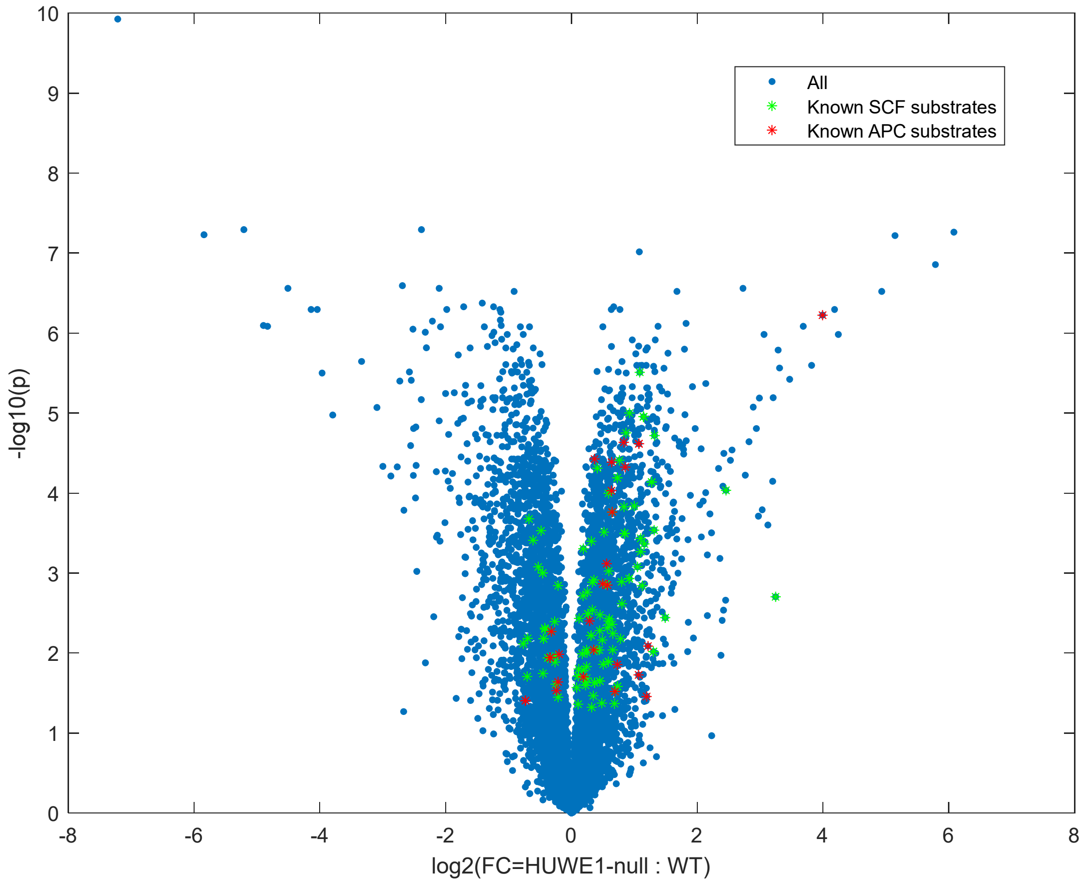


**Figure S13. Volcano plot of the fold change (FC) and p values of proteins in HUWE1Δ vs. WT cells.** The whole cell lysate of HUWE1Δ and WT HEK293 cells was analyzed by quantitative mass spectrometry (Fig. 4C). The FC (X) of identified proteins was plotted with the p values (Y). Known APC and SCF substrates *(75)* were highlighted with green and red asterisks.

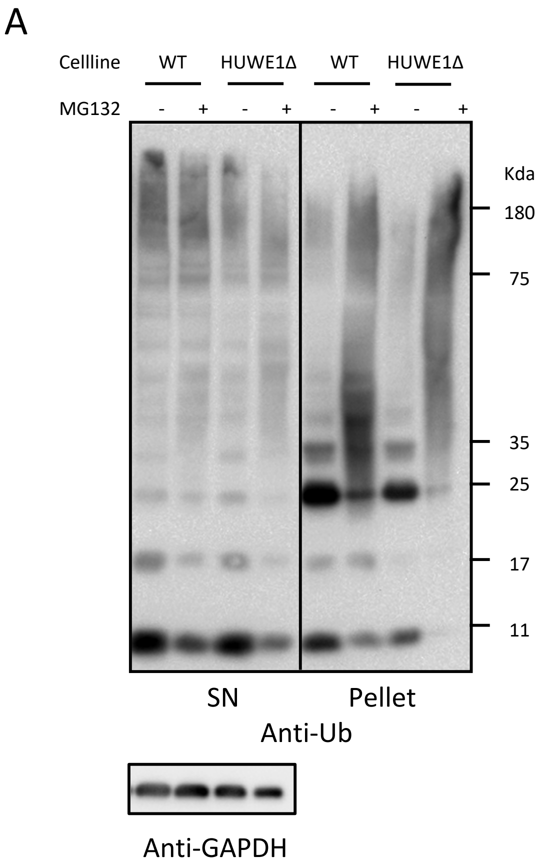


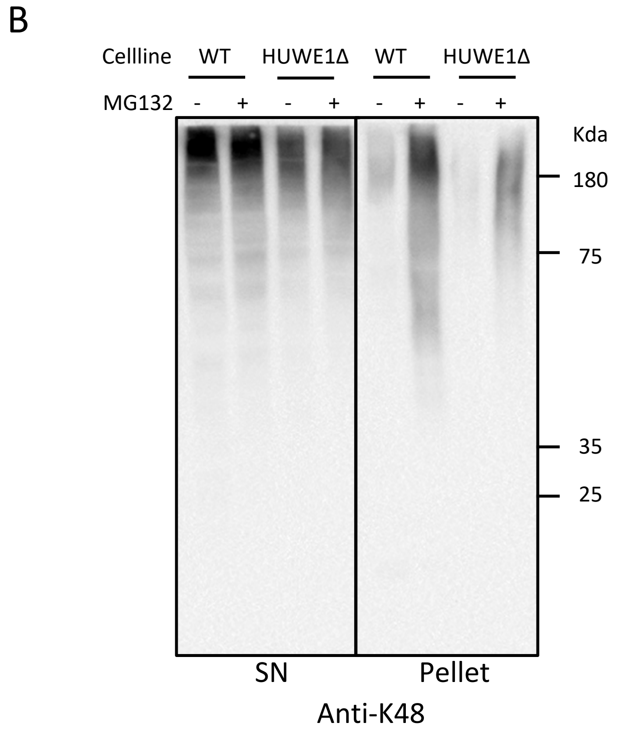


**Figure S14. HUWE1 knockout decreased the accumulation of K48-chain conjugates by proteasome inhibition, without affecting the overall ubiquitylation level. A.** WT HEK293 or HUWE1Δ cells were treated with MG132 or DMSO for 8hrs. The cells were lysed, and the soluble fraction(SN) and the pellet were separated by centrifugation and were probed by anti-Ub (**A**) and anti-K48 chain (**B**) western blotting. GAPDH was used as a loading control.

### Significantly up-regulated proteins in HUWE1Δ

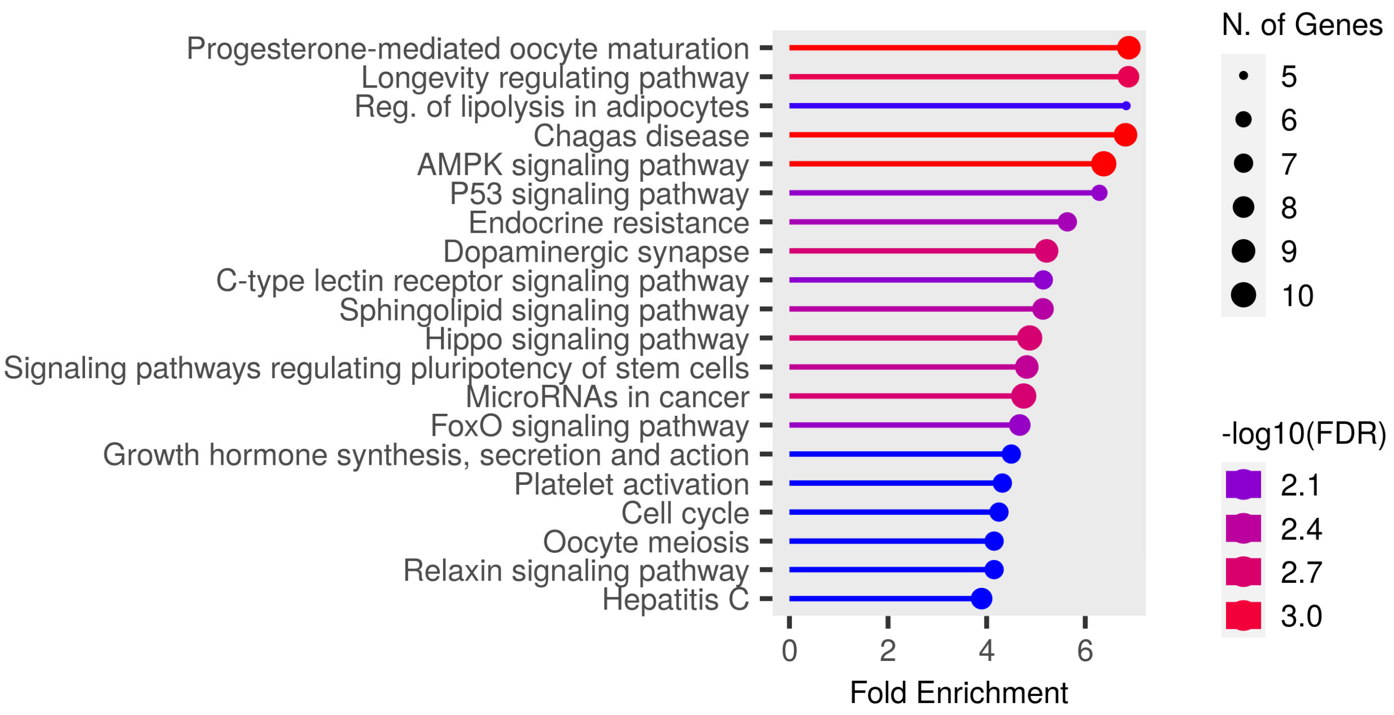


### Significantly down-regulated proteins in HUWE1Δ

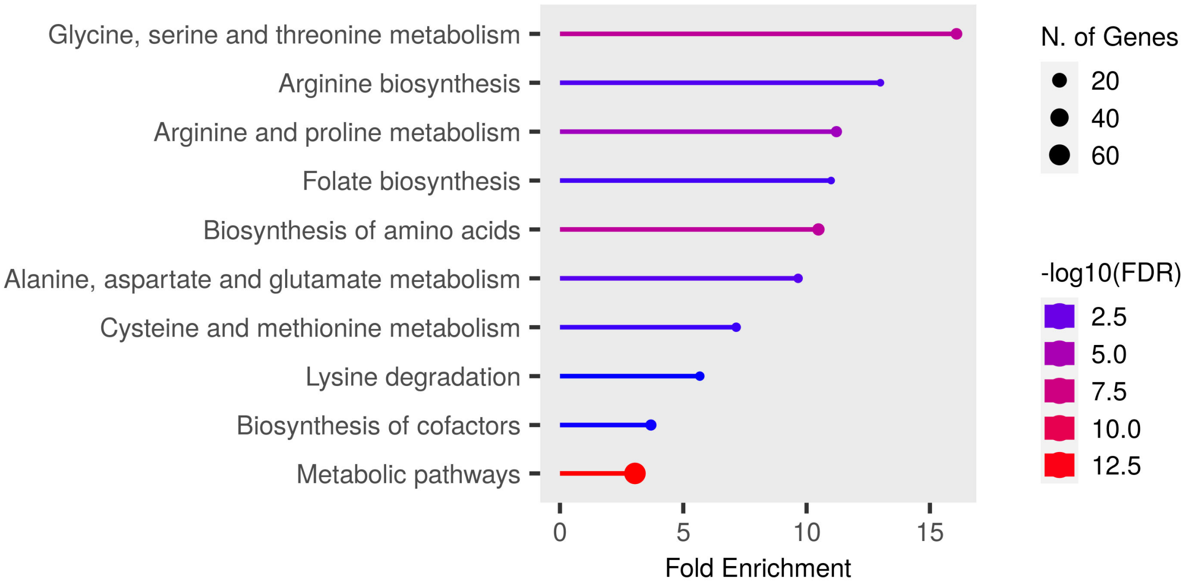


**Figure S15. Gene Ontology(GO) analysis of significantly up or down regulated proteins in HUWE1Δ cells.** Protein abundance in WT and HUWE1Δ HEK293 cells were compared using label-free quantitative mass spectrometry (methods). Proteins that are significantly ($P<0.05$) up or down regulated in HUWE1Δ (log2(fold change) >1 or < -1) are identified and analyzed by shinyGO (*44*). The gene names associated with each GO term can be found in Table S2.

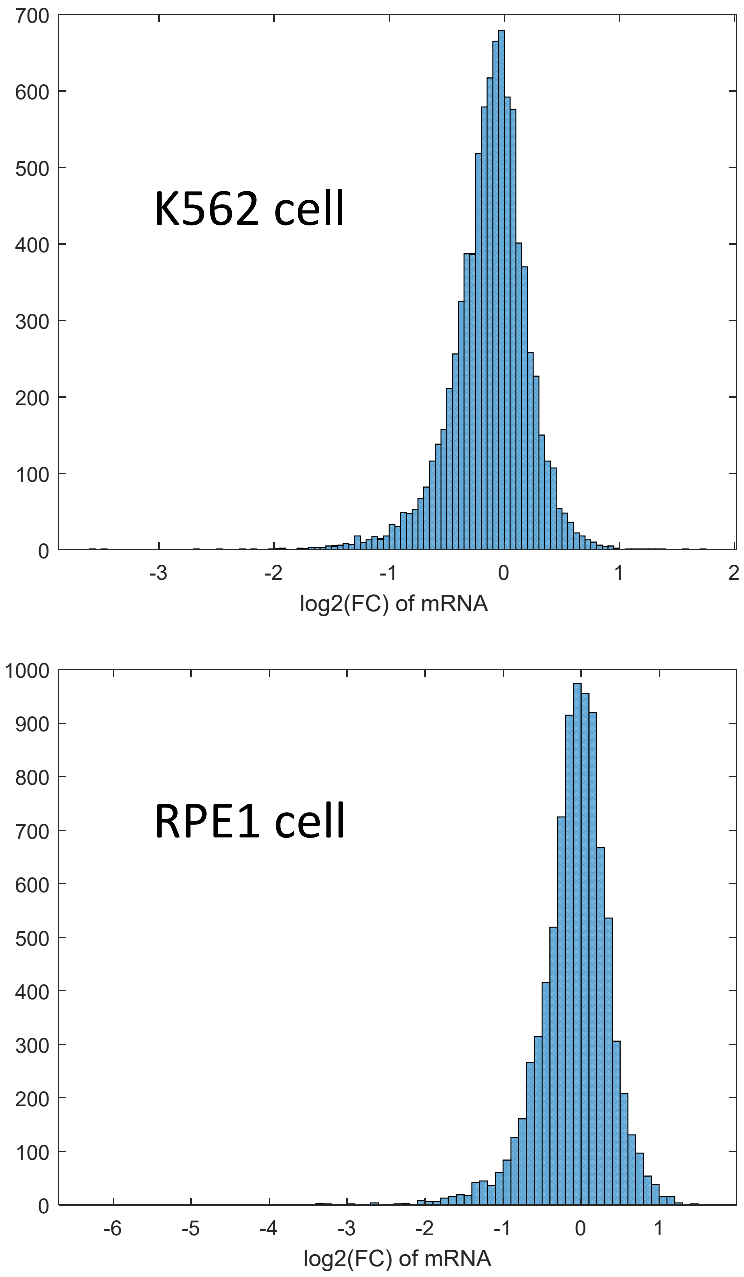


**Figure S16. HUWE1 knockout does not significantly affect most transcripts' levels.** The fold changes of ~8600 transcripts were extracted from a dataset generated by Perturb-seq *(45)* in either K562 and RPE1 cells, and the distribution is plotted. There are 11 and 29 transcripts whose levels increase by more than 2X in K562 and RPE1 cells respectively.

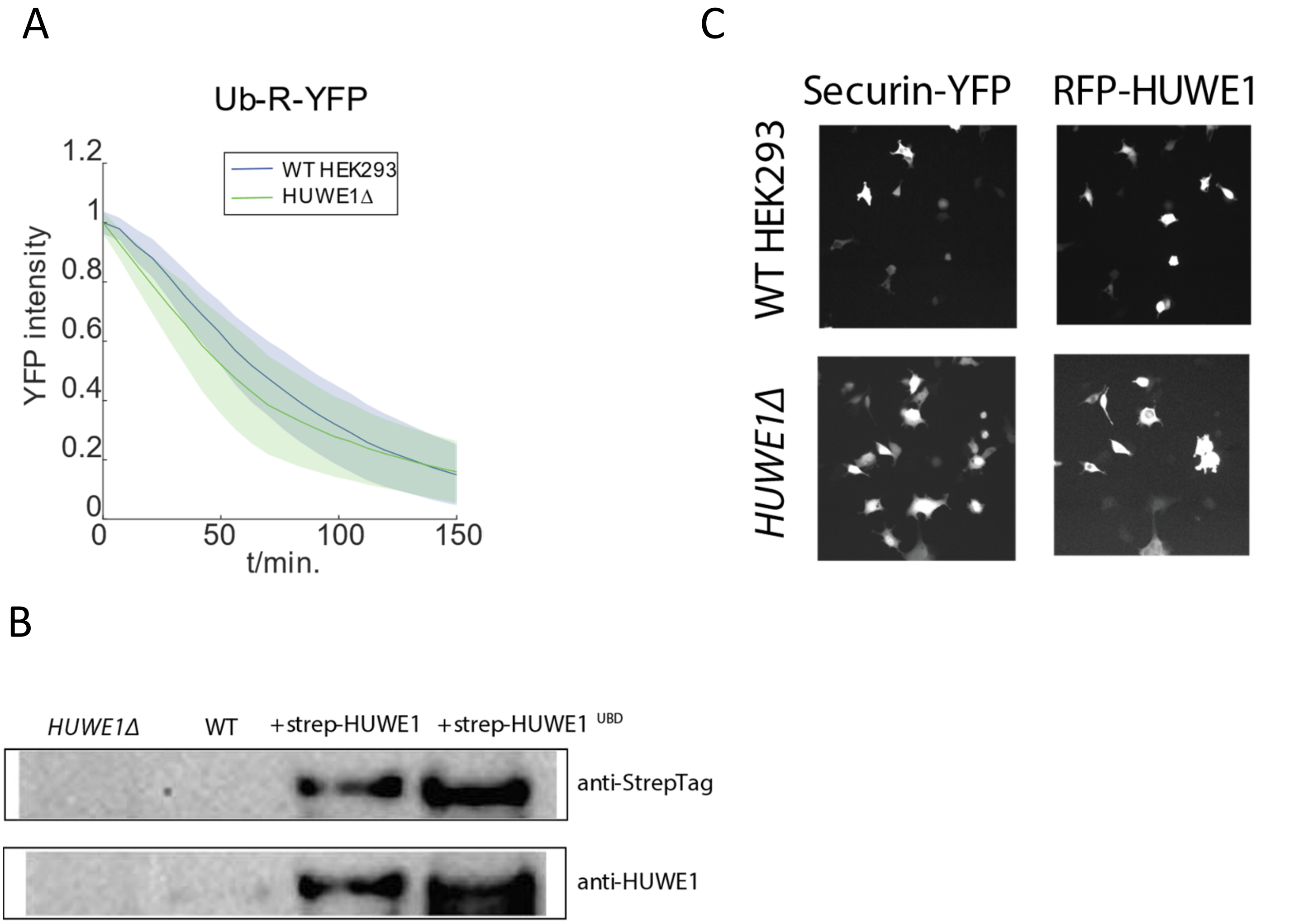


**Figure S17. Expression of RFP-HUWE1. A.** Ub-R-YFP was transiently expressed in WT HEK293T or *HUWE1Δ.* Cycloheximide was added at t=0 to stop protein synthesis, and the cells were imaged once every 7 minutes. The averaged degradation kinetics of these reporters were calculated from single-cell traces (Ub-R-YFP N=31). The shaped area represents the standard deviation. **B.** Anti-HUWE1 and Anti-streptag western blotting of cells expressing strepTag-RFP-HUWE1 or strepTag-RFP-HUWE1$^{UBDmu}$. **C.** Sample images of both securin-YFP and RFP-HUWE1 channels for WT or *HUWE1Δ* transiently transfected with these constructs.

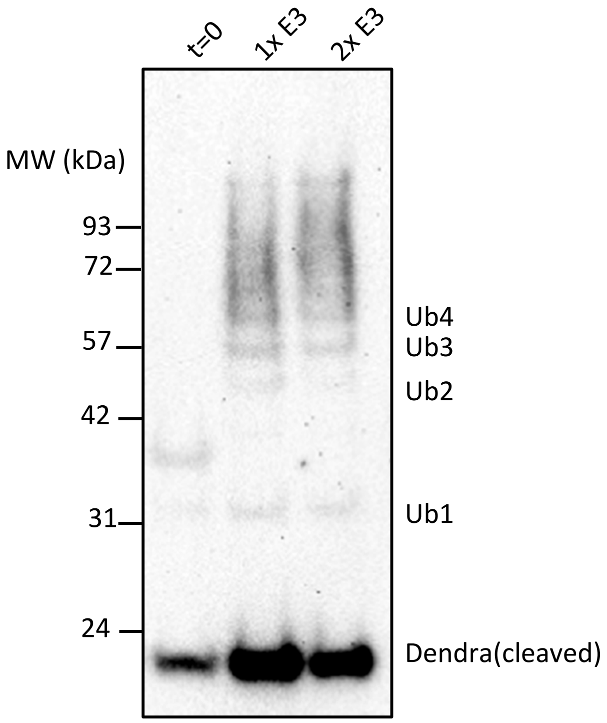


**Figure S18. Ubiquitylation of dendra substrate.** Recombinant SUMO-dendra2 was purified and photo-cleaved as in a previous study(*47*). After de-sumoylation, the substrate was incubated with 1x or 2x Ubr1 (E3), and was analyzed by anti-dendra western blotting.

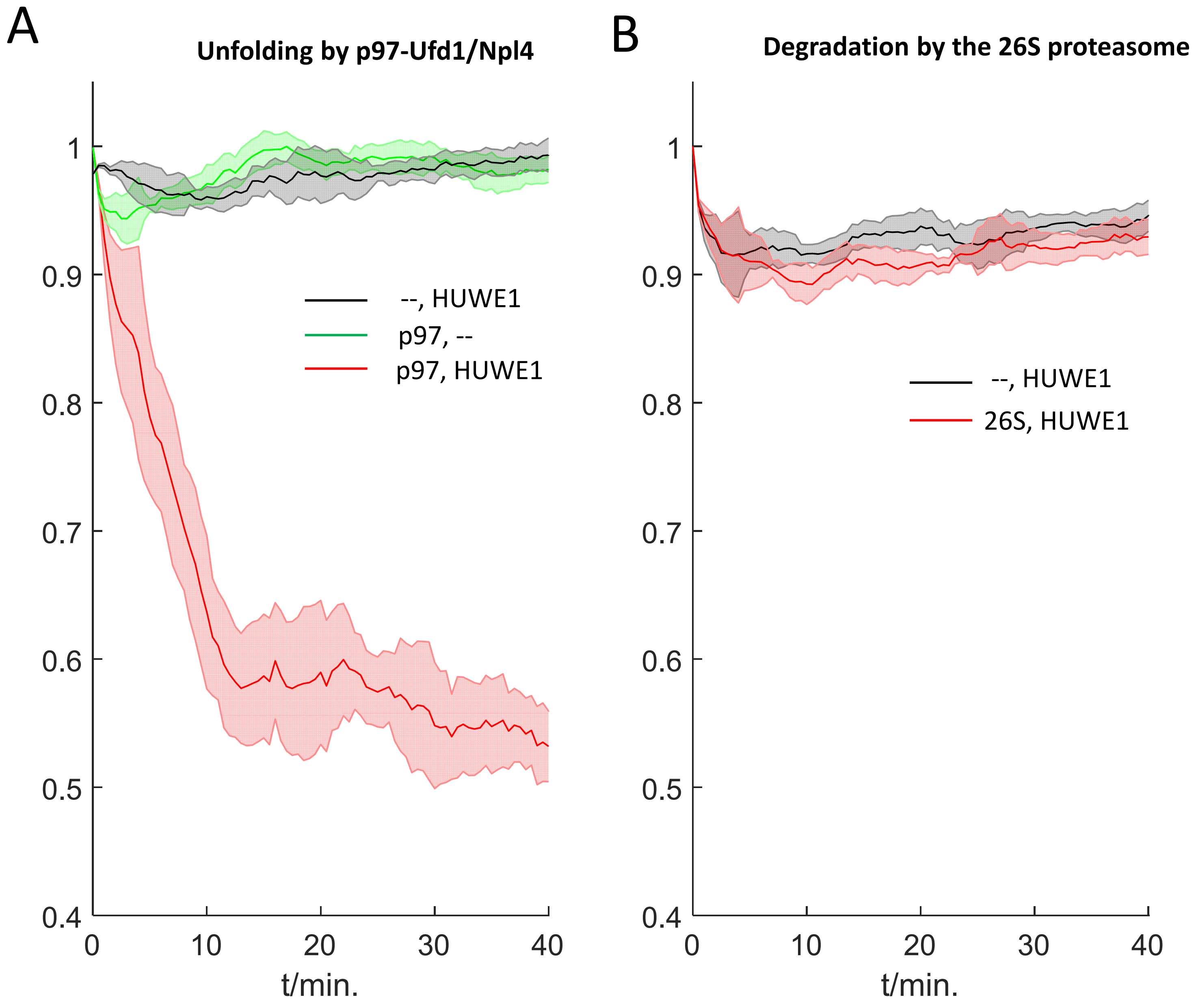


**Figure S19. UDL stimulates p97-mediated unfolding of a tetramer substrate. A.** HA-tagged synthetic substrate Ub(3)-VASP-cpGFP was ubiquitylated by rHUWE1 as in Figure 3A, and then was incubated with recombinant p97-UFD1-NPL4 complex in the presence of 1uM GroEL(D87K). The fluorescence intensity was monitored using a plate reader. **B.** the same substrate, after HUWE1 reaction, was incubated with purified human 26S proteasome.

A

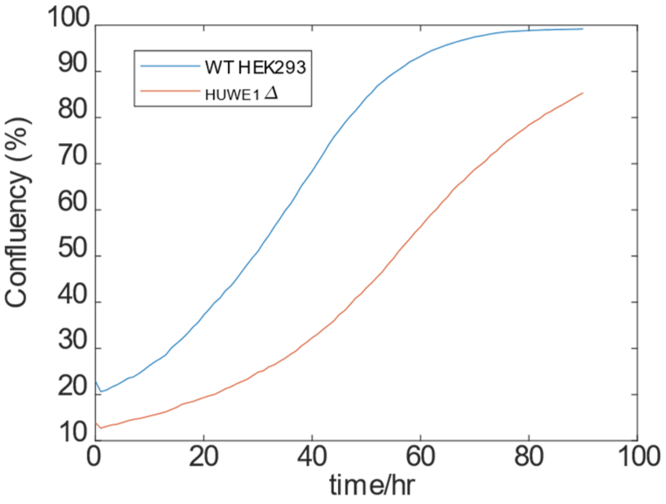


B

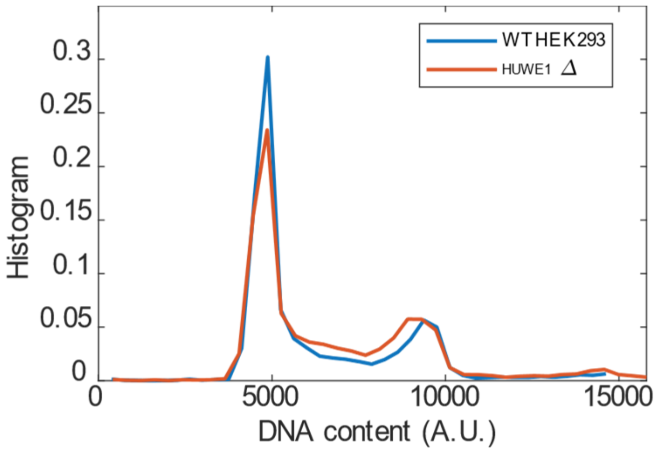


| | WT HEK293 | HUWE1Δ |
|---|---|---|
| G1 | 60% | 56% |
| S | 20% | 22% |
| G2/M | 20% | 22% |

**Figure S20. Characterization of HUWE1Δ HEK293 cells. A.** Growth curve of HUWE1Δ cells, determined by an Incucyte imager. **B.** DNA content of WT and HUWE1Δ, determined by FACS and Propidium Iodide staining.

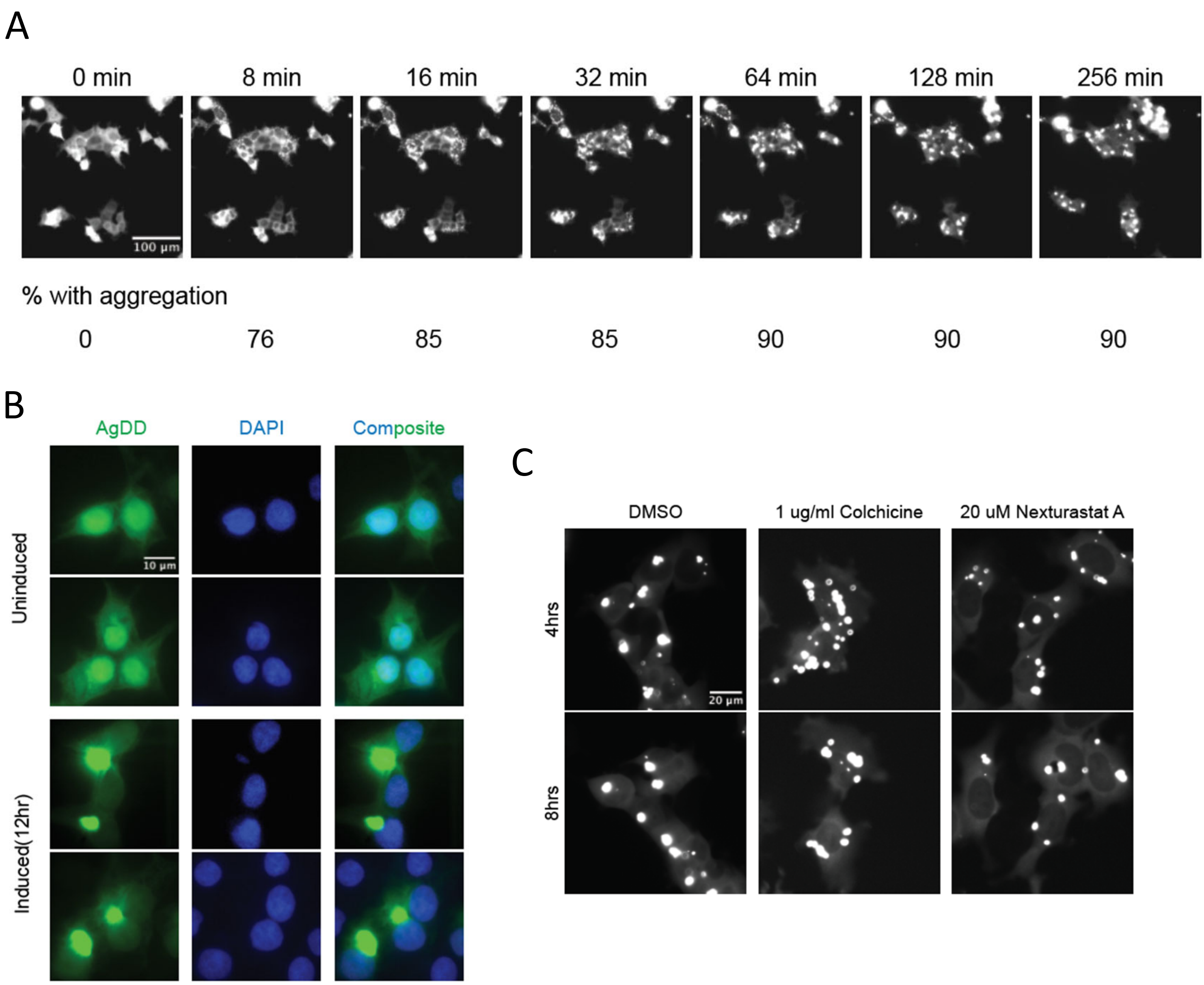


**Figure S21. AgDD forms an aggresome-like aggregates upon induction. A.** Kymograph images of agDD(GFP)-expressing HEK293 cells. Shield-1 was removed at t=0 to induce aggregation. **B.** Fluorescence image of fixed HEK293 cells containing agDD aggregates. **C.** Cells were treated as in A, but with colchicine or Nexturastat A (a HDAC6 inhibitor) added right after shield-1 removal. Images were taken 4 or 8hrs after the induction.

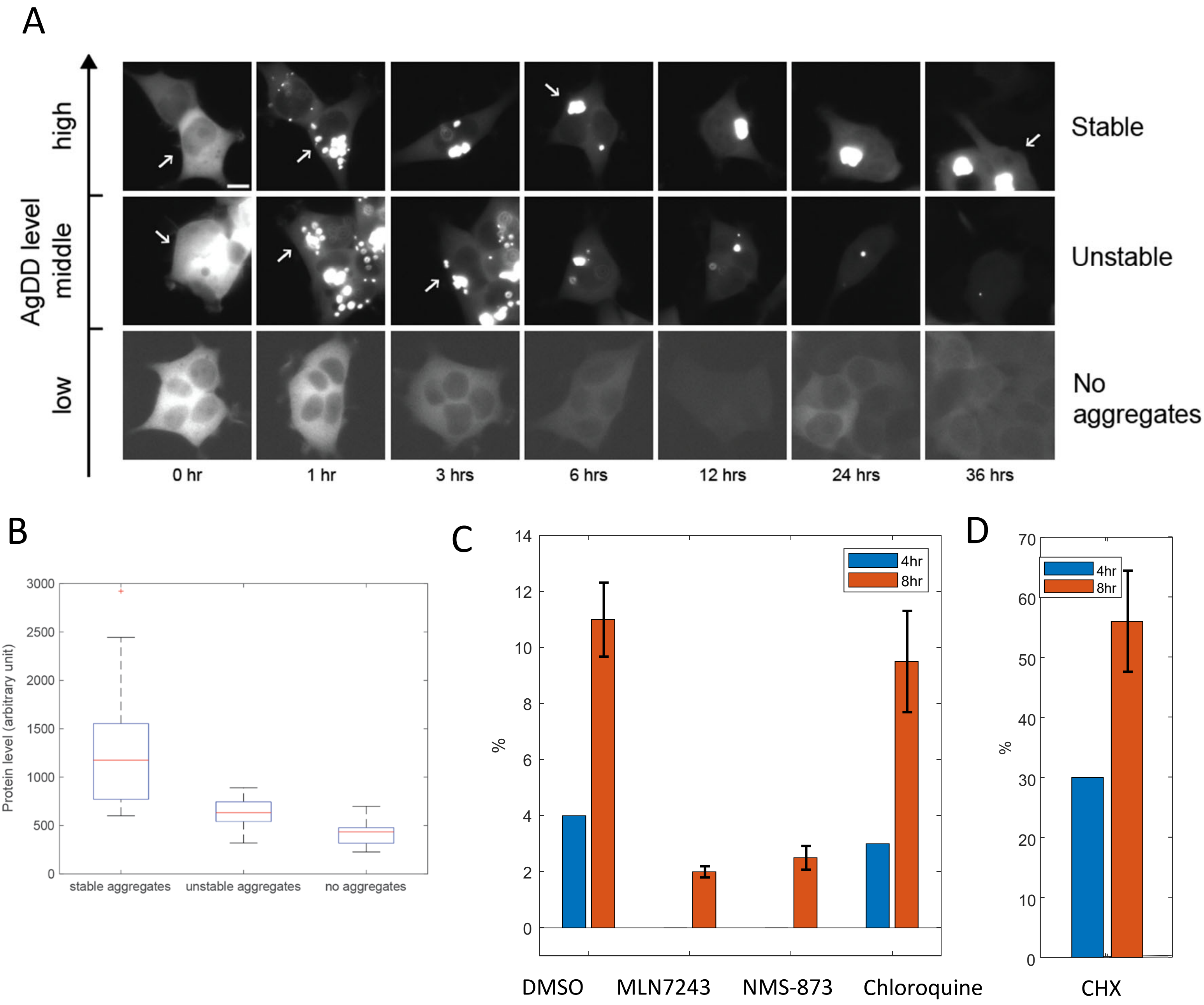


**Figure S22. Clearance of agDD aggregates in cells. A.** Example images of agDD-expressing cells exhibiting three different phenotypes: no aggregates, unstable aggregates that undergo spontaneous clearance, stable aggregates without obvious clearance. Shield-1 was removed at t=0 to induce aggregation. **B.** AgDD levels, as determined by the total fluorescence intensity in individual cells exhibiting the three phenotypes. **C.** The percentage of cells that have spontaneously cleared agDD aggregates at 4 or 8hrs after shield-1 removal, in the presence of indicated compounds. MLN7243: Ub E1 inhibitor; NMS-873: p97/VCP inhibitor; Chloroquine: autophagy inhibitor. **D.** As in the C, but cycloheximide (CHX) was added 30 minutes after shield-1 removal.

A

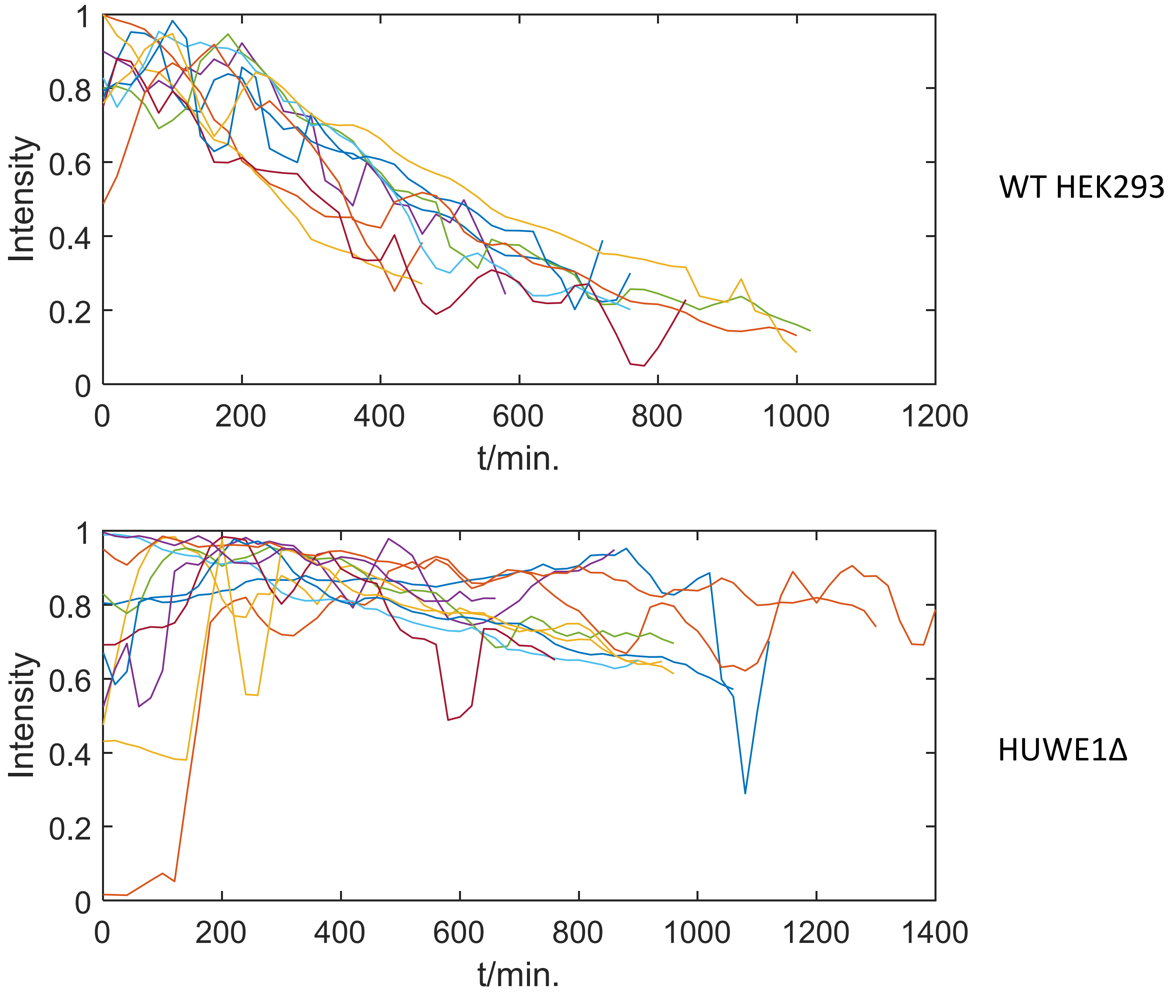


**Figure S23. Raw data for figure 7B.**

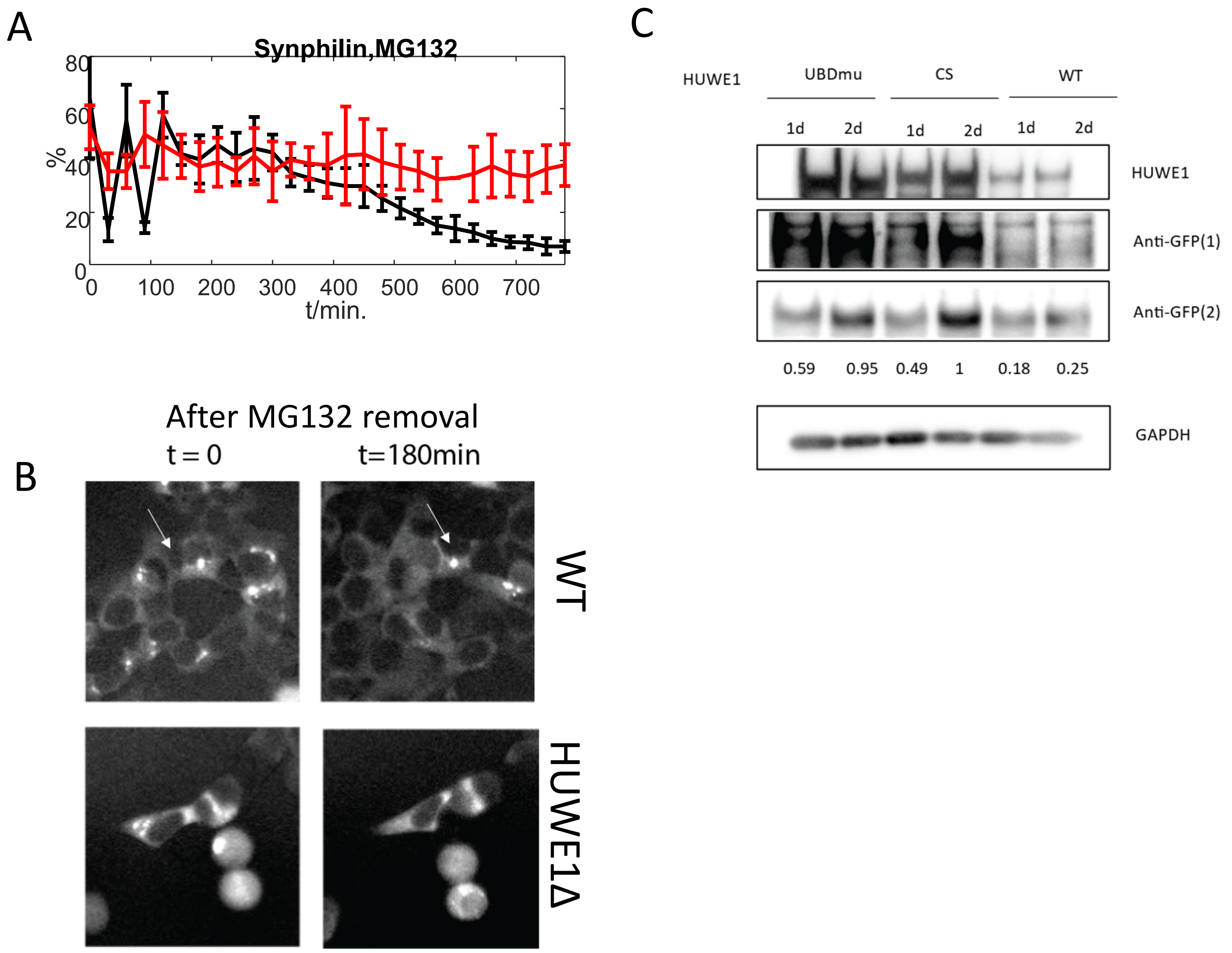


**Figure S24. HUWE1 mediates the clearance of synphilin aggresome and promotes the degradation of over-expressed synphilin. A.** Synphilin aggresomes were induced in WT or HUWE1Δ cells stably expressing synphilin-GFP by incubating the cells with MG132 for 8hrs (*63*). MG132 was washed off at t=0 to initiate the clearance of synphilin aggresome. The clearance process was monitored by timelapse microscopy (methods). Error bars represent the standard deviation of three replicates. **B.** Sample images. **C.** HUWE1 (WT, $HUWE1^{CS}$ or $HUWE1^{UBDmu}$) and synphilin-GFP plasmids were co-transfected into HEK293 cells. 1d or 2d after transfection, cells were harvested and the whole-cell lysate was analyzed by western blotting. Two anti-GFP antibodies were used to quantify the synphilin-GFP levels. Numbers are quantification of the relative synphilin concentrations (an average of the two anti-GFP blots).

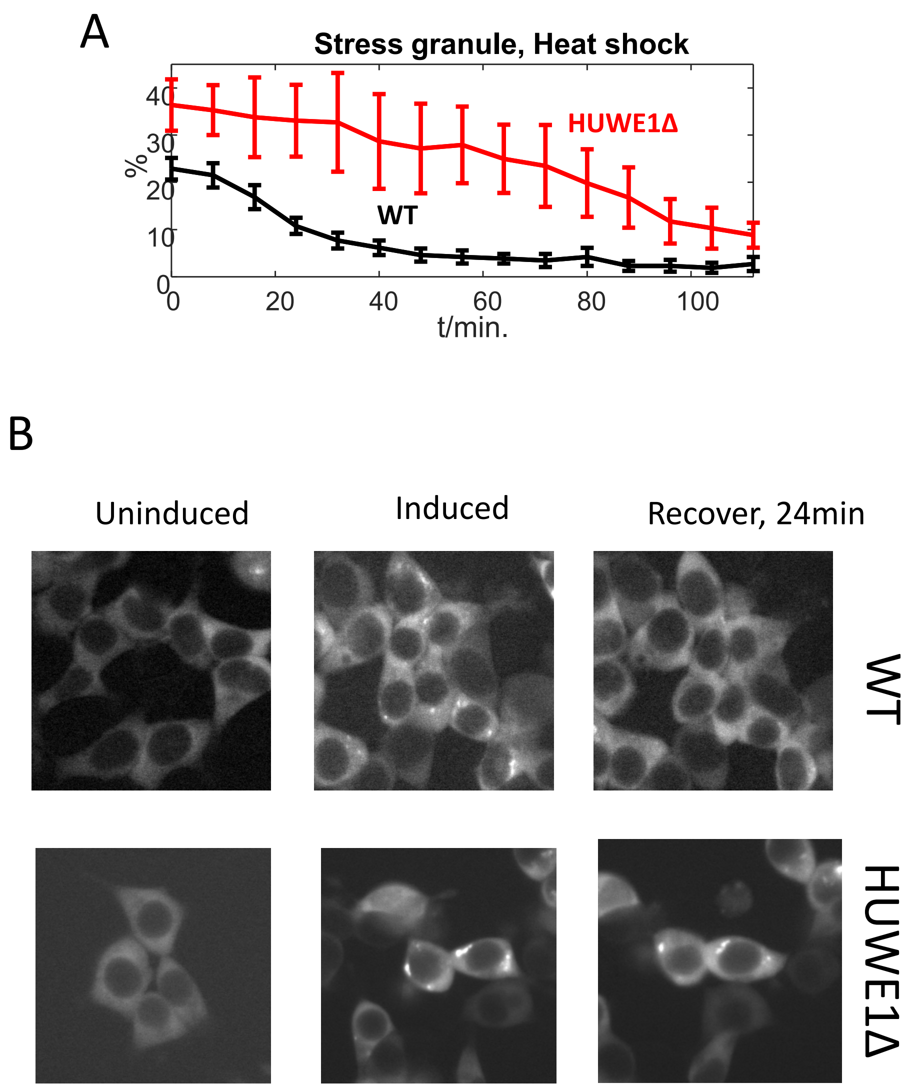


**Figure S25. Clearance of heat-shocked induced stress granules in WT or HUWE1Δ.** Experiment and analysis was performed as described in Figure 7E,F, except that stress granules were induced by heat shock at 43 degree for 60 minutes, and cells were moved to 37 degree for recovery. Error bars represent the standard deviation of three replicates.

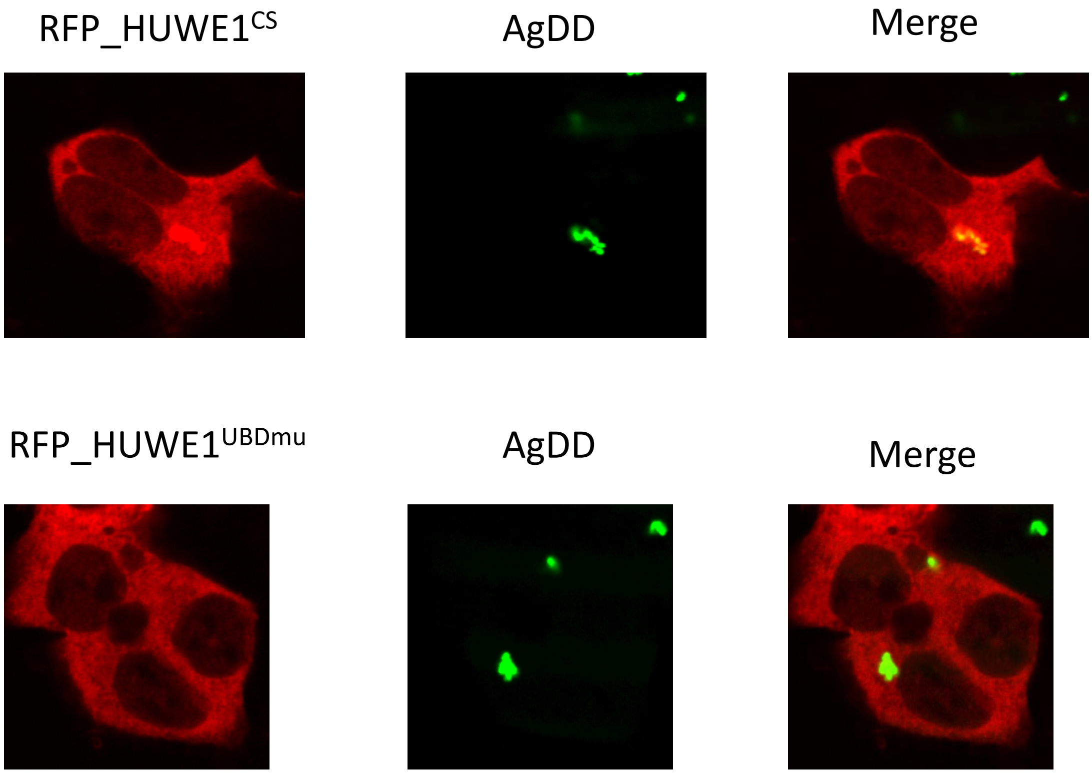


**Figure S26. Colocalization of RFP-HUWE1 with agDD aggregates requires its Ub-binding domains.** Experiment was performed as described in Figure 7G, but using either RFP-HUWE1$^{CS}$ or RFP-HUWE1$^{UBDmu}$.

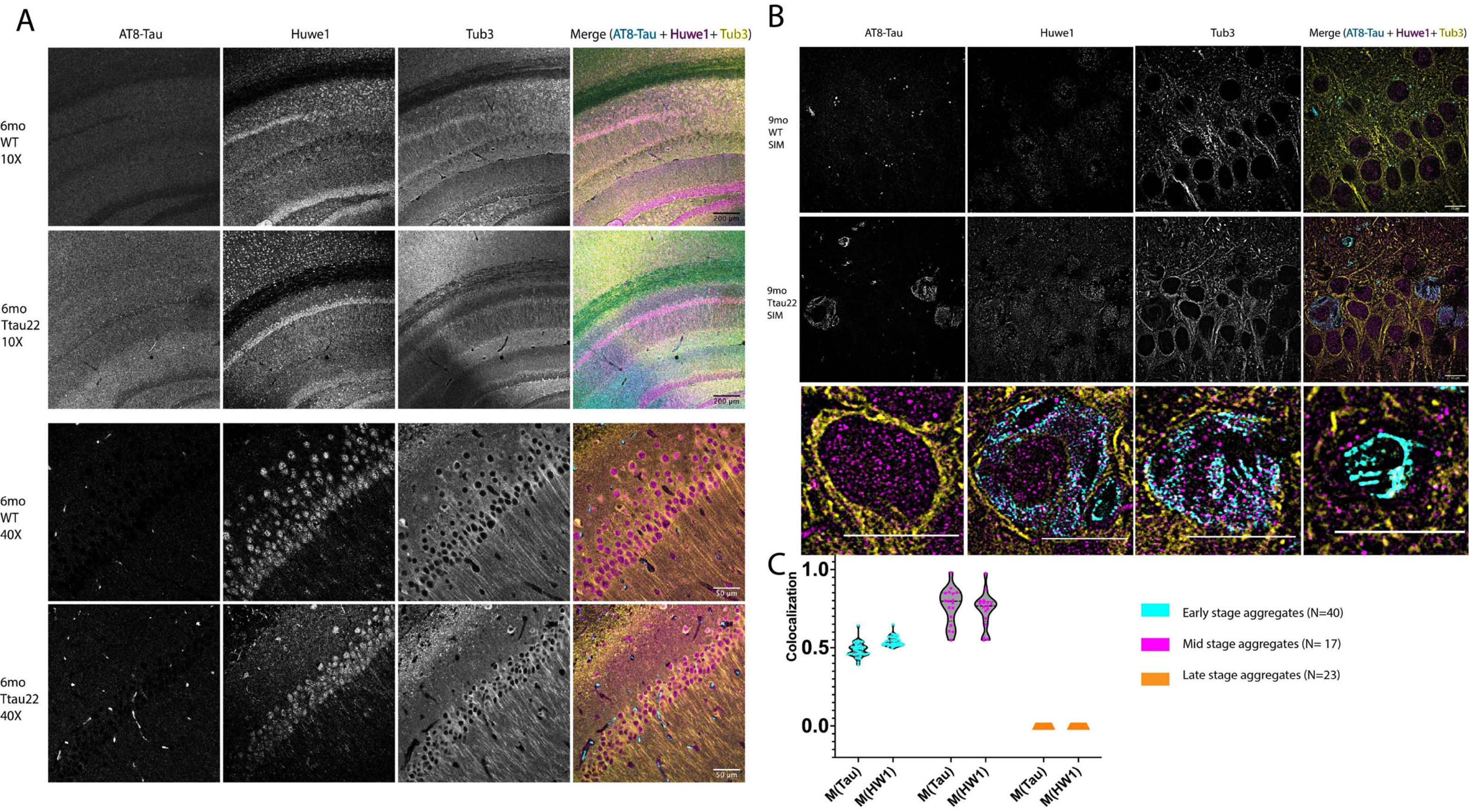


**Figure S27. Expression and distribution of HUWE1 in WT and THY-Tau22 mice. A.** Confocal images showed that HUWE1 was highly expressed in cortical neurons in both WT mice and THY-Tau22 mutant mice. **B.** Structured-Illumination-Microscopy (SIM) demonstrated neuronal colocalization of HUWE1 and Tau in the symptomatic THY-Tau22 mice. **C.** Quantification of colocalization using Manders' overlap coefficient showed that ~50% Tau is associated with HUWE1 in early-stage tau aggregates, ~ 75% in mid-stage tau aggregates, and 0% in late-stage tau aggregates.

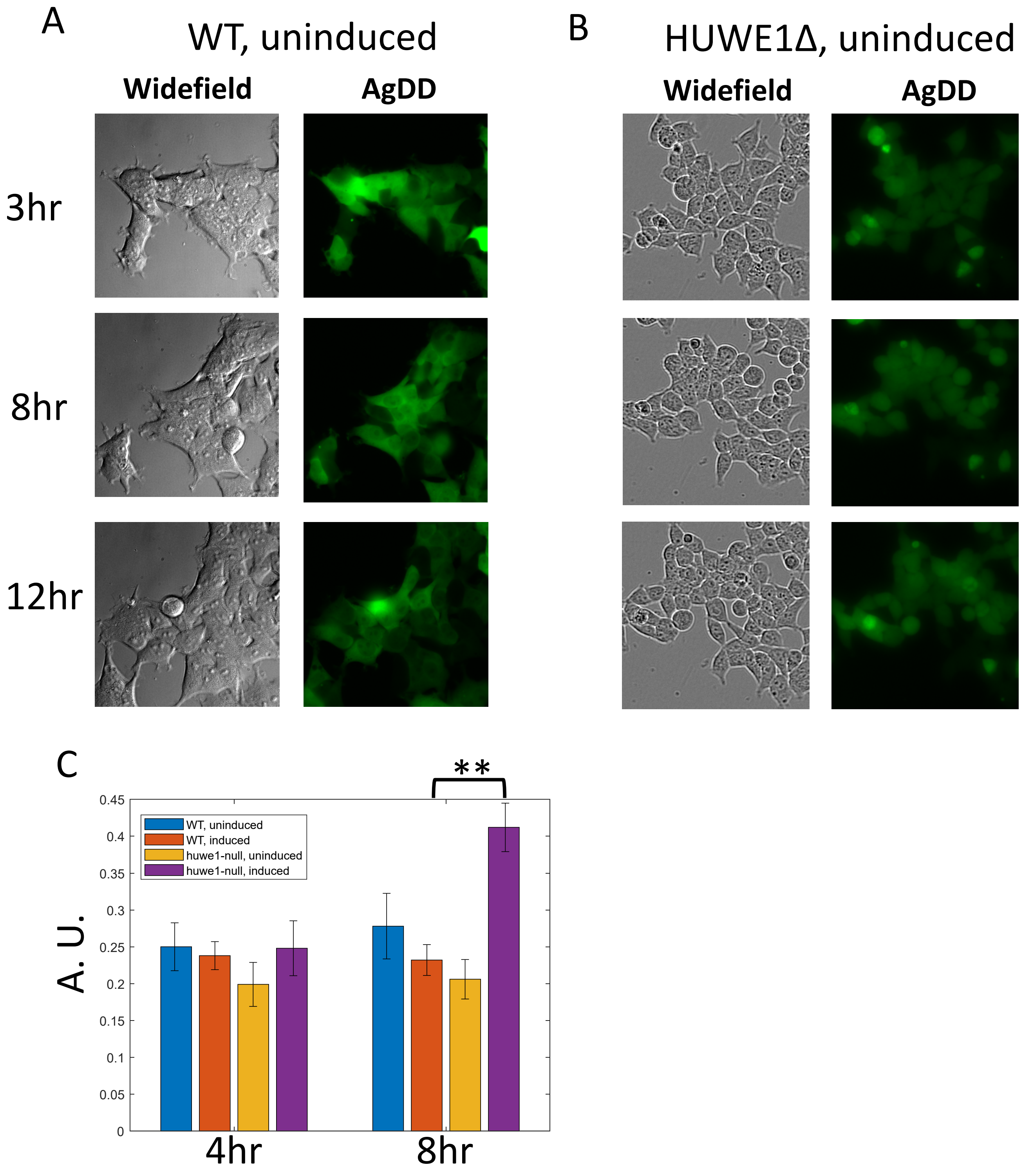


**Figure S28. HUWE1 controls the cytotoxicity of agDD aggregates (controls). A, B.** as described in Figure 8A,B, but without inducing agDD aggregates. **C.** As described in Figure 8A,B. The viability of the cells were determined using CytoTox-Glo(Promega).

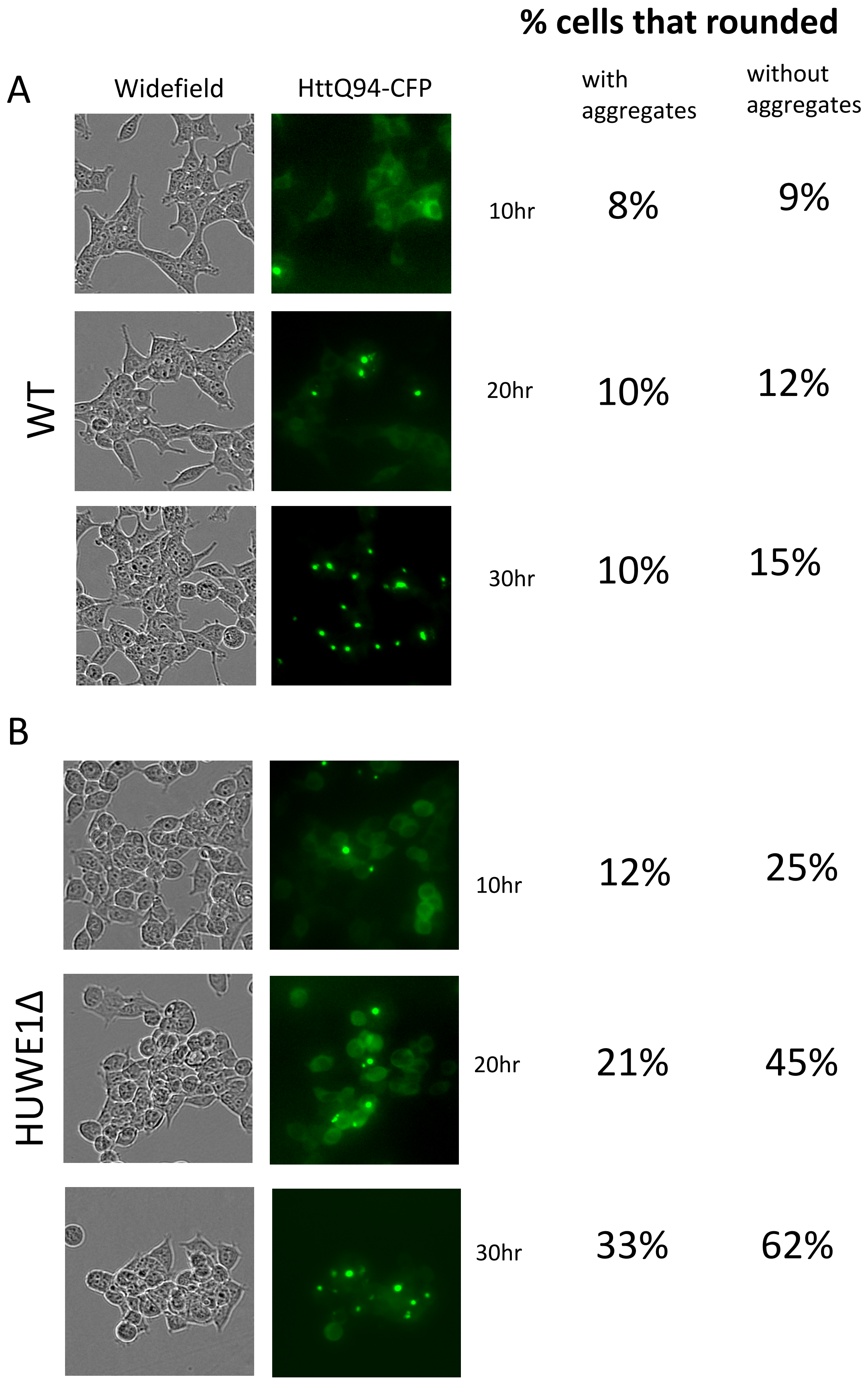


**Figure S29. HUWE1 controls the cytotoxicity of Htt(Q94) aggregates.** Sample images of the experiment described in figure 8D. The percentage of cells that have rounded at indicate time points, in the population with or without a clear Htt(Q94) aggregate (instead showing diffusive Htt-CFP signal), was counted.
